# CellBench: task-dependent performance and generalization limits of single-cell foundation models

**DOI:** 10.64898/2026.04.01.714123

**Authors:** Yongjie Xu, Yiyun Li, Yue Yuan, Qingtao Xia, Hangyao Tu, Fengjun Hu, Zelin Zang

**Author notes:** These authors contributed equally (Yongjie Xu, Yiyun Li, Yue Yuan).

## Abstract

Single-cell foundation models (scFMs) are increasingly used for transcriptomic analysis, yet it remains unclear what biological information their embeddings preserve and when they outper- form dataset-fitted methods. Here, we present CellBench, a unified benchmark of frozen scFM representations across tissues, species, and genetic perturbation screens. We find that different models capture distinct aspects of cellular biology, while classical baselines remain competitive across tasks. Human-pretrained representations transfer to supervised mouse analyses, but under a shared decoder they fail to improve genome-wide absolute error over predicting no change for unseen perturbation targets. Neither model size nor reported pretraining data volume reliably predicts performance. Excluding cohorts with sample or donor identifier matches to pretraining records substantially reduces annotation advantages without changing model rankings appreciably. Together, these findings show that the value of scFM representations depends on the biological task and the strength of the evidence supporting their performance.

**HIGHLIGHTS:**

- Single-cell foundation models preserve distinct biological information across tasks
- Human-to-mouse transfer does not ensure unseen perturbation prediction
- Model size and pretraining corpus size do not reliably predict performance
- Excluding identifier-matched cohorts preserves rankings but reduces annotation gains

**THE BIGGER PICTURE:** Foundation models are becoming common tools for analyzing single-cell data, but choosing the right model remains challenging. Researchers often rely on benchmark rankings or model size to guide their decisions, even though these measures may not reflect the biological information a model preserves. CellBench shows that different models capture different aspects of cellular biology and that simpler methods can perform just as well on some tasks. It also reveals an important distinction between representing observed cellular states and predicting how cells respond to genetic interventions. Human-pretrained representations can transfer to mouse tissues, but their success in observational tasks does not ensure accurate prediction of unseen perturbations. Moreover, stable model rankings can hide small or dataset-dependent advantages, especially when pretraining data are not fully documented. These findings highlight the need to evaluate foundation models according to specific biological questions rather than overall rankings. Strong classical baselines, appropriate controls, and transparent pretraining records are essential for determining when these models provide reliable benefits. Ultimately, better evaluation can guide the development of models that are not only more powerful but also more useful for biological research.

**IN BRIEF:** CellBench compares 13 frozen single-cell foundation models with classical baselines fitted on training data across 12 human and mouse cohorts and four genetic perturbation screens. Different models capture different biological information. Human-pretrained models transfer well to mouse tissues, yet with a shared decoder none beats predicting no change on unseen perturbations. Model size and training data volume poorly predict performance, and removing human studies overlapping pretraining data erases the human annotation gain over baselines.

## INTRODUCTION

Large-scale single-cell RNA-sequencing atlases have provided extensive data from human or-gans, diseases, and model organisms^1–3^. These resources have supported the development of more than a dozen single-cell foundation models (scFMs)^4–18^. Some models have been pretrained on tens to hundreds of millions of cell profiles to learn representations that can be used across tissues, datasets, and biological tasks. In practice, researchers often use released checkpoints as frozen feature extractors to avoid the computational cost of fine-tuning^19–22^. The resulting cell embeddings can be used directly for clustering and batch assessment or combined with prediction heads for cell-type annotation and gene-expression reconstruction. However, these models differ in architecture, input representation, parameter count, pretraining objectives, external biological priors^23,24^, and pretraining species coverage. It is therefore difficult to know which model is best suited to a particular task. It also remains unclear when pretrained representations offer an advantage over methods fitted directly to the target dataset, such as principal component analysis (PCA)^25^, uniform manifold approximation and projection (UMAP)^26^, and scVI^27^.

Previous benchmarks have compared scFMs across several downstream tasks, but their conclusions differ across models and evaluation settings^19–22^. Some studies report benefits from pretraining, while others find that PCA and scVI remain competitive. These comparisons provide useful evidence, but they do not fully explain what biological information makes one representation more useful than another. In particular, overall rankings may hide differences in the preservation of rare cell types, gene-expression patterns, and continuous cellular states. The limits of generalization also remain unclear. Recent perturbation benchmarks show that simple baselines can compete with deep learning methods on unseen genetic perturbations^28,29^. Specialized methods such as GEARS, scGen, and CPA have been developed for perturbation prediction^30–32^, but the ability of general-purpose frozen scFM embeddings to support this task has not been systematically compared with their performance on tissue datasets. Other studies have examined how tokenization and pretraining data diversity affect individual model families^33,34^. However, it remains unclear whether model size and pretraining data volume predict performance across different released checkpoints. Another concern is the possible reuse of public datasets in both pretraining and evaluation. Without detailed pretraining records, it is difficult to establish whether benchmark datasets are independent of the training data.

These gaps raise four connected questions about the value and reliability of frozen scFM representations. First, what biological information do different models preserve, and how does this information affect their performance across tasks? Distinguishing broad cell-type identity from rare populations, gene-expression patterns, and continuous transcriptional programs may help explain why models perform differently across biological applications. Second, how well do pretrained representations generalize beyond their original training settings? Applying humanpretrained models to mouse tissues tests whether learned cellular information can be reused across species, while predicting unseen genetic perturbations tests whether these representations can also support the prediction of responses to interventions. Third, do larger models and larger pretraining datasets consistently produce better representations? Comparing released checkpoints with different architectures can help establish whether parameter count and pretraining data volume are useful indicators of performance and how performance differences relate to model design. Finally, how reliable are model rankings when pretraining data are not fully documented? Understanding the effects of incomplete training records and potential dataset overlap is necessary to judge whether reported advantages over classical baselines are robust.

To address these questions, we developed CellBench, a unified benchmark for evaluating frozen scFM representations across biological tasks and datasets. We compared 13 released scFM checkpoints with PCA, UMAP, and scVI fitted strictly on training data, covering 12 human and mouse tissue cohorts and four CRISPR Perturb-seq screens. Using consistent data splits and downstream evaluation methods, we assessed unsupervised clustering, native batch structure, supervised cell-type annotation, and gene-expression reconstruction. We also used linear probes to examine how well different embeddings preserve rare cell populations and continuous cellular programs. To investigate generalization, we evaluated human-pretrained checkpoints on mouse tissues and tested frozen representations for genetic perturbation prediction using a shared residual decoder. Perturbation predictions were additionally compared with a zerochange baseline to determine whether they improved on predicting no transcriptional response. We further examined model parameter composition, reported pretraining corpus size, and the relationship between checkpoint characteristics and downstream performance. Finally, we audited pretraining data provenance and tested the sensitivity of model rankings and performance advantages to the exclusion of cohorts with identified sample or donor matches in pretraining records. Our results show that different models preserve different biological information and that no single representation consistently leads across tasks. Classical baselines remain competitive, while representations that transfer to supervised mouse tasks do not necessarily support accurate prediction of unseen genetic interventions under the shared decoder. We also find that model scale does not reliably predict performance across heterogeneous checkpoints and that stable rankings can conceal substantial changes in performance advantages over classical methods. Together, CellBench provides a systematic assessment of what scFM representations capture, when they offer practical benefits, and how reliably those benefits can be demonstrated.

## RESULTS

### CellBench enables systematic benchmarking of scFMs and dataset-fitted representations

A major challenge in benchmarking single-cell foundation models (scFMs) is separating the utility of pretrained representations from the effects of model-specific fine-tuning strategies. To mitigate this challenge, we developed CellBench, a unified evaluation framework that compares pretrained cellular representations using standardized donor-level data splits (Supplemental Table S1), consistent downstream probe architectures (Supplemental Table S2), and rank-based performance aggregation across datasets and tasks (**Figure 1**; Supplemental Table S3). We evaluated 13 publicly released scFM checkpoints, including CellFM^13^, CellPLM^14^, GeneCompass^12^, Geneformer^5^, LangCell^17^, Nicheformer^10^, Tabula^18^, TranscriptFormer^9^, UCE^8^, scFoundation^7^, scGPT^6^, scLong^11^, and scPRINT2^15,16^, alongside three dataset-fitted baselines, including PCA^25^, UMAP^26^, and scVI^27^.

**Figure 1:**
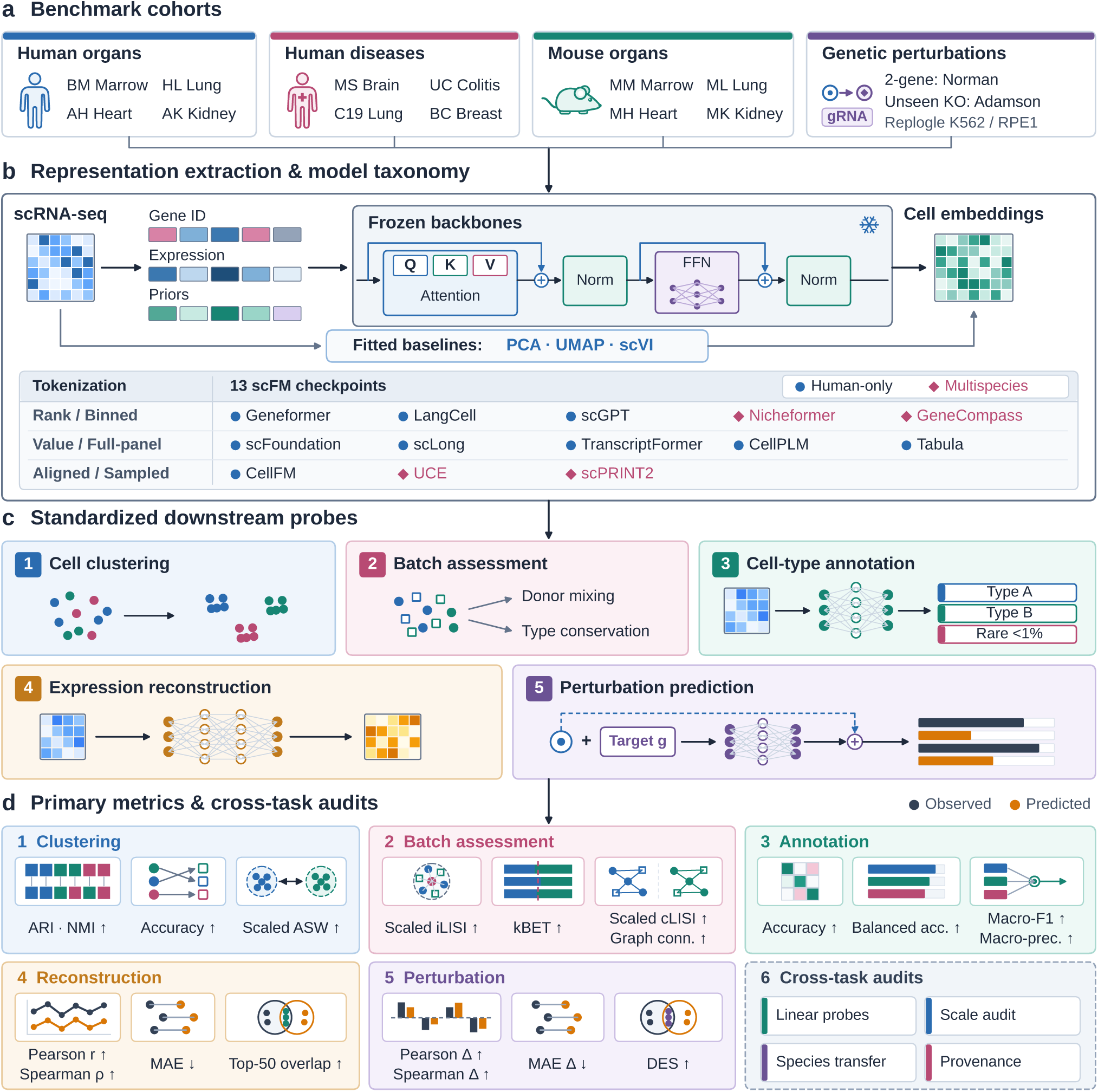
CellBench frozen-backbone evaluation. (a) Four evaluation panels: human organs, human diseases, matched mouse organs, and CRISPR Perturb-seq screens. (b) Thirteen pretrained checkpoints and three training-split baselines. (c) Shared probes for clustering, native batch assessment, annotation, reconstruction, and perturbation decoding. (d) Equal-weight ranking across four primary metrics per task (1–5), alongside cross-task audits of linear probes, scale attributes, cross-species transfer, and provenance (6).

CellBench assesses four complementary aspects of representation quality through unsupervised clustering^35^, batch-effect assessment in the native embedding space^37^, supervised closed-set cell-type annotation, and supervised reconstruction of highly variable gene (HVG) expression. We evaluated these tasks across 12 cohorts organized into three biological panels, encompassing healthy human organs, human diseases, and anatomically matched mouse organs. The healthy human panel included bone marrow mononuclear cells (BM)^38^, the Human Lung Cell Atlas (HL)^1^, adult heart (AH)^39^, and adult kidney (AK)^40^. The disease panel comprised multiple sclerosis brain (MS)^41^, ulcerative colitis epithelium (UC)^42^, lethal COVID-19 lung (C19)^43^, and triple-negative breast cancer (BC)^44^. The mouse panel included bone marrow (MM), lung (ML), heart (MH), and kidney (MK) from Tabula Muris Senis^2^ (Supplemental Table S4). To ensure a fair comparison and avoid information leakage during baseline fitting, the dataset-fitted baselines selected up to 3,000 HVGs and learned 256-dimensional representations exclusively from the 70% donor-partitioned training set for each random seed, without using cell-type labels or batch covariates. The fitted baselines were then applied to held-out validation (10%) and test (20%) donors through out-of-sample transformation or inference (Supplemental Table S1).

To examine whether representation quality observed in cellular atlases extends to experimental perturbations, we evaluated the same 16 methods on an independent panel of four CRISPR Perturb-seq datasets, comprising Norman^45^, Adamson^46^, Replogle K562, and Replogle RPE1^47^. Perturbation results were analyzed separately and excluded from both static representation rankings and cross-checkpoint attribute correlation analyses, so that observational and perturbation performance could be assessed independently.

### Pretrained models preserve distinct aspects of biological information

This section examines whether pretrained embeddings from different models capture a common dimension of biological representation quality or distinct biological properties across four static tasks. We evaluated unsupervised clustering and native batch effects directly in the embedding space without training task-specific heads, while using standardized task-specific prediction heads for closed-set annotation and HVG reconstruction. We further applied Ridge classification and regression probes for rare-class and gene-program information, without using an MLP.

Composite ranks showed that no scFM or dataset-fitted baseline consistently led across all comparisons, suggesting that different encoders capture distinct aspects of biological information (Supplemental Tables S5–S20).

#### Unsupervised clustering

This task tests whether embeddings recover major tissue lineages and cell-state neighborhoods without labels. Composite rankings differed across biological panels, with protein-language UCE^23^ leading healthy human organs at mean rank 2.604 and human diseases at 3.771, whereas a 256-component PCA fitted only on training-split HVGs led mouse organs at mean rank 2.500 versus 3.188 for UCE and outperformed UCE on the multidonor HL dataset at 2.667 versus 4.750. That overall ordering cannot distinguish agreement with annotated cell-type partitions from continuous separation between cell types (**Figure 2**; Supplemental Figures S1–S4). We therefore considered two biological interpretations, scoring Louvain partition agreement with annotated types by three metrics, including the unsupervised adjusted Rand index (uARI), normalized mutual information (NMI), and Hungarian matching accuracy, while scoring continuous cell-type separation by cell-type average silhouette width (ASW). However, the two interpretations identified different leading models. PCA reached the highest 12- cohort NMI of 0.706, ahead of UCE at 0.701, and uARI and matching accuracy followed the same order. These results suggest that PCA on HVGs can recover major cell-type partitions without requiring pretrained representations. By contrast, continuous separation told a different story. Dataset-fitted UMAP had the largest 12-cohort ASW of 0.578, with UCE next at 0.572. These results suggest that UCE preserves competitive continuous cell-type separation without consistently improving discrete clustering over dataset-fitted baselines.

**Figure 2:**
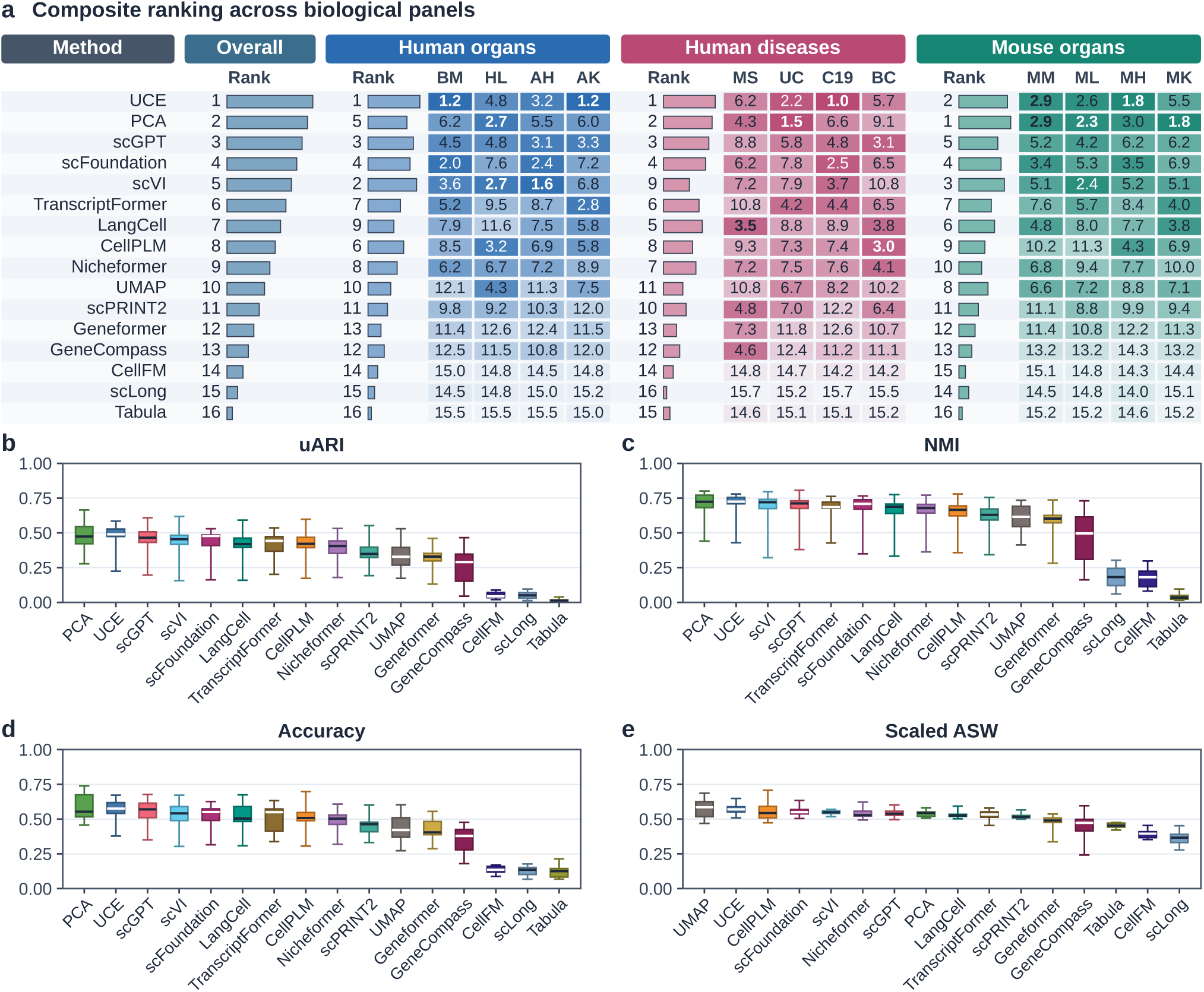
Unsupervised clustering. (a) Composite 1–16 ranking table across 12 cohorts. (b–e) Dataset-mean score distributions for uARI, NMI, matching accuracy, and scaled ASW.

#### Native batch assessment

This task tests whether embeddings can mix cells from different donors without collapsing cell-type structure. Composite ranks placed PCA first on human organs at 6.042, CellPLM first on human diseases at 7.594, and TranscriptFormer and UCE jointly first on mouse organs at 6.917. That overall ordering cannot tell whether a model mixes donors or conserves cell types (**Figure 3**; Supplemental Figures S5–S8). We therefore considered two biological interpretations, scoring donor mixing by scaled iLISI^48^ and kBET^49^, while scoring cell-type structure conservation by scaled cLISI^37,48^ and graph connectivity^37^. However, the two interpretations revealed different model strengths. On human organs, CellPLM showed greater donor mixing than UCE, with iLISI 0.310 versus 0.215, and kBET followed the same order at 0.325 versus 0.180. By contrast, cell-type conservation told a different story. UCE conserved cell types better than CellPLM, with scaled cLISI 0.989 versus 0.974, and graph connectivity followed the same order at 0.906 versus 0.782. Furthermore, high donor mixing coincided with poor clustering performance in several models. Tabula, CellFM, and mean-gene-pooled scLong showed high iLISI scores of 0.359, 0.303, and 0.297, respectively, yet showed some of the lowest clustering uARI scores across 12 cohorts, at 0.016, 0.052, and 0.051 (**Figure 2**). This pattern suggests that elevated donor mixing in these unconditioned embeddings may reflect broad loss of biological structure rather than selective removal of technical batch effects.

**Figure 3:**
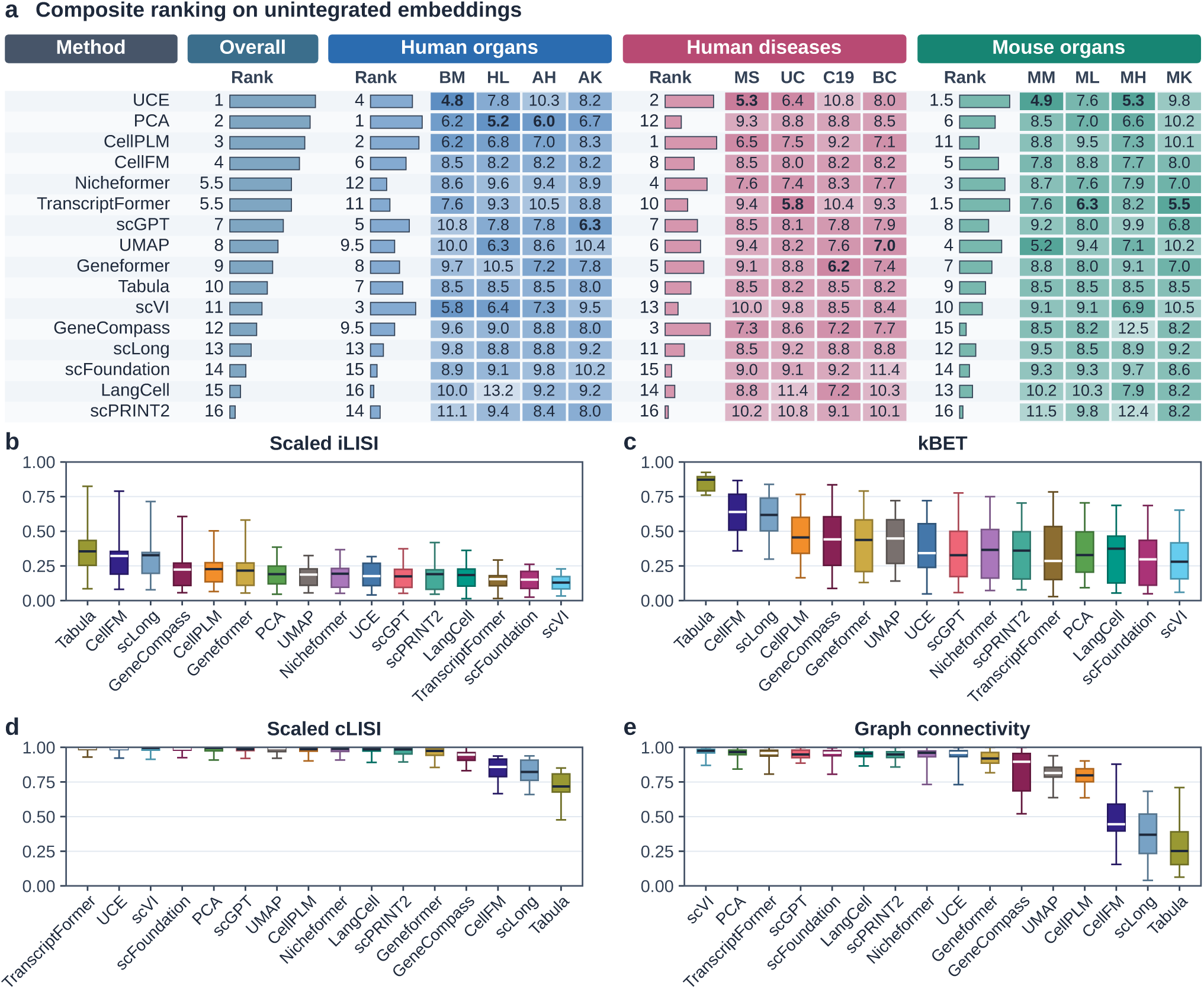
Native batch assessment. (a) Composite 1–16 ranking table on unintegrated embeddings. (b–e) Dataset-mean score distributions for scaled iLISI, kBET, scaled cLISI, and graph connectivity.

#### Supervised cell-type annotation

This task tests whether embeddings preserve fine-grained cell-type identity, including rare populations whose performance may be obscured by abundant lineages in overall accuracy. Full-panel scFoundation led all three panels on composite rank, with mean ranks of 2.854 on human organs, 2.156 on human diseases, and 1.979 on mouse organs. That overall ordering cannot distinguish gains in overall accuracy from improvements in rare-sensitive classification performance (**Figure 4**; Supplemental Figures S9–S12). We therefore considered two biological interpretations, scoring overall classification performance by overall accuracy, while assessing class-balanced performance using macro-F1, balanced accuracy, macro-precision, and macro-AUPRC. However, the two interpretations revealed different magnitudes of improvement. Relative to training-split PCA, scFoundation achieved only a 0.0074 advantage in 12-cohort overall accuracy, at 0.8816 versus 0.8742. By contrast, its macro-F1 advantage was more than three times larger, at 0.0234, with scores of 0.8334 versus 0.8099. Balanced accuracy and macro-precision followed the same order, while macro-AUPRC was 0.0267 higher, at 0.8719 versus 0.8452. scFoundation also exceeded scVI by 0.0126 in macro-F1 (Supplemental Tables S20 and S21). These results suggest that scFoundation’s advantage over PCA is more pronounced in class-balanced metrics than in overall accuracy, consistent with better preservation of rare cell-type information. However, the BM dataset provided an informative exception. In this dataset, 45 hematopoietic states were defined using CITE-seq surface-protein gating together with mRNA measurements. Training-split PCA outperformed scVI and every scFM on macro-F1, 0.5818 versus 0.5365 for scVI and 0.5309 for scFoundation. This exception suggests that locally fitted HVG representations can remain competitive for cell-state annotations informed by surface-protein measurements.

**Figure 4:**
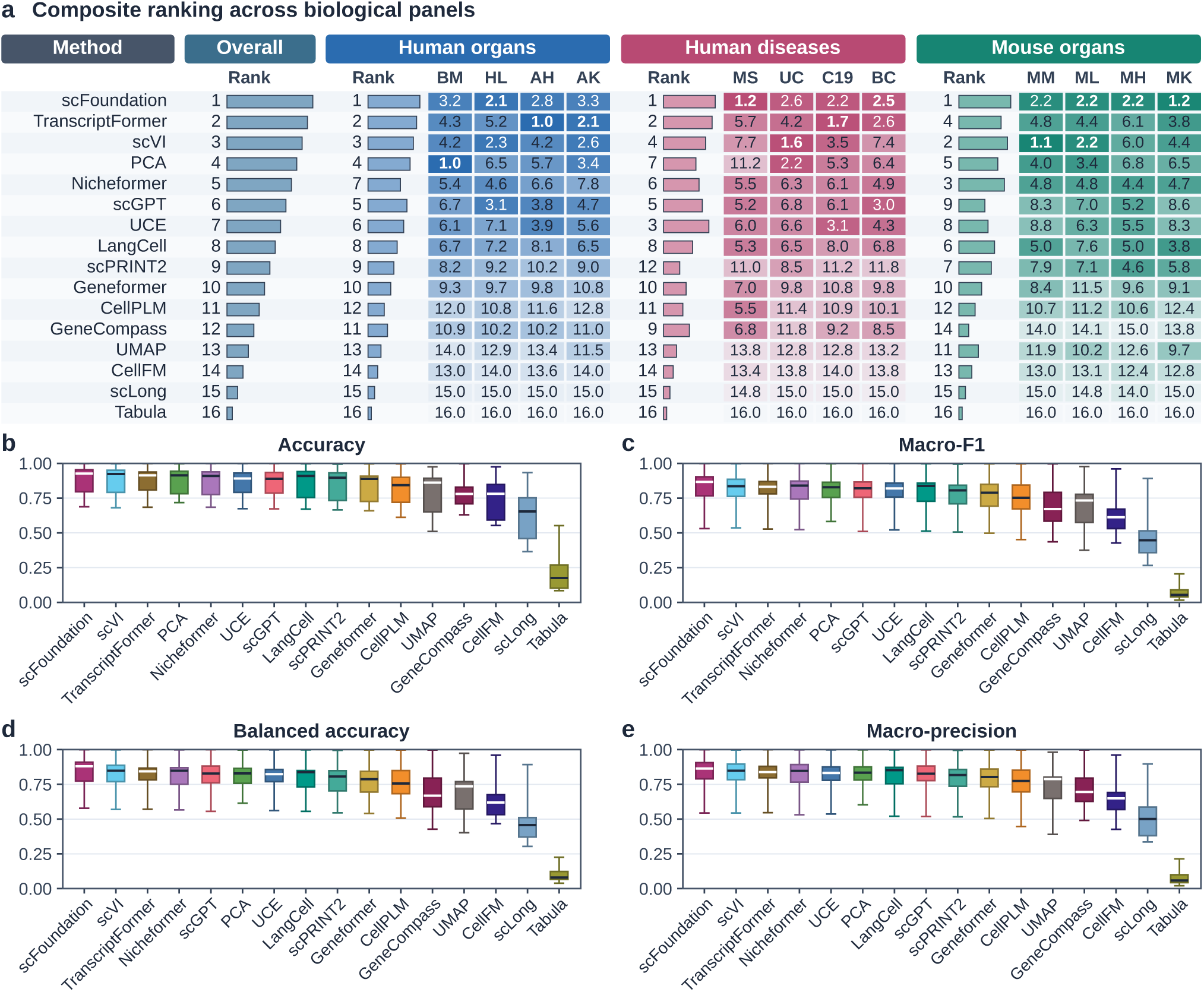
Cell-type annotation. (a) Composite 1–16 ranking table across human organs, human diseases, and mouse organs. (b–e) Dataset-mean score distributions for accuracy, macro-F1, balanced accuracy, and macro-precision.

#### Supervised expression reconstruction

This task tests how well cell embeddings preserved quantitative gene-expression information by reconstructing log-normalized highly variable gene (HVG) expression with a shared MLP decoder. Training-split PCA consistently led the composite rankings across human organs, human diseases, and mouse organs, with mean ranks of 1.000, 1.000, and 1.312, respectively, and an overall 12-cohort mean rank of 1.10 (**Figure 5**; Supplemental Figures S13–S16). Dataset-fitted scVI ranked second overall at 3.32, followed by

**Figure 5:**
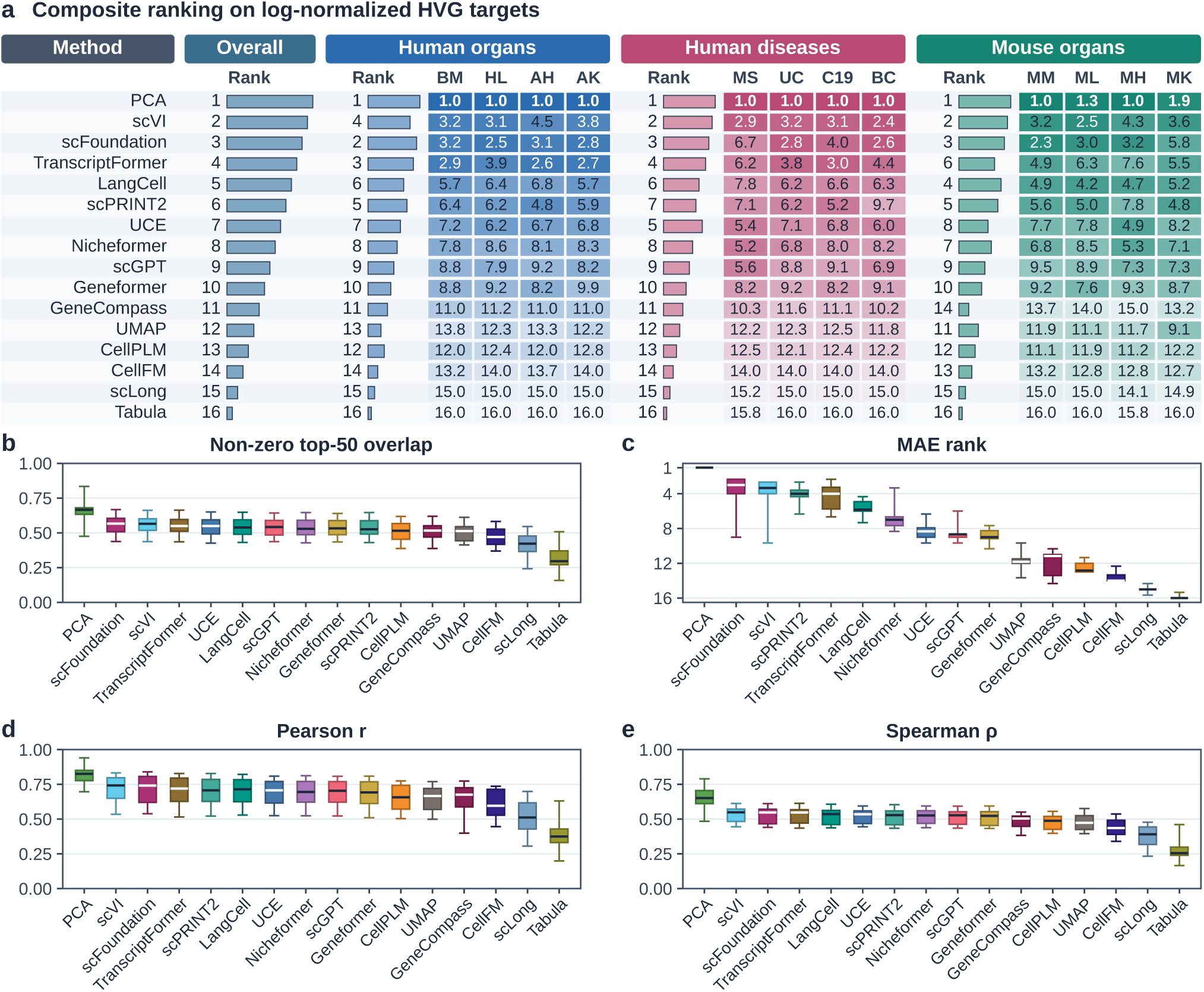
HVG expression reconstruction. (a) Composite 1–16 ranking table across human organs, human diseases, and mouse organs. (b–e) Dataset-mean score distributions on lognormalized HVG expression for non-zero top-50 overlap, MAE rank, per-cell Pearson *r*, and Spearman *ρ*.

scFoundation at 3.49 and TranscriptFormer at 4.48. To distinguish reconstruction of individual cell profiles from recovery of gene-specific expression variation across cells, we compared within-cell metrics (per-cell Pearson *r*, Spearman *ρ*, MAE, and non-zero top-50 overlap) with gene-wise Pearson correlation *r*_gene_ (Supplemental Tables S11, S15, S19, and S20). The within-cell metrics produced highly concordant model rankings, with a mean Spearman rank agreement of 0.968 between per-cell Pearson *r* and MAE when oriented consistently. PCA showed a substantial advantage on these measures, achieving a 12-cohort mean Pearson *r* of 0.8151, non-zero top-50 overlap of 0.6593, and MAE of 0.0593. By comparison, scFoundation and scVI achieved similar but lower within-cell fidelity, with Pearson *r* values of 0.7159 and 0.7174, respectively. However, this performance gap largely disappeared when evaluating gene-specific expression variation across cells. PCA achieved a mean *r*_gene_ of 0.4141, compared with 0.4093 for scVI and 0.4077 for scFoundation. Both scVI and scFoundation exceeded PCA on this metric in 6 of the 12 cohorts, indicating that their weaker reconstruction of complete cell profiles did not necessarily imply poorer recovery of expression variation for individual genes.

#### Linear rare-class probe

We applied five-fold cell-stratified Ridge classifiers to cached em-beddings from 1,000 cells sampled from each of the HL, MS, C19, and MH datasets to assess the linear accessibility of abundant and rare cell-type information without using an MLP (Supplemental Table S22). Across 26 abundant cell types representing at least 5% of their respective cohorts, training-split PCA achieved an F1 score of 0.844 and scVI reached 0.854, comparable to the strongest scFMs at 0.843 to 0.883. By contrast, larger differences emerged across 11 rare cell types representing less than 1% of their respective cohorts, including differentiating lymphatic endothelia and activated smooth muscle in the HL dataset, brush and mesothelial cells in the C19 dataset, and cardiac neurons and mast cells in the MH dataset. scFoundation achieved an F1 score of 0.735, followed by TranscriptFormer at 0.734 and UCE at 0.726. These three checkpoints outperformed PCA at 0.536 and scVI at 0.471 by approximately 19 to 26 percentage points. To check whether this gap reflected embedding dimensionality, single subsamples, or cell-level splitting, we repeated the probe across six sensitivity regimes spanning native versus PCA-192-matched embeddings, RidgeCV regularization, 10 repeated subsamples, and donor-grouped cross-validation on 3,000 cells per cohort (Supplemental Table S22). Under the strictest regime combining 3,000 cells, donor-grouped splits, and matched 192-dimensional em-beddings, scFoundation, UCE, scGPT, and TranscriptFormer retained rare-class F1 scores of 0.582, 0.569, 0.522, and 0.512, remaining well ahead of scVI at 0.287, training-split PCA at 0.196, and direct 192-dimensional PCA on log-normalized HVGs at 0.102. These results suggest that dataset-fitted PCA and scVI remain competitive for abundant cell types, whereas several pretrained foundation models retain more linearly accessible information for distinguishing rare cell populations.

#### Linear program probe

Cell-type identity is not the only biological information encoded in these embeddings. We therefore applied the same five-fold Ridge regression setup to assess the linear accessibility of continuous transcriptomic attributes and biological programs from frozen embeddings, including library depth, ribosomal and mitochondrial transcript fractions, cell-cycle proliferation, and interferon signaling (Supplemental Table S22). Because scFoundation receives total read depth as an input token, it achieved an expectedly high *R*^2^ of 0.990 for log library depth. It also achieved *R*^2^ values of 0.838 for ribosomal transcript fractions across the four probed cohorts (HL, MS, C19, and MH) and 0.847 for mitochondrial fractions across HL and C19. By contrast, scFoundation showed lower performance for cell-cycle proliferation and interferon signaling, with *R*^2^ values of 0.361 and 0.296. Among pretrained checkpoints, GeneCompass, which incorporates regulatory priors, led cell-cycle proliferation with an *R*^2^ of 0.510, ahead of scGPT at 0.412 and LangCell at 0.389, while text-aligned LangCell led interferon signaling at 0.352. Meanwhile, TranscriptFormer and UCE nearly matched scFoundation in rare-lineage classification, with F1 scores of 0.734 and 0.726 versus 0.735 for scFoundation. However, they showed substantially weaker recovery of mitochondrial transcript fractions, with *R*^2^ values of *−*0.408 and 0.318, and interferon signaling, with values of *−*0.334 and 0.039. These results suggest that linear accessibility of rare cell-type identity does not necessarily imply accurate recovery of other transcriptomic attributes or biological programs.

#### Cross-task concordance

We examined cross-model Spearman correlations across the 10 scFMs that did not collapse during unsupervised clustering, excluding Tabula, CellFM, and scLong (Supplemental Table S23). First, cell-type annotation showed strong concordance with gene-expression pattern reconstruction. Macro-F1 correlated strongly with gene-wise reconstruction performance *r*_gene_ at +0.939 and non-zero top-50 expressed-gene overlap at +0.879, whereas its association with quantitative expression accuracy (*−*MAE) was weaker at +0.707. This pattern suggests that annotation performance is more closely aligned with recovering geneexpression patterns than with accurately predicting expression magnitudes. Second, clustering performance showed concordance with cell-type neighborhood structure, gene-expression pattern recovery, and annotation. Louvain uARI correlated with ASW at +0.745 and scaled cLISI at +0.806. Beyond these clustering-related metrics, non-zero top-50 expressed-gene overlap correlated strongly with scaled cLISI at +0.782 and uARI at +0.758, while macro-F1 was also positively associated with scaled cLISI at +0.745 and uARI at +0.612. These associations indicate that models preserving cell-type neighborhood structure also tend to perform well in expression-pattern reconstruction and cell-type annotation. Third, donor mixing showed negative concordance with annotation and reconstruction performance. iLISI correlated negatively with macro-F1 at *−*0.796 and *r*_gene_ at *−*0.778. These inverse associations suggest that stronger donor mixing can coincide with weaker preservation of cell-type identity and gene-expression information, highlighting a potential tension between batch mixing and biological information preservation. Finally, quantitative expression accuracy showed limited concordance with clustering performance. Although *−*MAE was moderately associated with macro-F1 (+0.707), it showed little association with unsupervised cluster geometry, correlating with uARI at +0.280 and ASW at +0.079. This contrast suggests that accurately predicting expression magnitudes captures aspects of representation quality that are not strongly reflected in clustering performance. Together, these results suggest that cell-type identity, gene-expression reconstruction, clustering geometry, and donor mixing capture related but distinct properties of pretrained cellular representations, and that strong performance in one task does not necessarily translate into strong performance in another.

### Cross-species transfer to mouse exceeds generalization to unseen genetic perturbations

The four static tasks above evaluated pretrained representations on held-out donors within each study. These results do not establish whether the same embeddings remain informative when transferred across species or evaluated on unseen genetic perturbations. We therefore examined two distinct distribution shifts: cross-species transfer from human to mouse organs (**Table 1**; Supplemental Table S24) and prediction of transcriptional responses to held-out genetic perturbations (**Figure 6**; Supplemental Table S25).

**Figure 6:**
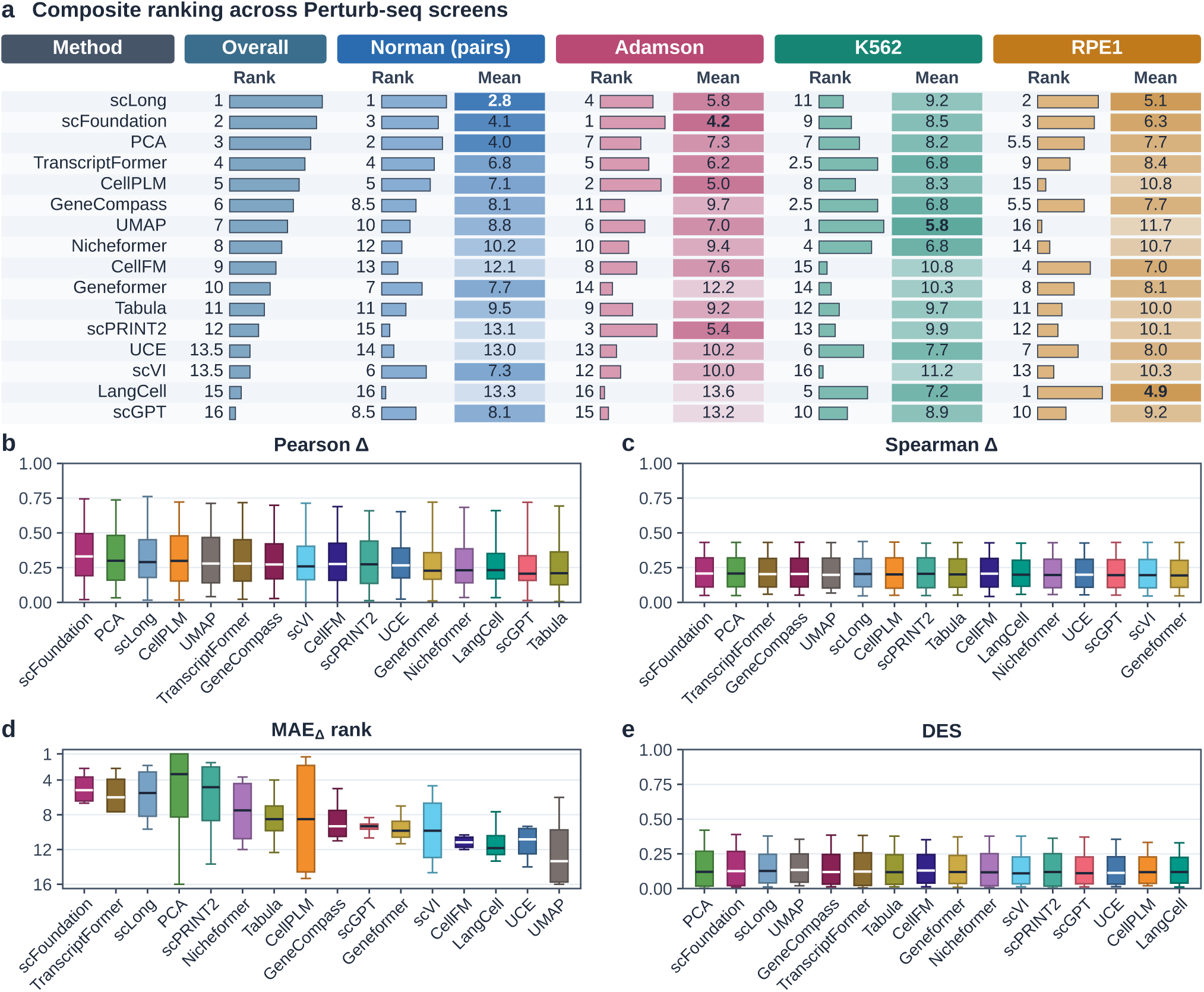
Perturbation prediction. (a) Composite 1–16 ranking table across four Perturbseq screens. (b–e) Four-screen distributions of Pearson Δ, Spearman Δ, MAE_Δ_ rank, and differential-expression overlap score, DES.

**Table 1:** Paired human-organ and mouse-organ ranks.

|  | Overall |  | Clustering |  | Batch |  | Annotation |  | Reconstruction |  |
|---|---|---|---|---|---|---|---|---|---|---|
| Method | Human | Mouse | Human | Mouse | Human | Mouse | Human | Mouse | Human | Mouse |
| scFoundation | 5.02 | **4.90** | 4.81 | **4.79** | 9.52 | **9.23** | 2.85 | **1.98** | **2.90** | 3.58 |
| TranscriptFormer | **5.44** | 6.05 | 6.54 | **6.42** | 9.04 | **6.92** | **3.17** | 4.78 | **3.02** | 6.08 |
| UCE | **5.70** | 6.11 | **2.60** | 3.19 | 7.79 | **6.92** | **5.67** | 7.23 | **6.75** | 7.12 |
| LangCell | 7.97 | **6.33** | 8.21 | **6.04** | 10.44 | **9.15** | 7.12 | **5.36** | 6.12 | **4.75** |
| Nicheformer | 7.66 | **6.97** | **7.23** | 8.46 | 9.12 | **7.79** | 6.10 | **4.68** | 8.19 | **6.94** |
| scGPT | **6.29** | 7.38 | **3.92** | 5.48 | **8.17** | 8.48 | **4.56** | 7.28 | 8.52 | **8.27** |
| scPRINT2 | 8.64 | **8.11** | 10.33 | **9.81** | **9.23** | 10.48 | 9.15 | **6.36** | 5.83 | **5.77** |
| Geneformer | 9.94 | **9.49** | 11.98 | **11.44** | 8.81 | **8.21** | 9.90 | **9.65** | 9.06 | **8.69** |
| CellPLM | **9.32** | 9.98 | **6.10** | 8.21 | **7.08** | 8.92 | 11.79 | **11.21** | 12.29 | **11.58** |
| CellFM | 12.61 | **12.09** | 14.77 | **14.65** | 8.31 | **8.04** | 13.65 | **12.81** | 13.73 | **12.88** |
| GeneCompass | **10.55** | 12.77 | **11.71** | 13.52 | **8.83** | 9.38 | **10.60** | 14.23 | **11.04** | 13.96 |
| scLong | 13.50 | **13.27** | 14.85 | **14.58** | 9.15 | **9.04** | 15.00 | **14.71** | 15.00 | **14.75** |
| Tabula | 13.94 | **13.88** | 15.38 | **15.06** | **8.38** | 8.50 | 16.00 | 16.00 | 16.00 | **15.96** |
| PCA | **4.07** | 4.27 | 5.08 | **2.50** | **6.04** | 8.08 | **4.15** | 5.19 | **1.00** | 1.31 |
| scVI | **4.47** | 5.05 | **3.67** | 4.46 | **7.25** | 8.90 | **3.33** | 3.43 | 3.62 | **3.42** |
| UMAP | 10.88 | **9.35** | 8.81 | **7.40** | 8.83 | **7.98** | 12.96 | **11.10** | 12.92 | **10.94** |

Rows within each block follow Mouse Overall rank (unweighted mean of the four task ranks across four anatomically matched organs). Boldface marks the stronger (lower) rank between human and mouse within each task for a given method (ties unbolded). Dataset-fitted baselines are listed below the rule.

#### Paired mouse transfer

This comparison tests whether checkpoints pretrained mainly on human cells remain competitive on mouse organs without mouse pretraining (**Table 1**). We therefore compared supervised annotation and reconstruction against unsupervised clustering across four anatomically matched human and mouse tissues, including bone marrow, lung, heart, and kidney (**Table 1**; mouse-minus-human annotation rank shifts in Supplemental Table S24). For supervised tasks, model rankings on mouse datasets aligned more closely with the corresponding checkpoints performance in human-organ annotation and reconstruction than with their mouse pretraining coverage. Predominantly human-pretrained scFoundation led mouse annotation at 1.979 and led pretrained scFMs in mouse reconstruction at 3.583 (behind training-split PCA at 1.312 and scVI at 3.417), ranking ahead of human-and-mouse Nicheformer at 4.677, multispecies scPRINT2 at 6.365, eight-species UCE at 7.229, and human-and-mouse GeneCompass at 14.229 in annotation. Similarly, TranscriptFormer’s human-only TF-Sapiens checkpoint achieved a mouse annotation rank of 4.781, closely matching Nicheformer’s rank of 4.677. Specifically, the same pattern held in the mixed-chemistry MH cohort combining 10x Genomics and Smart-seq2 data. In this dataset, scFoundation reached the highest full-cohort macro-F1 of 0.921, ahead of Nicheformer at 0.909, PCA at 0.887, and GeneCompass at 0.436 (Supplemental Table S17), while TranscriptFormer led linear probing of rare cardiac neurons and mast cells at 0.857 (Supplemental Table S22). By contrast, unsupervised mouse clustering told a different story (**Figure 2**). Training-split PCA fitted on mouse HVGs led mouse clustering at 2.500, ahead of UCE at 3.188 and scFoundation at 4.792, while scGPT dropped from 3.917 on human organs to 5.479 on mouse organs and CellPLM dropped from 6.104 to 8.208. These results suggest that human-pretrained embeddings preserve enough shared gene-expression structure for supervised mouse annotation and reconstruction, whereas dataset-fitted PCA remains stronger for unsupervised mouse clustering.

#### Perturbation prediction

This task tests whether frozen control-cell embeddings can predict transcriptional shifts (Δ) after held-out genetic perturbations when paired with a shared residual lookup decoder (**Figure 6**; Supplemental Figure S17; Supplemental Table S25). Across four Perturb-seq screens scored by Pearson Δ, Spearman Δ, genome-wide MAE_Δ_, and differentialexpression overlap score (DES), scLong achieved the best composite rank at 5.729, closely followed by scFoundation at 5.771 and training-split PCA at 6.792. However, this perturbation ranking did not match the static tissue rankings, and in several cases reversed them. scLong led perturbation rankings despite placing 15th in human-organ annotation and showing collapsed multi-lineage clustering across the 12 tissue cohorts at a uARI of 0.051, whereas top tissue clustering models such as UCE and scGPT ranked 13th and 16th at 9.708 and 9.875. This contrast may reflect differences in the biological variation captured by the two tasks. While tissue benchmarks evaluate the separation of multiple cell lineages, each Perturb-seq screen measures transcriptional responses within a single clonal cell line, either K562 or RPE1. Moreover, the shared residual decoder combines frozen cell embeddings with a 128-dimensional perturbation vector, allowing perturbation prediction to draw on information beyond the cell embeddings alone. Thus, models that perform poorly at separating tissue lineages may still retain information useful for predicting perturbation responses. However, relative rankings across the 16 methods do not establish whether their predicted shifts outperform a baseline that predicts no transcriptional change.

#### Zero-change control

We therefore evaluated perturbation predictions against a zero-change baseline (Δ̂*_g_* = **0**) that predicts no expression change relative to the unperturbed control mean^28,29^ (Supplemental Table S25). This comparison revealed contrasting results between squared-error and absolute-error metrics, particularly for unseen combinations of previously observed perturbation targets. On Norman, the decoder was trained on all 105 constituent single-gene perturbations and evaluated on 118 held-out two-gene combinations. Across all 16 methods, three-seed mean Pearson Δ scores ranged from 0.652 to 0.761, while MSE_Δ_ skill was consistently positive, ranging from +0.4078 to +0.6267. Training-split PCA achieved a three-seed mean MSE_Δ_ skill of +0.6157. On seed 41, all 16 methods also achieved positive MSE_Δ_ skill on top differentially expressed genes (+0.4840 to +0.5981 on DE-20; +0.4824 to +0.6253 on DE-100), exceeding the training-mean shift baseline (+0.3180 genome-wide, +0.3121 on DE-20, and +0.3187 on DE- 100). By contrast, genome-wide absolute-error performance told a different story. Run-level MAE_Δ_ skill ranged from *−*0.1261 to +0.0144, with positive values in only 3 of 48 runs. None of the 16 methods achieved a positive three-seed mean skill, trailing the training-mean baseline (+0.2543), while summing the two single-gene training shifts reached +0.7273 in MSE_Δ_ and +0.4086 in MAE_Δ_, outperforming all 16 methods on both metrics. This discrepancy may re-flect small prediction errors across many genes with little true expression change, which can contribute to genome-wide absolute error despite improvements in squared-error performance. Together, these results indicate that models can recover aspects of the transcriptional responses to unseen combinations of previously observed perturbation targets and reduce squared prediction errors without consistently outperforming a zero-change baseline in genome-wide absolute error.

#### Unseen perturbation targets

Predicting responses to perturbation targets absent from decoder training was substantially more challenging across Adamson, Replogle K562, and Replogle RPE1. None of the 16 methods outperformed the zero-change baseline in genome-wide MAE_Δ_, with skill scores ranging from *−*1.4408 to *−*0.3496. On both genome-scale Replogle screens, all methods also performed worse than the zero-change baseline in squared error, with MSE_Δ_ skill ranging from *−*1.0463 to *−*0.0296. Across-gene expression-change correlations were close to zero on Replogle K562, with three-seed mean Pearson Δ ranging from *−*0.013 to +0.041. Restricting evaluation to the top differentially expressed genes did not reverse this pattern. On seed 41, all 16 methods retained negative MSE_Δ_ skill on K562, ranging from *−*0.7447 to *−*0.2494 for DE-20 and from *−*0.8483 to *−*0.3066 for DE-100. On RPE1 and Adamson, the highest DE-20 skill reached only +0.0438 and +0.2127, respectively. In contrast, the training-mean shift baseline achieved positive genome-wide MSE_Δ_ skill ranging from +0.0316 to +0.1737 and MAE_Δ_ skill ranging from +0.0160 to +0.1156 across all three screens. This baseline also outperformed all 16 methods on DE-20, with skill values ranging from +0.0352 to +0.2179, and on DE-100, with values ranging from +0.0354 to +0.1906 (Supplemental Table S25). Importantly, the shared lookup decoder represented unseen perturbation targets using untrained condition vectors sampled from *N* (0, 1), without incorporating external target-specific information from Gene Ontology or protein-language priors. These results indicate that, under this target-uninformed decoder, frozen control-cell embeddings combined with untrained perturbation condition vectors were insufficient to outperform the zero-change baseline in predicting genome-wide transcriptional responses to unseen genetic perturbations.

### Model scale alone does not reliably predict representation quality

To investigate whether model scale explains differences in frozen representation quality, we examined the relationships between audited parameter counts, reported pretraining corpus sizes, and performance across the four static tasks. We further assessed parameter composition, extracted embedding width, and pretraining design to explore other model characteristics associated with ownstream performance (**Table 2**, Supplemental Figures S18–S19, and Supplemental Table S26).

**Table 2:** Checkpoint attributes and four-task ranks.

| Method | Corpus | Params | Width | Input | Clustering | Batch | Annotation | Reconstruction |
|---|---|---|---|---|---|---|---|---|
| scFoundation | 50 | 119 | 3072 | F+V | 5.11 | 9.48 | 2.33 | 3.49 |
| TranscriptFormer | 57 | 430 | 2048 | V | 6.49 | 8.23 | 3.83 | 4.48 |
| Nicheformer | 110 | 39 | 512 | R+C | 7.42 | 8.23 | 5.49 | 7.40 |
| scGPT | 33 | 51 | 512 | B | 5.00 | 8.24 | 5.70 | 8.12 |
| UCE | 36 | 849 | 1280 | S+P | 3.19 | 7.44 | 5.97 | 6.74 |
| LangCell | 28 | 39 | 256 | R | 6.83 | 9.67 | 6.38 | 5.86 |
| scPRINT2 | – | 102 | 424 | A+C | 9.25 | 9.92 | 8.70 | 6.21 |
| Geneformer | 30 | 40 | 512 | R | 11.34 | 8.30 | 9.62 | 8.81 |
| CellPLM | – | 66 | 512 | V | 7.03 | 7.86 | 10.83 | 12.06 |
| GeneCompass | 102 | 18 | 256 | R+V | 11.68 | 8.63 | 11.31 | 11.94 |
| CellFM | 100 | 797 | 1536 | A | 14.63 | 8.20 | 13.41 | 13.53 |
| scLong | 48 | 1,006 | 200 | F+V | 14.99 | 9.00 | 14.89 | 14.94 |
| Tabula | – | 13 | 192 | V | 15.15 | 8.42 | 16.00 | 15.97 |
| scVI | – | – | 256 | HVG | 5.18 | 8.43 | 3.93 | 3.32 |
| PCA | – | – | 256 | HVG | 4.32 | 7.66 | 5.21 | 1.10 |
| UMAP | – | – | 256 | HVG | 8.40 | 8.29 | 12.40 | 12.03 |

Corpus: reported pretraining profiles (millions); dashes mark family-only or unresolved checkpoint corpus estimates. Params: audited instantiated parameters (millions). Width: extracted cell-embedding dimension *d*. Overlapping input codes: A, gene-aligned; V, gene identity and values; R, ranked genes; C, context; S, sampled genes; P, protein-derived features; F, fixed gene panel; B, expression bins; HVG, dataset-fitted on training-split HVGs. Task columns report 12-cohort mean ranks (lower is better) separately for each task without averaging across tasks. Rows within each block are ordered by 12-cohort annotation mean rank; dataset-fitted baselines are shown below the rule.

#### Model parameter count

We first examined whether pretrained checkpoints with more parameters achieved better performance across the four static tasks. Across all 13 scFMs, audited parameter counts ranged from 13.2M to 1,006.1M. Despite this substantial variation, Spearman correlations between parameter count and task mean rank were weak, with values of *−*0.242 for clustering, *−*0.121 for annotation, *−*0.143 for reconstruction, and *−*0.228 for native batch assessment. Because lower mean ranks indicate better performance, negative correlations suggest that larger models tend to perform better, although this tendency was weak in our results. Moreover, excluding individual models sometimes reversed the direction of these correlations, indicating that the associations were sensitive to which checkpoints were included. This lack of a consistent advantage for larger models was also evident in individual model comparisons. For annotation across the 12 cohorts, scLong at 1,006.1M and CellFM at 797.4M achieved relatively poor mean ranks of 14.89 and 13.41, respectively. In contrast, the substantially smaller scFoundation at 119.3M and scGPT at 50.8M performed considerably better, with mean ranks of 2.33 and 5.70.

#### Parameter composition

To better understand why total parameter count may not fully reflect encoder capacity, we examined how parameters were distributed across different checkpoint components (Supplemental Table S3). Our audit included both trainable and frozen registered parameters but excluded module buffers. In UCE, 744.8M of its 848.9M audited parameters (87.7%) belonged to a frozen ESM-2 protein lookup table, while its four-layer transformer encoder contained 78.7M parameters. Similarly, scPRINT2 contained 85.8M frozen gene-encoder parameters and 16.1M trainable transformer parameters within its 101.9M total. In contrast, GeneCompass stored 155.4M regulatory-prior elements as module buffers, which were excluded from its audited parameter count of 18.1M. These differences illustrate why total registered parameter count does not directly reflect the size of the active cell-representation encoder.

#### Pretraining corpus size

Beyond model parameters, we next investigated whether larger pretraining corpora were associated with better downstream performance. Across the 10 checkpoints with available corpus-size estimates (Supplemental Table S3), Spearman correlations with task mean rank were +0.321 for clustering, *−*0.353 for native batch assessment, *−*0.079 for annotation, and +0.152 for reconstruction, providing little evidence of a consistent advantage from larger pretraining corpora. This pattern was further illustrated by Nicheformer, pretrained on more than 110M cells, and GeneCompass, pretrained on 101.8M cells, both of which ranked behind scFoundation, pretrained on more than 50M cells, in clustering, annotation, and reconstruction. Together, these findings indicate that neither parameter count nor reported pretraining corpus size alone provides a reliable basis for selecting checkpoints with strong downstream performance.

#### Embedding width

Having found no consistent performance advantage associated with model size or pretraining data volume, we turned to the dimensionality of extracted cell embeddings and its relationship with supervised annotation performance. Across the 13 scFMs, embedding width ranged from 192 to 3,072 dimensions and showed a stronger Spearman correlation with 12-cohort annotation macro-F1, at +0.675, compared with +0.121 for total parameter count. After excluding three models with collapsed representations, the correlation between embedding width and macro-F1 remained positive at +0.721. Consistent with this pattern, Tabula with *d* = 192 and scLong with *d* = 200 had the two worst annotation mean ranks despite a 76-fold difference in parameter count. Together, these observations suggest that extracted embedding width may be more informative than nominal model size when comparing supervised annotation performance, although these associations do not establish a causal effect of embedding dimensionality.

#### Pretraining design

Finally, we compared Geneformer and LangCell to explore how closely related encoders with similar parameter counts but different pretraining strategies performed across downstream tasks. LangCell initializes its 12-layer BERT cell encoder from Geneformer and retains rank-ordered gene tokenization and a 2,048-gene context window. Their audited cell-encoder parameter counts were also similar, at 39.6M for Geneformer and 39.4M for Lang- Cell. However, LangCell incorporates additional cell–cell and cell–text contrastive learning and cell–text matching alongside masked gene modeling, using paired cell descriptions during pretraining. Despite their similar encoder sizes, LangCell achieved better 12-cohort mean ranks than Geneformer across clustering, improving from 11.34 to 6.83, annotation from 9.62 to 6.38, and reconstruction from 8.81 to 5.86. At the metric level, uARI increased by 27.6% from 0.326 to 0.416, while cell-type ASW increased from 0.483 to 0.531. Direct linear probes showed similar improvements, with out-of-fold *R*^2^ increasing from 0.192 to 0.389 for proliferation and from 0.101 to 0.352 for interferon signaling. However, performance declined on the de-confounded canonical ISG score, with *R*^2^ values of *−*0.466 for Geneformer and +0.056 for LangCell (Supplemental Tables S20 and S22). This improvement did not extend to native batch assessment, where the mean rank increased from 8.30 for Geneformer to 9.67 for LangCell. Because the two checkpoints also differ in pretraining data and supervision objectives, this comparison cannot isolate the contribution of any single training component. Nevertheless, it illustrates that closely related encoders with similar parameter counts can produce substantially different downstream representations.

### Pretraining overlap audits reveal stable rankings but sensitive annotation gains

Because public single-cell evaluation datasets may overlap with studies included in model pretraining corpora, apparent performance gains require careful examination of data provenance. We therefore audited potential pretraining overlap between each of the 13 scFM checkpoints and all 16 evaluation datasets, covering 208 model–dataset combinations (Supplemental Table S27). To assess ranking robustness, we performed leave-one-dataset-out (LODO) and leaveone-metric-out (LOMO) analyses across the 12 static cohorts (Supplemental Table S28). Finally, we examined how excluding the two human cohorts with confirmed sample or donor identifier matches in scFoundation’s pretraining records affected annotation rankings and macro-F1 differences relative to dataset-fitted baselines (Supplemental Table S29).

#### Pretraining provenance and documentation gaps

To assess the transparency of pretraining data sources, we classified all 208 combinations of scFM checkpoints and evaluation datasets into four mutually exclusive provenance categories (Supplemental Table S27). For 154 combinations (74.0%), sampleor donor-level overlap could not be determined because checkpoint-specific training manifests were unavailable. Another 28 combinations (13.5%) could be excluded based on documented species scope, corresponding to the seven strictly humanpretrained checkpoints (TranscriptFormer TF-Sapiens, scFoundation, scGPT, Geneformer, Lang- Cell, CellFM, and scLong) evaluated on the four mouse cohorts. A further 24 combinations (11.5%) involved shared public data sources, including CZ CELLxGENE Census^3^ and scBaseC-ount, but lacked the cell-level records needed to verify overlap; these included 8 combinations involving TranscriptFormer and human datasets, 6 involving scGPT and human datasets, and 10 involving scPRINT2 and human or mouse datasets. The remaining two combinations (1.0%) showed confirmed sample or donor identifier matches between scFoundation’s pretraining records and the evaluation datasets, comprising 19 matching sample identifiers for adult human heart (AH) and 6 matching donor identifiers for multiple sclerosis brain (MS). Notably, scFoundation also achieved its largest single-cohort macro-F1 advantages on MS, exceeding PCA by +0.1477 and scVI by +0.0910 (Supplemental Tables S13 and S21).

#### Rank robustness

We first assessed whether composite model rankings were sensitive to random seeds or to the removal of individual evaluation datasets or metrics. Across clustering, annotation, and reconstruction, rankings remained highly consistent across random seeds and under both LODO and LOMO analyses, with Spearman rank correlations ranging from 0.915 to 1.000 under LODO and from 0.968 to 1.000 under LOMO (Supplemental Figure S20 and Supple-mental Table S28). Perturbation rankings showed moderate sensitivity to screen removal (LODO 0.657 to 0.912; LOMO 0.823 to 0.970). However, native batch assessment was more sensitive to metric removal, with LOMO correlations ranging from 0.291 to 0.671 (Supplemental Figure S21 and Supplemental Table S28). This difference is consistent with the competing objectives of donor mixing and cell-type conservation, which are captured by different evaluation metrics.

#### Overlap-exclusion sensitivity

We next examined whether annotation rankings and macro-F1 advantages remained stable after excluding the two human cohorts with confirmed sample or donor identifier matches in scFoundation’s pretraining records. After removing AH and MS, scFoundation remained the top-ranked model in both human panels. Its mean annotation rank changed only slightly, from 2.854 to 2.889 for human organs and from 2.156 to 2.458 for human diseases (Supplemental Table S29). However, its macro-F1 advantage over dataset-fitted baselines decreased substantially. Across all 12 cohorts, scFoundation initially exceeded training-split PCA by +0.0234 in mean macro-F1, but this advantage fell by 51.9% to +0.0113 after excluding AH and MS. Its advantage over scVI similarly decreased from +0.0126 to +0.0041. Focusing on human cohorts alone, scFoundation’s advantage over PCA declined by 97.7%, from +0.0214 across eight cohorts to just +0.0005 across the remaining six. Mean macro-F1 after exclusion was 0.7949 for scFoundation, 0.7944 for PCA, and 0.7965 for scVI. In contrast, scFoundation maintained a +0.0274 advantage over PCA across the four mouse cohorts, with scores of 0.8861 and 0.8587, respectively. This mouse-cohort advantage accounted for nearly all of the remaining difference across the 10 retained cohorts. However, removing AH and MS also changes the tissue and cell-type composition of the evaluation datasets, so we cannot determine whether the reduced performance advantage was caused by pretraining overlap. Nevertheless, these results show that model rankings can remain stable even when the annotation advantage over classical baselines nearly disappears.

## DISCUSSION

The central finding of CellBench is that the value of a cellular representation depends on the biological information required by the downstream task, rather than on a single measure of model performance. Cell-type identity, continuous transcriptional programs, gene-expression patterns, and donor-related variation represent different aspects of cellular biology that are not necessarily captured equally well by the same embedding. This helps explain why previous benchmarks have sometimes reached different conclusions about the benefits of single-cell foundation models^19–22^. A model that performs well in supervised annotation may not be equally effective at unsupervised clustering or expression reconstruction, while strong donor mixing can coexist with substantial loss of cell-type structure. The continued competitiveness of dataset-fitted PCA and scVI further suggests that pretraining is not inherently advantageous for every biological task. Rather than identifying a universally superior checkpoint, benchmarking should therefore establish which biological properties different representations preserve and under what conditions those properties provide an advantage over simpler alternatives.

Model scale alone is also insufficient to explain what makes a cellular representation useful. Although larger models and broader pretraining datasets are often expected to yield more transferable features, this expectation does not translate into a consistent performance advantage across heterogeneous released checkpoints. This distinction is important when interpreting controlled studies that demonstrate the benefits of particular tokenization strategies or greater pretraining data diversity within individual model families^33,34^. Such experiments address the effect of specific design choices under relatively controlled conditions, whereas comparisons across released scFMs involve simultaneous differences in architecture, input representation, training data, and learning objectives. Moreover, total parameter count does not consistently describe the capacity of the cell encoder because checkpoints differ in how they allocate parameters to frozen lookup tables, trainable components, and other stored information. Our observations that embedding width was more strongly associated with annotation performance than parameter count, and that closely related Geneformer and LangCell encoders performed differently, further illustrate the limitations of using model size as a proxy for representation quality. These associations do not establish which architectural or training choices cause better performance. Instead, they motivate more controlled investigation of how input representations, pretraining objectives, and embedding design influence the biological information available to downstream tasks.

The ability to represent cellular states should not be equated with the ability to predict how those states will respond to genetic interventions. The competitive performance of several human-pretrained checkpoints on supervised mouse tasks suggests that useful cellular information can transfer across species, even when the pretraining data do not explicitly cover the target organism. However, this transfer remains task-dependent, as dataset-fitted representations continued to perform well in unsupervised mouse clustering. More importantly, the success of a frozen representation in observational tasks did not ensure accurate prediction of responses to unseen genetic perturbations. In our shared decoder framework, models could recover aspects of responses to new combinations of previously observed perturbation targets, yet improvements in squared error did not consistently translate into lower genome-wide absolute error. For entirely unseen targets, the evaluated methods failed to outperform a zero-change baseline in absolute error. These findings should not be interpreted as a general inability of scFMs to support perturbation prediction, because the decoder represented unseen targets using untrained condition vectors without external target-specific information. Instead, they highlight a fundamental distinction between learning representations of observed cellular variation and learning how genetic interventions alter that variation. Bridging this distinction may require more explicit modeling of perturbation targets and their biological relationships, a direction that should be tested through controlled comparisons with specialized approaches^28–32^.

Reliable evaluation of single-cell foundation models requires more than stable rankings. A model can consistently occupy a leading position without offering a substantial or robust improvement over appropriately fitted classical baselines. This distinction becomes particularly important when the independence of benchmark datasets from pretraining corpora cannot be established. Public single-cell resources are widely reused, yet checkpoint-specific training manifests often lack the sample-level information needed to determine whether evaluation studies were included during pretraining. Our provenance audit illustrates the extent of this uncertainty, while the exclusion of two human cohorts with confirmed sample or donor identifier matches demonstrates how substantially performance margins can change even when model rankings remain largely intact (Supplemental Tables S27–S29). Because excluding these cohorts also changes the composition of the benchmark, the reduced margins cannot be attributed causally to pretraining overlap. Nevertheless, the analysis exposes an important limitation of leaderboard-based evaluation: ranking stability alone cannot establish the independence, magnitude, or generalizability of a reported advantage. More credible benchmarks therefore need to combine transparent pretraining provenance with strong dataset-fitted baselines, raw performance differences, and sensitivity analyses that reveal how conclusions depend on the evaluation data.

Together, these findings suggest that progress in single-cell foundation modeling should be assessed by the biological utility and reliability of learned representations rather than by model scale or leaderboard position alone. For model users, this means selecting checkpoints according to the biological questions of interest and comparing them with classical methods trained under the same data restrictions. For benchmark developers, it means evaluating complementary biological properties, distinguishing donor mixing from biological conservation, and using meaningful controls such as the zero-change predictor in perturbation studies. For model developers, it calls for greater attention to what pretraining objectives and input representations encourage models to preserve, alongside controlled studies of backbone adaptation, cross-species transfer, and perturbation-target information. Transparent, privacy-conscious documentation of pretraining datasets is equally important for establishing credible evidence of generalization. Ultimately, the goal should not be to develop a single foundation model that leads every benchmark, but to build and evaluate representations whose biological strengths, limitations, and conditions of successful transfer are clearly understood.

### Limitations of the study

CellBench was designed to assess the out-of-the-box utility of released scFM checkpoints using frozen backbones and standardized downstream heads. This approach enables comparisons under a consistent evaluation protocol, but it does not establish whether parameter-efficient adaptation, such as LoRA^50^, or full end-to-end fine-tuning would change their relative performance against training-split-fitted PCA and scVI. Our benchmark also excluded task-specialized atlas tools such as SCimilarity^51^ and scTab^52^. Furthermore, the findings reflect the specific checkpoints and embedding extraction procedures evaluated, including Tabula’s released pancreas checkpoint^18^, mean pooling of scLong’s decoded gene representations^11^, local execution ports for Geneformer and scGPT, and scVI training without batch covariates. Because runtime and GPU memory usage were not evaluated on identical dedicated hardware, our comparisons do not support conclusions about computational efficiency.

An important limitation concerns the interpretation of perturbation prediction results. We paired each frozen control-cell representation with a shared residual lookup decoder that did not incorporate external Gene Ontology or protein-language priors for perturbation targets. Targets unseen during decoder training were represented by untrained condition vectors sampled from *N* (0, 1). Consequently, the failure to outperform the zero-change baseline in genome-wide absolute error on unseen targets applies specifically to this decoder setting. It should not be generalized to scFM-based perturbation prediction in other settings or to specialized perturbation architectures^30–32^.

Additional analytical limitations affect the interpretation of our diagnostic analyses. The base direct linear probes were evaluated on a single 1,000-cell subsample from each of four selected cohorts (HL, MS, Ci9, and MH) using seed 42 (Supplemental Table S22), where some rare cell populations contained fewer than five sampled cells. Although six sensitivity regimes spanning 10 repeated subsamples and 3,000-cell donor-grouped cross-validation confirmed the rare-class ordering, these probes operated on cached embeddings without refitting PCA, UMAP, or scVI within each cross-validation fold, so they do not provide fully cross-validated estimates for the complete representation pipelines. The pretraining provenance audit was also constrained by incomplete documentation. Because 74.0% of checkpoint and dataset pairs lacked sample-level training manifests, we could not establish whether evaluation cohorts without identified matches were independent of pretraining data.

## METHODS

### Benchmark datasets and preprocessing

To evaluate representation quality across healthy organs, disease conditions, and species, we organized the static benchmark into three biological panels, each containing four cohorts (Supplemental Table S4). The healthy human-organ panel included bone marrow mononuclear cells (BM)^38^, the Human Lung Cell Atlas (HL)^1^, adult heart (AH)^39^, and adult kidney (AK)^40^. The human-disease panel included multiple sclerosis brain (MS)^41^, ulcerative colitis epithelium (UC)^42^, lethal COVID-19 lung (C19)^43^, and triple-negative breast cancer (BC)^44^. The mouseorgan panel included four anatomically matched tissues from Tabula Muris Senis, comprising mouse bone marrow (MM), mouse lung (ML), mouse heart (MH), and mouse kidney (MK). The MH cohort combined droplet-based 10x Genomics and plate-based Smart-seq2 profiles^2^. During standardized preprocessing, cohorts containing more than 30,000 cells were limited to 30,000 cells, and all preserved cells were used in subsequent evaluations without further runtime subsampling. Separately, four CRISPR Perturb-seq screens, comprising Norman^45^, Adamson^46^, Replogle K562, and Replogle RPE1^47^, formed an independent fourth panel used exclusively for genetic perturbation response prediction.

### Model configurations and baseline implementation

All 13 pretrained scFM checkpoints were evaluated with their backbone weights frozen, using standardized model-specific adapters for input preprocessing and cell-embedding extraction (Supplemental Table S3). Rank-ordered and expression-binned sequence encoders included Geneformer with version 1 rank-value tokenization and a 2,048-token context^5^, LangCell with a 12-layer Geneformer-derived cell BERT and its pretrained 256-dimensional projection head^17^, and Nicheformer with a 1,500-token context^10^. This group also included the knowledge-informed GeneCompass^12^ and the whole-human scGPT checkpoint using 51 expression bins and a 1,200-token context^6^. Encoders incorporating protein-derived or graph-based gene information included the TranscriptFormer TF-Sapiens checkpoint with a 1,023-token context and ESM-2 protein features^9,23^, UCE with a four-layer transformer and frozen ESM-2 protein embeddings^8,23^, and scPRINT2 with protein-derived gene representations^15,16^. We also evaluated scLong, which combines dual Performer encoders with Gene Ontology graph convolutions and uses mean pooling over decoded gene representations to generate cell-level embeddings^11,53,54^. The remaining models included scFoundation, which processes expression across 19,264 genes with explicit read-depth conditioning^7^, alongside CellFM^13^, CellPLM^14^, and the tabular foundation model Tabula, evaluated using its released pancreas checkpoint^18^. For mouse cohorts, we used modelnative cross-species gene vocabularies for UCE, scPRINT2, Nicheformer, and GeneCompass. Checkpoints with human-specific gene vocabularies instead received inputs mapped using fixed one-to-one mouse-to-human orthology tables, which retained 80.4% of genes and 75.2% to 87.5% of total expression across the four mouse tissues (Supplemental Table S3).

To prevent information leakage from held-out donors into the three dataset-fitted baselines, up to 3, 000 HVGs were selected using only the training split for each seed, following library-size normalization to 10^4^ counts per cell and log(1 + *x*) transformation in Scanpy^55^. PCA fitted a 256-component linear projection on training cells^25^. UMAP learned a 256-dimensional embedding using 15 nearest neighbors on training cells and transformed validation and test cells out of sample^26^. Baseline scVI trained a two-layer negative-binomial variational autoencoder with a 256-dimensional latent space on raw training-split counts, without batch covariates, for up to 400 epochs before encoding held-out donor cells^27^. For parameter auditing, we counted unique registered parameter elements in the embedding-producing backbones, including both trainable and frozen weights. These included UCE’s 744.8M frozen ESM-2 protein lookup parameters and scPRINT2’s 85.8M frozen gene-encoder parameters. Non-parameter module buffers, such as GeneCompass’s 155.4M regulatory-prior elements, were not included in parameter counts. Task-specific classification heads not used for embedding extraction were also excluded from the audited totals (Supplemental Table S3).

### Experimental setup and evaluation metrics

All evaluations were repeated across random seeds 41, 42, and 43. For supervised cell-type annotation and expression reconstruction, donors were assigned to disjoint training (70%), validation (10%), and test (20%) splits across all 12 static cohorts (Supplemental Table S1). Because donor-level splitting can leave some rare cell types absent from the training set, closed-set annotation was restricted to labels represented among training donors. Accordingly, validation and test cells with labels absent from the corresponding training split were excluded from annotation evaluation. These exclusions affected 84 and 42 BM cells in seeds 41 and 42, respectively, 4 ML cells in seed 42, and 128 and 3 MK cells in seeds 42 and 43, respectively. No cells were excluded for this reason in any other cohort or seed. Before supervised annotation, batchsilhouette scoring, and expression reconstruction, extracted cell embeddings were standardized using *z*-score means and standard deviations calculated exclusively from the training split and applied unchanged to validation and test cells^25^.

All 16 representations, comprising 13 scFMs and three dataset-fitted baselines, were evaluated using consistent downstream head architectures to minimize differences attributable to head design (Supplemental Table S2). Supervised cell-type annotation used a three-layer MLP with two hidden layers of 256 units each and a *C*-class output layer, incorporating batch normalization, ReLU activations, dropout of 0.5, and *L*_2_-normalized output logits. Annotation heads were trained with Adam at a learning rate of 10*^−^*^3^ and a batch size of 64 for up to 500 epochs, retaining the checkpoint with minimum validation cross-entropy. Supervised HVG reconstruction predicted log-normalized expression, computed as log(1 + CP10k) (library-size normalization to 10^4^ counts per cell followed by log(1 + *x*)), over up to 400 HVGs selected strictly from the training split. It used a four-layer MLP with three hidden layers of 512, 1,024, and 512 units (batch normalization, GELU activations, dropout of 0.5) and a nonnegative ReLU output layer over the selected HVGs. Reconstruction heads were trained with Adam at a learning rate of 2 *×* 10*^−^*^3^ and an adaptive mini-batch size of 256 to 2,048 for up to 150 epochs under MSE loss, checking validation loss every 5 epochs with early stopping after four consecutive checks without improvement. For unsupervised clustering and native batch assessment, embeddings were reduced by PCA to a maximum of 256 principal components, subject to the number of cells and the original embedding dimension, before constructing 15-nearest-neighbor graphs in Scanpy^55^. Louvain community detection was performed at a resolution of 1.0 for unsupervised clustering^35^, without post-hoc batch alignment.

Each of the four static tasks was evaluated using four primary metrics, with complete mathematical definitions provided in the Supplementary Information (Supplemental Methods). Unsupervised clustering was assessed using unsupervised adjusted Rand index (uARI)^56^, normalized mutual information (NMI)^57^, scaled cell-type average silhouette width (ASW)^58^, and Hungarian optimal-matching accuracy^59^. Native batch assessment used scaled integration local inverse Simpson’s index (iLISI)^48^, within-cell-type kBET acceptance^49^, graph connectivity^37^, and scaled cell-type local inverse Simpson’s index (cLISI)^37,48^. Supervised annotation was evaluated using macro-F1, overall accuracy, balanced accuracy, and macro-precision^25^, with macroaveraged area under the precisionrecall curve (macro-AUPRC) additionally reported in Supplemental Table S20. Expression reconstruction was evaluated on log-normalized expression (log(1 + CP10k)) using coverage-penalized per-cell Pearson *r* and Spearman *ρ* across HVGs^60^, expression mean absolute error (MAE), and non-zero top-50 expressed-HVG overlap, defined as *|T_c_ ∩ P_c_|/* min(50*, |T_c_|*) over observed expressed HVGs (*t >* 0) and positive predictions (*p >* 10*^−^*^8^). Across-cell gene-wise Pearson correlation (*r*_gene_) was additionally computed in Supplemental Table S20, and cross-model correlations between evaluation metrics are reported in Supplemental Table S23.

### Perturbation prediction framework and evaluation

To prevent leakage of post-intervention expression into model inputs, each perturbed cell was randomly paired with an observed unperturbed control cell from the same cell-type stratum, using a fixed matching seed of 42. The pretrained backbone encoded only the matched control cell. Perturbation targets were represented by a trainable 128-dimensional gene lookup table, with two-gene combinations encoded as the mean of their constituent gene vectors. For heldout single-gene targets, the corresponding condition vectors were initialized from *N* (0, 1) and remained untrained. Unstandardized control-cell embeddings were concatenated with perturbation condition vectors and passed to a shared residual MLP with three hidden layers of 512, 1,024, and 512 units, using batch normalization, GELU activations, and dropout of 0.5. The decoder predicted post-perturbation expression across all *G* genes by adding the predicted residual to the matched control-cell expression and applying a nonnegative ReLU activation (Equation 1):

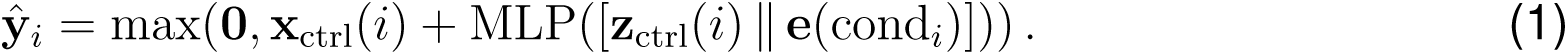

Decoders were trained across all *G* genes using Adam at a learning rate of 10*^−^*^3^ and a batch size of 64 for up to 500 epochs. Validation MSE was evaluated every 20 epochs, with early stopping after five consecutive checks without improvement. On Norman, training included all 105 constituent single-gene perturbations, while held-out two-gene combinations were used for validation and testing. On Adamson, Replogle K562, and Replogle RPE1, 10% and 20% of single-gene perturbation targets were reserved for validation and testing, respectively.

Perturbation responses were evaluated separately for each perturbation condition within each cell type. Mean predicted and observed expression changes were compared relative to the mean expression of unperturbed control cells of the same cell type. The four primary metrics were across-gene Pearson and Spearman correlations of expression changes (Δ), genome-wide MAE_Δ_, and differential-expression overlap score (DES). We additionally assessed genome-wide MAE_Δ_and MSE_Δ_ skill relative to a zero-change predictor that assumes no perturbation-induced expression shift. Positive skill indicates lower prediction error than this baseline. For reference, we also evaluated a training-mean shift baseline across all four screens, a single-gene additive baseline summing constituent single-gene training shifts on Norman held-out combinations, and seed-41 MSE_Δ_ skill restricted to the top 20 and top 100 differentially expressed genes (DE-20 and DE-100). Complete metric definitions and scoring conventions are provided in the Supplementary Information (Supplemental Methods), with screen-specific holdout information and zero-change skill results reported in Supplemental Table S25.

### Direct linear probes of cellular programs and rare cell types

To assess the linear accessibility of biological information in frozen embeddings beyond nonlinear downstream prediction, we evaluated cellular-program and rare-cell-type probes (Supplemental Table S22). At seed 42, we sampled 1,000 cells from each of the HL, MS, C19, and MH cohorts and aligned them with cached embeddings from all 16 methods, yielding 64 cohort–method evaluations. The dataset-fitted baseline representations were not refitted within probe crossvalidation folds.

Continuous targets included library depth, ribosomal transcript fraction, and interferon signaling across all four cohorts. Library depth was defined as log_10_(UMI + 1), while expression-based program scores were derived from profiles normalized to 10^4^ counts per cell and transformed using log(1+*x*). Ribosomal markers comprised *RPL\** and *RPS\** genes, while interferon markers included *ISG15*, *IFIT1*, *IFIT3*, *MX1*, *STAT1*, *OAS1*, *IRF7*, *B2M*, *HLA-A*, and their mouse orthologs, including *H2-K1*. For HL and C19, we additionally evaluated proliferation using *MKI67*, *TOP2A*, *PCNA*, *MCM2*, *CDK1*, *CCNB1*, and *TYMS*, together with mitochondrial transcript fraction based on *MT-\** genes. To test whether interferon decodability was confounded by constitutive transcripts or sequencing covariates, we additionally scored a canonical interferon-stimulated gene (ISG) target excluding B2M, HLA-A, and mouse H2-K1 and regressed out log library depth and ribosomal transcript fraction within each cohort.

Linear decodability was assessed using Ridge regression with *α* = 10 and five-fold crossvalidation, applying *z*-score standardization within each training fold. Regression used donorgrouped folds where possible and cell-stratified folds when fewer than five donors were available. Performance was summarized by pooled out-of-fold *R*^2^. Separately, five-fold cell-stratified Ridge classification with the same regularization and standardization procedure assessed rarecell-type discrimination. Out-of-fold classification F1 was averaged separately across 11 rare cell types with cohort frequencies below 1% and 26 abundant cell types with frequencies of at least 5%. Rare classes with fewer than five sampled cells were preserved, although their representation in every fold could not be guaranteed. To test sensitivity to embedding dimensionality, regularization, subsampling, and donor structure, rare-class probes were additionally repeated across six control regimes spanning native versus PCA-192-matched embeddings, nested RidgeCV regularization, 10 repeated subsamples, and donor-grouped five-fold crossvalidation on 3,000 cells per cohort, alongside direct 192-dimensional PCA on log-normalized HVGs (Supplemental Table S22). Complete scoring definitions are provided in the Supplementary Information (Supplemental Methods).

### Rank aggregation, robustness analyses, and pretraining provenance

Within each dataset, seed, and metric evaluation block, all 16 methods were ranked from 1 (best) to 16 (worst), using average ranks for exact ties and orienting error metrics so that lower values received better ranks. Ranks were averaged with equal weight across three seeds and four primary metrics per dataset, four datasets per biological panel, and all 12 cohorts for cross-panel summaries. Ranking robustness was assessed using 48 leave-one-dataset-out (LODO) and 48 leave-one-metric-out (LOMO) analyses across the 12 static task–panel combinations, together with 4 LODO and 4 LOMO analyses across the perturbation panel, excluding one dataset or one metric at a time, respectively (Supplemental Table S28).

Associations between model attributes and downstream performance were assessed using Spearman rank correlation^60^. Audited parameter counts were compared with task mean ranks across all 13 scFMs, while reported pretraining corpus sizes were evaluated across the 10 scFMs with available estimates. We also examined the association between extracted embedding dimension *d* and supervised annotation macro-F1 across all 13 scFMs. This analysis was repeated after excluding three models with collapsed representations to assess whether the association remained consistent. Leave-one-model-out analyses assessed the sensitivity of these associations to individual checkpoints, without relying on parametric significance tests (Supplemental Table S26).

We examined potential pretraining data overlap between each of the 13 scFM checkpoints and all 16 evaluation datasets, covering 208 modeldataset combinations. The audit used primary publications, supplemental sample manifests, and public repository metadata (Supplemental Table S27). To assess sensitivity to confirmed study overlap, human-organ annotation rankings were recalculated after excluding AH, and human-disease annotation rankings were recalculated after excluding MS, the two cohorts with matching sample or donor identifiers in scFoundation’s pretraining records (Supplemental Table S29).

## RESOURCE AVAILABILITY

### Lead contact

Requests for further information and resources should be directed to and will be fulfilled by the lead contact, Zelin Zang.

### Materials availability

This study did not generate new unique reagents.

### Data and code availability

All 16 evaluation datasets were obtained from publicly accessible repositories, including GEO, CZ CELLxGENE Discover^3^, the Broad Single Cell Portal, and original Perturb-seq data releases^45–47^. Source collection URLs and prepared matrix dimensions are provided in Supplemental Table S4. The benchmark code, frozen-backbone model adapters, dataset and checkpoint download manifests, diagnostic analysis scripts, run-level evaluation results, parameter and pretraining-provenance audits, and supplemental workbooks are publicly available at https://github.com/BiologyNet/CellBench-scFM. The repository also provides instructions for obtaining the original datasets and pretrained model checkpoints from their respective sources. Additional information required to reanalyze the results reported in this study is available from the lead contact upon reasonable request.

## ACKNOWLEDGMENTS

This work was supported by the National Science and Technology Major Project of China (grant no. 2022ZD0115100), the National Natural Science Foundation of China (grant no. U21A20427), and the Center of Synthetic Biology and Integrated Bioengineering of Westlake University (grant no. WU2022A009). We thank the Westlake University HPC Center for computational support.

## AUTHOR CONTRIBUTIONS

Conceptualization, Z.Z. and F.H.; methodology, Y.X., Y.L., and Y.Y.; data curation, Y.L., Q.X., and H.T.; investigation, Y.X., Y.L., and Y.Y.; writing–original draft, Y.X., Y.L., and Y.Y.; writing–review & editing, all authors.

## DECLARATION OF GENERATIVE AI AND AI-ASSISTED TECH-NOLOGIES

During the preparation of this work, the authors used Gemini, Claude, and Codex to assist with manuscript outline structuring, cross-table numerical verification, Cell Press *Patterns* formatting checks, and English prose editing. After using these tools, the authors reviewed, verified against raw benchmark exports, and edited the content as needed and take full responsibility for the content of the publication.

## SUPPLEMENTARY INFORMATION INDEX

Supplementary Information. Supplemental Methods, Figures S1–S21, and Tables S1–S29

## References

[1] Sikkema, L., Ramírez-Suástegui, C., Strobl, D.C., et al. (2023). An integrated cell atlas of the lung in health and disease. Nat. Med. 29, 1563–1577. 10.1038/s41591-023-02327-2.

[2] The Tabula Muris Consortium (2020). A single-cell transcriptomic atlas characterizes ageing tissues in the mouse. Nature 583, 590–595. 10.1038/s41586-020-2496-1.

[3] CZI Cell Science Program, Abdulla, S., Aevermann, B., et al. (2025). CZ CELLxGENE Discover: A single-cell data platform for scalable exploration, analysis and modeling of aggregated data. Nucleic Acids Res. 53, D886–D900. 10.1093/nar/gkae1142.

[4] Yang, F., Wang, W., Wang, F., et al. (2022). scBERT as a large-scale pretrained deep language model for cell type annotation of single-cell RNA-seq data. *Nat*. Mach. Intell. 4, 852–866. 10.1038/s42256-022-00534-z.

[5] Theodoris, C.V., Xiao, L., Chopra, A., et al. (2023). Transfer learning enables predictions in network biology. Nature 618, 616–624. 10.1038/s41586-023-06139-9.

[6] Cui, H., Wang, C., Maan, H., et al. (2024). scGPT: Toward building a foundation model for single-cell multi-omics using generative AI. Nat. Methods 21, 1470–1480. 10.1038/s41592-024-02201-0.

[7] Hao, M., Gong, J., Zeng, X., et al. (2024). Large-scale foundation model on single-cell transcriptomics. Nat. Methods 21, 1481–1491. 10.1038/s41592-024-02305-7.

[8] Rosen, Y., Roohani, Y., et al. (2026). Universal cell embedding provides a foundation model for cell biology. Nature 656, 183–191. 10.1038/s41586-026-10689-z.

[9] Pearce, J.D., Simmonds, S.E., Mahmoudabadi, G., et al. (2026). TranscriptFormer: A generative cell atlas across 1.5 billion years of evolution. Science 393, eaec8514. 10.1126/science.aec8514.

[10] Tejada-Lapuerta, A., Schaar, A.C., et al. (2025). Nicheformer: A foundation model for singlecell and spatial omics. Nat. Methods 22, 2525–2538. 10.1038/s41592-025-02814-z.

[11] Bai, D., et al. (2026). scLong: A billion-parameter foundation model for capturing long-range gene context in single-cell transcriptomics. Nat. Commun. 17, 2380. 10.1038/s41467-026-69102-y.

[12] Yang, X., Liu, G., Feng, G., et al. (2024). GeneCompass: Deciphering universal gene regulatory mechanisms with a knowledge-informed cross-species foundation model. Cell Res. 34, 830–845. 10.1038/s41422-024-01034-y.

[13] Zeng, Y., Xie, J., Shangguan, N., et al. (2025). CellFM: A large-scale foundation model pre-trained on transcriptomics of 100 million human cells. Nat. Commun. 16, 4679. 10.1038/s41467-025-59926-5.

[14] Wen, H., Tang, W., Dai, X., et al. (2024). CellPLM: Pre-training of cell language model beyond single cells. In International Conference on Learning Representations (ICLR). https://proceedings.iclr.cc/paper_files/paper/2024/hash/173bcdbd820edbe43284e0941109ada9-Abstract-Conference.html.

[15] Kalfon, J., Samaran, J., Peyré, G., and Cantini, L. (2025). scPRINT: Pre-training on 50 million cells allows robust gene network predictions. Nat. Commun. 16, 3607. 10.1038/s41467-025-58699-1.

[16] Kalfon, J., Peyré, G., and Cantini, L. (2025). scPRINT-2: Towards the next-generation of cell foundation models and benchmarks. Preprint at bioRxiv, 2025.12.11.693702. 10.64898/2025.12.11.693702.

[17] Zhao, S., Zhang, J., Wu, Y., Luo, Y., and Nie, Z. (2024). LangCell: Language-cell pre-training for cell identity understanding. In Proceedings of the 41st International Conference on Machine Learning (ICML), PMLR 235, 61159–61185. https://proceedings.mlr.press/v235/zhao24u.html.

[18] Ding, J., Lin, J., Jiang, S., et al. (2025). Tabula: A tabular self-supervised foundation model for single-cell transcriptomics. In Advances in Neural Information Processing Systems (NeurIPS). https://papers.nips.cc/paper_files/paper/2025/hash/95d590995a8722259c61e094b62b25ac-Abstract-Conference.html.

[19] Qiu, P., Chen, Q., Qin, H., et al. (2025). BioLLM: A standardized framework for integrating and benchmarking single-cell foundation models. Patterns 6, 101326. 10.1016/j.patter.2025.101326.

[20] Wu, J., Ye, Q., Wang, Y., et al. (2025). Biology-driven insights into the power of single-cell foundation models. Genome Biol. 26, 334. 10.1186/s13059-025-03781-6.

[21] Kędzierska, K.Z., Crawford, L., Amini, A.P., and Lu, A.X. (2025). Zero-shot evaluation reveals limitations of single-cell foundation models. Genome Biol. 26, 101. 10.1186/s13059-025-03574-x.

[22] Boiarsky, R., Singh, N.M., Buendia, A., Getz, G., and Sontag, D. (2023). A deep dive into single-cell RNA sequencing foundation models. Preprint at bioRxiv, 2023.10.19.563100. 10.1101/2023.10.19.563100.

[23] Lin, Z., Akin, H., Rao, R., et al. (2023). Evolutionary-scale prediction of atomic-level protein structure with a language model. Science 379, 1123–1130. 10.1126/science.ade2574.

[24] Chen, Y., and Zou, J. (2025). Simple and effective embedding model for single-cell biology built from ChatGPT. *Nat*. Biomed. Eng. 9, 483–493. 10.1038/s41551-024-01284-6.

[25] Pedregosa, F., Varoquaux, G., Gramfort, A., et al. (2011). Scikit-learn: Machine learning in Python. J. Mach. Learn. Res. 12, 2825–2830. https://www.jmlr.org/papers/v12/pedregosa11a.html.

[26] McInnes, L., Healy, J., and Melville, J. (2018). UMAP: Uniform Manifold Approximation and Projection for dimension reduction. J. Open Source Softw. 3, 861. 10.21105/joss.00861.

[27] Lopez, R., Regier, J., Cole, M.B., Jordan, M.I., and Yosef, N. (2018). Deep generative modeling for single-cell transcriptomics. Nat. Methods 15, 1053–1058. 10.1038/s41592-018-0229-2.

[28] Ahlmann-Eltze, C., Huber, W., and Anders, S. (2025). Deep-learning-based gene perturbation effect prediction does not yet outperform simple linear baselines. Nat. Methods 22, 1657–1661. 10.1038/s41592-025-02772-6.

[29] Miller, H.E., Mejia, G.M., Leblanc, F.J.A., Swain, B., Wang, B., and de Lima Camillo, L.P. (2026). Deep learning perturbation models can outperform baselines on calibrated metrics. Nat. Biotechnol. 10.1038/s41587-026-03307-w.

[30] Roohani, Y., Huang, K., and Leskovec, J. (2024). Predicting transcriptional outcomes of novel multigene perturbations with GEARS. Nat. Biotechnol. 42, 927–935. 10.1038/s41587-023-01905-6.

[31] Lotfollahi, M., Wolf, F.A., and Theis, F.J. (2019). scGen predicts single-cell perturbation responses. Nat. Methods 16, 715–721. 10.1038/s41592-019-0494-8.

[32] Lotfollahi, M., Klimovskaia Susmelj, A., De Donno, C., et al. (2023). Predicting cellular responses to complex perturbations in high-throughput screens. Mol. Syst. Biol. 19, e11517. 10.15252/msb.202211517.

[33] Haber, E., Alam, S., Ho, N., et al. (2025). HEIMDALL: Disentangling tokenizer design for robust transfer in single-cell foundation models. Preprint at bioRxiv, 2025.11.09.687403v2. 10.1101/2025.11.09.687403.

[34] DenAdel, A., Hughes, M., Thoutam, A., et al. (2026). Evaluating the role of pretraining dataset size and diversity on single-cell foundation model performance. Nat. Methods 23, 1447–1457. 10.1038/s41592-026-03120-y.

[35] Blondel, V.D., Guillaume, J.-L., Lambiotte, R., and Lefebvre, E. (2008). Fast unfolding of communities in large networks. J. Stat. Mech. 2008, P10008. 10.1088/1742-5468/2008/10/P10008.

[36] Traag, V.A., Waltman, L., and van Eck, N.J. (2019). From Louvain to Leiden: Guaranteeing well-connected communities. Sci. Rep. 9, 5233. 10.1038/s41598-019-41695-z.

[37] Luecken, M.D., Büttner, M., Chaichoompu, K., et al. (2022). Benchmarking atlas-level data integration in single-cell genomics. Nat. Methods 19, 41–50. 10.1038/s41592-021-01336-8.

[38] Luecken, M.D., Burkhardt, D.B., Cannoodt, R., et al. (2021). A sandbox for prediction and integration of DNA, RNA, and proteins in single cells. In NeurIPS Datasets and Benchmarks. https://datasets-benchmarks-proceedings.neurips.cc/paper/2021/hash/158f3069a435b314a80bdcb024f8e422-Abstract-round2.html.

[39] Litviňuková, M., Talavera-López, C., Maatz, H., et al. (2020). Cells of the adult human heart. Nature 588, 466–472. 10.1038/s41586-020-2797-4.

[40] Lake, B.B., Menon, R., Winfree, S., et al. (2023). An atlas of healthy and injured cell states and niches in the human kidney. Nature 619, 585–594. 10.1038/s41586-023-05769-3.

[41] Jäkel, S., Agirre, E., Mendanha Falcão, A., et al. (2019). Altered human oligodendrocyte heterogeneity in multiple sclerosis. Nature 566, 543–547. 10.1038/s41586-019-0903-2.

[42] Smillie, C.S., Biton, M., Ordovas-Montanes, J., et al. (2019). Intra- and inter-cellular rewiring of the human colon during ulcerative colitis. Cell 178, 714–730.e22. 10.1016/j.cell.2019.06.029.

[43] Melms, J.C., Biermann, J., Huang, H., et al. (2021). A molecular single-cell lung atlas of lethal COVID-19. Nature 595, 114–119. 10.1038/s41586-021-03569-1.

[44] Yan, Y., Lin, Y., Kumar, T., et al. (2026). Ecotypes of triple-negative breast cancer in response to chemotherapy. Nature 654, 1088–1097. 10.1038/s41586-026-10469-9.

[45] Norman, T.M., Horlbeck, M.A., Replogle, J.M., et al. (2019). Exploring genetic interaction manifolds constructed from rich single-cell phenotypes. Science 365, 786–793. 10.1126/science.aax4438.

[46] Adamson, B., Norman, T.M., Jost, M., et al. (2016). A multiplexed single-cell CRISPR screening platform enables systematic dissection of the unfolded protein response. Cell 167, 1867–1882.e21. 10.1016/j.cell.2016.11.048.

[47] Replogle, J.M., Saunders, R.A., Pogson, A.N., et al. (2022). Mapping informationrich genotype-phenotype landscapes with genome-scale Perturb-seq. Cell 185, 2559–2575.e28. 10.1016/j.cell.2022.05.013.

[48] Korsunsky, I., Millard, N., Fan, J., et al. (2019). Fast, sensitive and accurate integration of single-cell data with Harmony. Nat. Methods 16, 1289–1296. 10.1038/s41592-019-0619-0.

[49] Büttner, M., Miao, Z., Wolf, F.A., Teichmann, S.A., and Theis, F.J. (2019). A test metric for assessing single-cell RNA-seq batch correction. Nat. Methods 16, 43–49. 10.1038/s41592-018-0254-1.

[50] Hu, E.J., Shen, Y., Wallis, P., et al. (2022). LoRA: Low-rank adaptation of large language models. In International Conference on Learning Representations (ICLR). https://openreview.net/forum?id=nZeVKeeFYf9.

[51] Heimberg, G., Kuo, T., DePianto, D.J., et al. (2025). A cell atlas foundation model for scalable search of similar human cells. Nature 638, 1085–1094. 10.1038/s41586-024-08411-y.

[52] Fischer, F., Fischer, D.S., Mukhin, R., et al. (2024). scTab: Scaling cross-tissue singlecell annotation models. Nat. Commun. 15, 6611. 10.1038/s41467-024-51059-5.

[53] Choromanski, K., Likhosherstov, V., Dohan, D., et al. (2021). Rethinking attention with Performers. In International Conference on Learning Representations (ICLR). https://openreview.net/forum?id=Ua6zuk0WRH.

[54] Kipf, T.N., and Welling, M. (2017). Semi-supervised classification with graph convolutional networks. In International Conference on Learning Representations (ICLR). https://openreview.net/forum?id=SJU4ayYgl.

[55] Wolf, F.A., Angerer, P., and Theis, F.J. (2018). SCANPY: Large-scale single-cell gene expression data analysis. Genome Biol. 19, 15. 10.1186/s13059-017-1382-0.

[56] Hubert, L., and Arabie, P. (1985). Comparing partitions. J. Classif. 2, 193–218. 10.1007/BF01908075.

[57] Strehl, A., and Ghosh, J. (2002). Cluster ensembles—a knowledge reuse framework for combining multiple partitions. J. Mach. Learn. Res. 3, 583–617. https://www.jmlr.org/papers/v3/strehl02a.html.

[58] Rousseeuw, P.J. (1987). Silhouettes: A graphical aid to the interpretation and validation of cluster analysis. J. Comput. Appl. Math. 20, 53–65. 10.1016/0377-0427(87)90125-7.

[59] Kuhn, H.W. (1955). The Hungarian method for the assignment problem. Nav. Res. Logist. Q. 2, 83–97. 10.1002/nav.3800020109.

[60] Spearman, C. (1904). The proof and measurement of association between two things. Am. J. Psychol. 15, 72–101. 10.2307/1412159.

