## Supplementary Information for "CellBench: task-dependent performance and generalization limits of single-cell foundation models"

[1]School of Information Science & Electronic Engineering, Zhejiang University, No. 866 Yuhangtang Road, Hangzhou 310058, Zhejiang, China
[2]School of Engineering, Westlake University, No. 600 Dunyu Road, Hangzhou 310030, Zhejiang, China
[3]Yazhouwan National Laboratory, No. 8 Huanjin Road, Sanya 572025, Hainan, China
[4]Faculty of Computing, Harbin Institute of Technology, No. 92 Xidazhi Street, Harbin 150040, Heilongjiang, China
[5]Department of Public Health & Primary Care, School of Medicine, KU Leuven, No. 49 Herestraat Street, 3000 Leuven, Belgium
[6]School of Information Science & Technology, Zhejiang Shuren University, No. 8 Shuren Street, Hangzhou 310015, Zhejiang, China
[7]These authors contributed equally
[8]Lead contact

### SUPPLEMENTARY INFORMATION INDEX

This Supplementary Information contains Supplemental Methods (evaluation metrics and ranking conventions), Figures S1–S21, and Tables S1–S29. Complete machine-readable workbooks are publicly available at `https://github.com/BiologyNet/CellBench-scFM`.


| Display items | Section / group | Contents and alignment with main text | Page |
|---|---|---|---|
| Figures S1–S4 | Unsupervised clustering | Matches Figure 2: uARI (Figure S1), NMI (Figure S2), scaled ASW (Figure S3), and matching accuracy (Figure S4). | 11–14 |
| Figures S5–S8 | Native batch assessment | Matches Figure 3: scaled iLISI (Figure S5), kBET (Figure S6), scaled cLISI (Figure S7), and graph connectivity (Figure S8). | 15–18 |
| Figures S9–S12 | Cell-type annotation | Matches Figure 4: macro-F1 (Figure S9), accuracy (Figure S10), balanced accuracy (Figure S11), and macro-precision (Figure S12). | 19–22 |
| Figures S13–S16 | HVG reconstruction | Matches Figure 5 on $\log(1+\mathrm{CP10k})$: Pearson $r$ (Figure S13), Spearman $\rho$ (Figure S14), MAE (Figure S15), and non-zero top-50 overlap (Figure S16). | 23–26 |
| Figure S17 | Perturbation prediction | Matches Figure 6: Pearson $\Delta$, Spearman $\Delta$, MAE $\Delta$, and DES across Norman, Adamson, Replogle K562, and Replogle RPE1. | 27 |
| Figures S18–S19 | Model scale | Parameter count versus task rank (Figure S18) and reported corpus size versus task rank (Figure S19). | 28–29 |
| Figures S20–S21 | Rank stability | Seed-to-seed rank range (Figure S20) and pairwise metric-rank agreement (Figure S21). | 30–31 |

### SUPPLEMENTAL METHODS: EVALUATION METRICS

This section provides the mathematical definitions, normalization bounds, edge-case rules, and ranking orientations (↑ higher is better; ↓ lower is better) for all primary and diagnostic metrics evaluated in the main text and Supplementary Information. Subsections and individual metrics are ordered to match their sequence of appearance in the main-text Results: unsupervised clustering (Figure 2), native batch assessment (Figure 3), supervised cell-type annotation (Figure 4), supervised HVG expression reconstruction (Figure 5), direct linear probes (Table S22), genetic perturbation prediction and reference-baseline diagnostics (Figure 6 and Table S25), and composite ranking with cross-model rank correlations (Figure 1d and Tables S23, S26, and S28). Throughout, $i$ indexes cells, $j$ indexes genes, $c$ indexes cell types, and $b$ indexes donors or batches.

#### 1. Unsupervised clustering metrics

Let $\tilde{\mathbf{Z}} \in \mathbb{R}^{N \times d'}$ denote full-cohort cell embeddings projected onto $d' = \min(256, N, d)$ principal components across $N$ cells with ground-truth cell-type labels $\mathbf{y} = (y_1, \ldots, y_N) \in \{1, \ldots, C\}^N$. Louvain community detection (`sc.tl.louvain`) at fixed resolution $1.0$ over the $k$-nearest-neighbor graph ($k = \min(15, N-1)$) produces discrete cluster assignments $\hat{\mathbf{c}} = (\hat{c}_1, \ldots, \hat{c}_N) \in \{1, \ldots, K\}^N$. Verification with Leiden clustering at the same resolution and with held-out donor splits produced nearly identical 12-cohort rankings ($\rho = 0.994$ and $\rho = 0.977$, respectively). Let $n_{ab} = \sum_{i=1}^{N} \mathbb{I}(y_i = a,\, \hat{c}_i = b)$ denote the contingency count between cell type $a$ and cluster $b$, with row totals $u_a = \sum_{b=1}^{K} n_{ab}$ and column totals $v_b = \sum_{a=1}^{C} n_{ab}$. The first three metrics measure agreement between $\hat{\mathbf{c}}$ and $\mathbf{y}$ from this contingency table, whereas the fourth evaluates continuous cell-type separation directly in $\tilde{\mathbf{Z}}$.

**Unsupervised adjusted Rand index (uARI; range $[-1, 1]$, ↑).** Chance-corrected pairwise partition agreement is computed via the Hubert–Arabie formulation:

$$\mathrm{uARI} = \frac{\sum_{a=1}^{C} \sum_{b=1}^{K} \binom{n_{ab}}{2} - \left[\sum_{a=1}^{C} \binom{u_a}{2} \sum_{b=1}^{K} \binom{v_b}{2}\right] \Big/ \binom{N}{2}}{\frac{1}{2}\left[\sum_{a=1}^{C} \binom{u_a}{2} + \sum_{b=1}^{K} \binom{v_b}{2}\right] - \left[\sum_{a=1}^{C} \binom{u_a}{2} \sum_{b=1}^{K} \binom{v_b}{2}\right] \Big/ \binom{N}{2}}, \tag{S1}$$

which equals $0$ in expectation under random label permutations and $1$ for identical partitions.

**Normalized mutual information (NMI; range $[0, 1]$, ↑).** Using $0 \log 0 \equiv 0$ and setting $\mathrm{NMI} = 1$ when $H(\mathbf{y}) = H(\hat{\mathbf{c}}) = 0$, mutual information normalized by arithmetic-mean Shannon entropy is

$$\mathrm{NMI}(\mathbf{y}, \hat{\mathbf{c}}) = \frac{2\, I(\mathbf{y}; \hat{\mathbf{c}})}{H(\mathbf{y}) + H(\hat{\mathbf{c}})}, \tag{S2}$$

where the mutual information and marginal entropies are

$$I(\mathbf{y}; \hat{\mathbf{c}}) = \sum_{a=1}^{C} \sum_{b=1}^{K} \frac{n_{ab}}{N} \log \frac{N\, n_{ab}}{u_a v_b}, \qquad H(\mathbf{y}) = -\sum_{a=1}^{C} \frac{u_a}{N} \log \frac{u_a}{N}, \qquad H(\hat{\mathbf{c}}) = -\sum_{b=1}^{K} \frac{v_b}{N} \log \frac{v_b}{N}. \tag{S3}$$

**Hungarian optimal-matching accuracy (ACC; range $[0, 1]$, ↑).** Solving the maximum-weight one-to-one bipartite assignment $\mathcal{M} \subset \{1, \ldots, C\} \times \{1, \ldots, K\}$ with $|\mathcal{M}| = \min(C, K)$ on the contingency matrix via the Kuhn–Munkres algorithm yields

$$\mathrm{ACC}_{\mathrm{cluster}} = \frac{1}{N} \max_{\mathcal{M}} \sum_{(a,b) \in \mathcal{M}} n_{ab}. \tag{S4}$$

**Scaled cell-type average silhouette width (Scaled ASW; range $[0, 1]$, ↑).** For each cell $i$ with label $y_i$ ($1 < C < N$), let $a(i)$ be its mean Euclidean distance in $\tilde{\mathbf{Z}}$ to all other cells sharing label $y_i$, and let $b(i) = \min_{c \neq y_i} d(i, c)$ be its minimum mean Euclidean distance to cells of any other cell type $c$. The per-cell silhouette coefficient $s(i)$ (set to $0$ for singleton classes) is averaged across all $N$ cells and linearly mapped to $[0, 1]$:

$$s(i) = \frac{b(i) - a(i)}{\max\{a(i),\, b(i)\}} \in [-1, 1], \qquad \mathrm{ASW}_{\text{scaled}} = \frac{1}{2}\left(\frac{1}{N}\sum_{i=1}^{N} s(i) + 1\right). \tag{S5}$$

#### 2. Native-embedding batch-assessment metrics

Native batch assessment evaluates unintegrated projected embeddings $\tilde{\mathbf{Z}} \in \mathbb{R}^{N \times d'}$ across each cohort's batch labels $\mathbf{b} \in \{1, \ldots, B\}^N$ ($B \geq 2$; identical to donor identity in 8 of 12 cohorts, and corresponding to batch, study, source, or sample identifiers in BM, HL, AK, and UC; Table S4) and cell-type labels $\mathbf{y} \in \{1, \ldots, C\}^N$ ($C \geq 2$). Main-text Figure 3 and composite native batch rankings weight two donor-mixing metrics (scaled iLISI and kBET) equally against two clustering-free cell-type conservation metrics (scaled cLISI and graph connectivity), following `scIB`.

**Scaled integration LISI (Scaled iLISI; range $[0, 1]$, ↑ donor mixing).** On the $k_0$-nearest-neighbor graph with $k_0 = \min(90, N - 1)$, let $p(b \mid i)$ denote the Gaussian-kernel-weighted local neighborhood probability of batch $b$ around cell $i$. The local inverse Simpson index $\mathrm{iLISI}(i) \in [1, B]$ measures the effective number of mixed batches per neighborhood; taking the cell-level median and scaling linearly by $B$ yields

$$\mathrm{iLISI}(i) = \left[\sum_{b=1}^{B} p(b \mid i)^2\right]^{-1}, \qquad \mathrm{iLISI}_{\text{scaled}} = \frac{\mathrm{median}_{i=1,\ldots,N}\, \mathrm{iLISI}(i) - 1}{B - 1}. \tag{S6}$$

**Within-cell-type kBET acceptance rate (kBET; range $[0, 1]$, ↑ donor mixing).** kBET is evaluated separately within every eligible cell type $c \in \mathcal{C}_{\text{elig}}$ containing $N_c \geq 10$ cells across $B_c \geq 2$ present batches. Letting $f_{c,b} = n_{c,b}/N_c$ denote within-type batch proportions and $\bar{n}_c = N_c/B_c$ the mean per-batch count in cell type $c$, the adaptive neighborhood size is

$$k_0(c) = \max\Big(2,\ \min\big(N_c - 1,\ \min(70,\ \max(10, \lfloor \bar{n}_c/4 \rfloor))\big)\Big). \tag{S7}$$

For each cell $i$ in stratum $c$, observed batch counts $O_{i,b}$ among its $k_0(c)$ Euclidean nearest neighbors in $\tilde{\mathbf{Z}}$ (excluding $i$) are compared against expected counts $E_{c,b} = k_0(c)\, f_{c,b}$ via Pearson's $\chi^2$ test (with cumulative distribution $F_{\chi^2_{B_c-1}}$ on $B_c - 1$ degrees of freedom):

$$\chi_i^2 = \sum_{b=1}^{B_c} \frac{(O_{i,b} - E_{c,b})^2}{E_{c,b}}, \qquad r_c = \frac{1}{N_c}\sum_{i:y_i=c} \mathbb{I}\Big(1 - F_{\chi^2_{B_c-1}}(\chi_i^2) < 0.05\Big), \tag{S8}$$

$$\mathrm{kBET} = 1 - \frac{1}{|\mathcal{C}_{\text{elig}}|}\sum_{c \in \mathcal{C}_{\text{elig}}} r_c. \tag{S9}$$

**Scaled cell-type LISI (Scaled cLISI; range $[0, 1]$, ↑ cell-type conservation).** Computed on the same $k_0 = \min(90, N - 1)$ graph over local cell-type probabilities $p(c \mid i)$, where $\mathrm{cLISI}(i) \in [1, C]$ equals $1$ for pure single-cell-type neighborhoods; inverting and scaling to $[0, 1]$ gives

$$\mathrm{cLISI}(i) = \left[\sum_{c=1}^{C} p(c \mid i)^2\right]^{-1}, \qquad \mathrm{cLISI}_{\text{scaled}} = \frac{C - \mathrm{median}_{i=1,\ldots,N}\, \mathrm{cLISI}(i)}{C - 1}. \tag{S10}$$

**Graph connectivity (range $(0, 1]$, ↑ cell-type conservation).** On the $k = \min(15, N-1)$ nearest-neighbor graph $G$, let $G_c$ be the subgraph induced by cells $V_c = \{i : y_i = c\}$ of cell type $c$, and let $|\mathrm{LCC}(G_c)|$ denote the number of vertices in the largest connected component of $G_c$:

$$\text{Graph connectivity} = \frac{1}{C} \sum_{c=1}^{C} \frac{|\mathrm{LCC}(G_c)|}{|V_c|}. \tag{S11}$$

#### 3. Supervised cell-type annotation metrics

Annotation metrics are evaluated on $N_{\text{test}}$ held-out test-donor cells over the set of test-supported classes $\mathcal{C}_{\text{test}} = \{c : N_{\text{test},c} > 0\}$ ($C_{\text{test}} = |\mathcal{C}_{\text{test}}|$). Let $\hat{y}_i$ denote the predicted label for cell $i$, and let $\mathrm{TP}_c$, $\mathrm{FP}_c$, and $\mathrm{FN}_c$ denote one-vs-rest true positives, false positives, and false negatives for class $c \in \mathcal{C}_{\text{test}}$, with per-class precision $\mathrm{Prec}_c = \mathrm{TP}_c/(\mathrm{TP}_c + \mathrm{FP}_c)$ (set to $0$ if $\mathrm{TP}_c + \mathrm{FP}_c = 0$), recall $\mathrm{Rec}_c = \mathrm{TP}_c/(\mathrm{TP}_c + \mathrm{FN}_c)$, and $\mathrm{F1}_c = 2\,\mathrm{TP}_c/(2\,\mathrm{TP}_c + \mathrm{FP}_c + \mathrm{FN}_c)$ (set to $0$ if the denominator is $0$).

**Overall accuracy, macro-F1, balanced accuracy, and macro-precision (all range $[0, 1]$, ↑).** In the order reported in main-text Figure 4, the four primary annotation metrics are

$$\mathrm{Acc} = \frac{1}{N_{\text{test}}} \sum_{i=1}^{N_{\text{test}}} \mathbb{I}(\hat{y}_i = y_i), \tag{S12}$$

$$\text{Macro-F1} = \frac{1}{C_{\text{test}}} \sum_{c \in \mathcal{C}_{\text{test}}} \mathrm{F1}_c = \frac{1}{C_{\text{test}}} \sum_{c \in \mathcal{C}_{\text{test}}} \frac{2\,\mathrm{TP}_c}{2\,\mathrm{TP}_c + \mathrm{FP}_c + \mathrm{FN}_c}, \tag{S13}$$

$$\text{Balanced Acc} = \frac{1}{C_{\text{test}}} \sum_{c \in \mathcal{C}_{\text{test}}} \mathrm{Rec}_c, \tag{S14}$$

$$\mathrm{Precision}_{\text{macro}} = \frac{1}{C_{\text{test}}} \sum_{c \in \mathcal{C}_{\text{test}}} \mathrm{Prec}_c. \tag{S15}$$

**Macro-AUPRC (Table S20; range $[0, 1]$, ↑).** From normalized softmax class probabilities $\hat{p}_{i,c}$, one-vs-rest step-interpolated average precision is averaged across classes $\mathcal{C}_{\text{prob}} = \{c : 0 < N_{\text{test},c} < N_{\text{test}}\}$:

$$\mathrm{AUPRC}_{\text{macro}} = \frac{1}{|\mathcal{C}_{\text{prob}}|} \sum_{c \in \mathcal{C}_{\text{prob}}} \sum_{m} (R_{c,m} - R_{c,m-1})\, P_{c,m}, \tag{S16}$$

where $(P_{c,m}, R_{c,m})$ denote precision and recall at the $m$-th sorted probability threshold.

#### 4. Supervised HVG expression reconstruction metrics

Let $\hat{\mathbf{Y}}, \mathbf{Y} \in \mathbb{R}^{N_{\text{test}} \times M}$ denote predicted and observed expression on the $\log(1 + \mathrm{CP10k})$ scale across $N_{\text{test}}$ held-out test cells and $M \leq 400$ training-split-selected HVGs. For any two non-constant vectors $\mathbf{u}, \mathbf{v} \in \mathbb{R}^L$ with sample means $\bar{u}$ and $\bar{v}$, let

$$\mathrm{corr}(\mathbf{u}, \mathbf{v}) = \frac{\sum_{\ell=1}^{L} (u_\ell - \bar{u})(v_\ell - \bar{v})}{\sqrt{\sum_{\ell=1}^{L} (u_\ell - \bar{u})^2} \sqrt{\sum_{\ell=1}^{L} (v_\ell - \bar{v})^2}}, \tag{S17}$$

and let $\mathrm{rank}(\mathbf{u})$ map entries of $\mathbf{u}$ to their average-tie ranks. The four primary reconstruction metrics follow the order in main-text Figure 5, followed by the gene-wise diagnostic channel reported in Tables S20 and S23.

**Non-zero top-50 expressed-HVG overlap (range $[0, 1]$, ↑).** To prevent zero-expression ties from inflating overlap, let $\mathcal{V}_{\text{nz}} = \{i : n_i^{\text{obs}} = |\{j : Y_{ij} > 0\}| \geq 1\}$ denote test cells with at least one expressed target HVG. For each $i \in \mathcal{V}_{\mathrm{nz}}$, let $K_i = \min(50, n_i^{\mathrm{obs}})$, let $T_i$ be the index set of the $K_i$ highest-expressed observed HVGs ($Y_{ij} > 0$), and let $P_i$ be the index set of the $\min(K_i, |\{j : \hat{Y}_{ij} > 10^{-8}\}|)$ highest positive predicted HVGs (setting $|T_i \cap P_i| = 0$ if no predicted entry exceeds $10^{-8}$):

$$\text{Top-50 overlap} = \frac{1}{|\mathcal{V}_{\mathrm{nz}}|} \sum_{i \in \mathcal{V}_{\mathrm{nz}}} \frac{|T_i \cap P_i|}{K_i}. \tag{S18}$$

**Expression mean absolute error ($\mathrm{MAE}_{\mathrm{expr}}$; range $[0, \infty)$, $\downarrow$).** Element-wise absolute error on the $\log(1 + \mathrm{CP10k})$ scale averaged across all held-out cells and target HVGs is

$$\mathrm{MAE}_{\mathrm{expr}} = \frac{1}{N_{\mathrm{test}}\, M} \sum_{i=1}^{N_{\mathrm{test}}} \sum_{j=1}^{M} |\hat{Y}_{ij} - Y_{ij}|. \tag{S19}$$

**Coverage-penalized per-cell Pearson $r$ and Spearman $\rho$ (range $[-1, 1]$, $\uparrow$).** Evaluated across all test cells with non-constant observed profiles, $\mathcal{V}_{\mathrm{cell}}^{\mathrm{obs}} = \{i : M > 1,\ \max_j Y_{ij} > \min_j Y_{ij}\}$, assigning a penalized correlation of $0$ whenever predicted profile $\hat{\mathbf{Y}}_{i\cdot}$ is constant:

$$r_{\mathrm{expr}} = \frac{1}{|\mathcal{V}_{\mathrm{cell}}^{\mathrm{obs}}|} \sum_{i \in \mathcal{V}_{\mathrm{cell}}^{\mathrm{obs}}} \mathbb{I}(\mathrm{std}(\hat{\mathbf{Y}}_{i\cdot}) > 0)\, \mathrm{corr}(\hat{\mathbf{Y}}_{i\cdot},\, \mathbf{Y}_{i\cdot}), \tag{S20}$$

$$\rho_{\mathrm{expr}} = \frac{1}{|\mathcal{V}_{\mathrm{cell}}^{\mathrm{obs}}|} \sum_{i \in \mathcal{V}_{\mathrm{cell}}^{\mathrm{obs}}} \mathbb{I}(\mathrm{std}(\hat{\mathbf{Y}}_{i\cdot}) > 0)\, \mathrm{corr}\big(\mathrm{rank}(\hat{\mathbf{Y}}_{i\cdot}),\, \mathrm{rank}(\mathbf{Y}_{i\cdot})\big). \tag{S21}$$

**Across-cell gene-wise correlation ($r_{\mathrm{gene}}$, Tables S20 and S23; range $[-1, 1]$, $\uparrow$).** Evaluated along the cell axis across all HVG columns with non-constant observed profiles $\mathcal{V}_{\mathrm{gene}}^{\mathrm{obs}} = \{j : \max_i Y_{ij} > \min_i Y_{ij}\}$, assigning $0$ to constant predicted columns:

$$r_{\mathrm{gene}} = \frac{1}{|\mathcal{V}_{\mathrm{gene}}^{\mathrm{obs}}|} \sum_{j \in \mathcal{V}_{\mathrm{gene}}^{\mathrm{obs}}} \mathbb{I}(\mathrm{std}(\hat{\mathbf{Y}}_{\cdot j}) > 0)\, \mathrm{corr}(\hat{\mathbf{Y}}_{\cdot j},\, \mathbf{Y}_{\cdot j}). \tag{S22}$$

**Trivial reference predictors.** On the exact same donor-partitioned test splits and $\log(1 + \mathrm{CP10k})$ HVG targets, we evaluated three embedding-free baselines: predicting the training-split gene mean for every test cell, predicting the training-split gene median, and predicting all zeros.

#### 5. Direct linear probes of rare cell types and cellular programs

Base direct linear probes (Table S22) are evaluated on $N_{\mathrm{sub}} = 1{,}000$ randomly sampled cells per cohort (HL, MS, C19, and MH; seed 42) using five-fold cross-validation with Ridge regularization ($\alpha = 10$) and within-fold $z$-standardization of cached embeddings.

**Stratum-specific linear classification F1 ($\mathrm{F1}_{\mathrm{rare}}$ and $\mathrm{F1}_{\mathrm{abundant}}$; range $[0, 1]$, $\uparrow$).** Pooled out-of-fold predictions from five-fold cell-stratified Ridge classification yield one-vs-rest per-class $\mathrm{F1}_c$ ($0/0 \equiv 0$), averaged unweighted across rare cell types $\mathcal{C}_{\mathrm{rare}}$ ($< 1\%$ full-cohort frequency) and abundant cell types $\mathcal{C}_{\mathrm{abundant}}$ ($\geq 5\%$ full-cohort frequency):

$$\mathrm{F1}_{\mathrm{rare}} = \frac{1}{|\mathcal{C}_{\mathrm{rare}}|} \sum_{c \in \mathcal{C}_{\mathrm{rare}}} \mathrm{F1}_c, \qquad \mathrm{F1}_{\mathrm{abundant}} = \frac{1}{|\mathcal{C}_{\mathrm{abundant}}|} \sum_{c \in \mathcal{C}_{\mathrm{abundant}}} \mathrm{F1}_c. \tag{S23}$$

To test sensitivity to embedding width, regularization strength, subsampling, and donor leakage, we repeated the rare-class probe across six additional control regimes combining native versus PCA-192-matched embeddings, nested `RidgeCV` regularization, 10 repeated subsamples, and donor-grouped five-fold cross-validation on $N_{\mathrm{sub}} = 3{,}000$ cells per cohort, alongside a direct 192-dimensional PCA on $\log(1 + \mathrm{CP10k})$ HVGs (Table S22).

**Pooled out-of-fold $R^2$ for cellular programs (range $(-\infty, 1]$, ↑).** Out-of-fold Ridge regression predictions $\hat{y}_i$ from five donor-grouped folds (or five cell-stratified folds when fewer than five donors are sampled) are pooled across all $N_{\mathrm{sub}}$ cells for each continuous target $y_i$ (library depth $\log_{10}(\mathrm{UMI} + 1)$, ribosomal and mitochondrial transcript fractions, cell-cycle proliferation, and interferon signaling):

$$R^2 = 1 - \frac{\sum_{i=1}^{N_{\mathrm{sub}}} (y_i - \hat{y}_i)^2}{\sum_{i=1}^{N_{\mathrm{sub}}} (y_i - \bar{y})^2}, \tag{S24}$$

where $\bar{y} = \frac{1}{N_{\mathrm{sub}}} \sum_{i=1}^{N_{\mathrm{sub}}} y_i$, and $R^2 < 0$ indicates cross-donor linear prediction error exceeding the cohort-mean predictor. To check whether interferon $R^2$ was driven by constitutive MHC-I genes or sequencing covariates, we additionally scored a clean canonical-ISG target that excluded *B2M*, *HLA-A*, and mouse *H2-K1* and regressed out library depth and ribosomal transcript fraction before five-fold evaluation (Table S22).

#### 6. Genetic perturbation response and reference-baseline diagnostics

Across $S = |\mathcal{G}_{\mathrm{pert}}|$ held-out $(\text{condition}, \text{cell type})$ test strata $g = (q, c) \in \mathcal{G}_{\mathrm{pert}}$ and all $G$ dataset genes, let $\bar{\mathbf{y}}_{\mathrm{pert},g} \in \mathbb{R}^G$ and $\hat{\bar{\mathbf{y}}}_{\mathrm{pert},g} \in \mathbb{R}^G$ denote observed and predicted mean expression (from the shared residual decoder in main-text Equation 1) over test perturbed cells in stratum $g$, and let $\bar{\mathbf{y}}_{\mathrm{ctrl},c} \in \mathbb{R}^G$ denote mean expression across all observed unperturbed control cells of matching cell type $c$. Observed and predicted mean expression-shift vectors in the dataset's evaluation expression space are

$$\Delta_g = \bar{\mathbf{y}}_{\mathrm{pert},g} - \bar{\mathbf{y}}_{\mathrm{ctrl},c}, \qquad \hat{\Delta}_g = \hat{\bar{\mathbf{y}}}_{\mathrm{pert},g} - \bar{\mathbf{y}}_{\mathrm{ctrl},c}. \tag{S25}$$

The four primary perturbation metrics follow the order in main-text Figure 6, followed by the zero-change, training-mean, and single-gene additive diagnostics in Table S25.

**Across-gene shift correlations (Pearson $\Delta$ and Spearman $\Delta$; range $[-1, 1]$, ↑).** Averaged over strata $\mathcal{V}_\Delta = \{g \in \mathcal{G}_{\mathrm{pert}} : \max_j \hat{\Delta}_{g,j} > \min_j \hat{\Delta}_{g,j},\ \max_j \Delta_{g,j} > \min_j \Delta_{g,j}\}$ across all $G$ genes using $\mathrm{corr}(\cdot, \cdot)$ from Equation (S17):

$$\text{Pearson } \Delta = \frac{1}{|\mathcal{V}_\Delta|} \sum_{g \in \mathcal{V}_\Delta} \mathrm{corr}(\hat{\Delta}_g,\ \Delta_g), \tag{S26}$$

$$\text{Spearman } \Delta = \frac{1}{|\mathcal{V}_\Delta|} \sum_{g \in \mathcal{V}_\Delta} \mathrm{corr}\big(\mathrm{rank}(\hat{\Delta}_g),\ \mathrm{rank}(\Delta_g)\big). \tag{S27}$$

**Genome-wide shift mean absolute error ($\mathrm{MAE}_\Delta$; range $[0, \infty)$, ↓).** Weighting all $S$ strata and $G$ genes equally:

$$\mathrm{MAE}_\Delta = \frac{1}{S\,G} \sum_{g \in \mathcal{G}_{\mathrm{pert}}} \sum_{j=1}^{G} |\hat{\Delta}_{g,j} - \Delta_{g,j}|. \tag{S28}$$

**Differential-expression overlap score (DES; range $[0, 1]$, ↑).** Following the Virtual Cell Challenge overlap-at-$N$ protocol, within each stratum $g = (q, c)$ with $n_{\mathrm{pert},g} \geq 3$ perturbed test cells and $n_{\mathrm{ctrl},c} \geq 3$ matching control cells, two-sided Mann–Whitney $U$ tests are run per gene $j \in \{1, \ldots, G\}$ (exact distribution when $\min(n_{\mathrm{pert},g}, n_{\mathrm{ctrl},c}) \leq 8$ without ties; asymptotic normal approximation otherwise), comparing observed perturbed cells against observed controls and independently comparing predicted perturbed cells against the same observed controls. Within each comparison, gene-wise $p$-values are adjusted by Benjamini–Hochberg false discovery rate

($q < 0.05$). Let $D_g^{\text{true}}$ denote the observed significant gene set ($N_g = |D_g^{\text{true}}|$), and let $D_g^{\text{pred}}$ denote predicted significant genes ($q_j^{\text{pred}} < 0.05$); ranking $D_g^{\text{pred}}$ by descending $|\hat{\Delta}_{g,j}|$ (ties broken stably) and retaining the top $\min(N_g, |D_g^{\text{pred}}|)$ genes gives $\tilde{D}_g^{\text{pred}}$. Averaging over valid strata $\mathcal{G}_{\text{valid}} = \{g : N_g \geq 1\}$ yields

$$\text{DES} = \frac{1}{|\mathcal{G}_{\text{valid}}|} \sum_{g \in \mathcal{G}_{\text{valid}}} \frac{|D_g^{\text{true}} \cap \tilde{D}_g^{\text{pred}}|}{N_g}. \tag{S29}$$

**Zero-change skill scores and DE-gene subsets ($\text{MSE}_\Delta$, $\text{MAE}_\Delta$ skill, and $\text{MSE}_\Delta$ skill, Table S25; skill range $(-\infty, 1]$, ↑).** Genome-wide mean squared error and the reference errors of the zero-change predictor ($\hat{\Delta}_g = \mathbf{0}$) are

$$\text{MSE}_\Delta = \frac{1}{S\,G} \sum_{g \in \mathcal{G}_{\text{pert}}} \sum_{j=1}^{G} (\hat{\Delta}_{g,j} - \Delta_{g,j})^2, \tag{S30}$$

$$\text{MAE}_{\text{control}} = \frac{1}{S\,G} \sum_{g \in \mathcal{G}_{\text{pert}}} \sum_{j=1}^{G} |\Delta_{g,j}|, \qquad \text{MSE}_{\text{control}} = \frac{1}{S\,G} \sum_{g \in \mathcal{G}_{\text{pert}}} \sum_{j=1}^{G} \Delta_{g,j}^2. \tag{S31}$$

Normalizing each model's error against these zero-change baselines gives the skill scores

$$\text{MAE}_\Delta \text{ skill} = 1 - \frac{\text{MAE}_\Delta}{\text{MAE}_{\text{control}}}, \qquad \text{MSE}_\Delta \text{ skill} = 1 - \frac{\text{MSE}_\Delta}{\text{MSE}_{\text{control}}}, \tag{S32}$$

where $0$ matches predicting unperturbed control expression and positive values indicate lower error than predicting no perturbation shift. To separate signal on responsive genes from genome-wide background noise, the same skill formulas were also evaluated on the top 20 and top 100 differentially expressed genes per stratum (DE-20 and DE-100, selected by observed perturbation response).

**Training-mean and single-gene additive reference baselines.** On all four Perturb-seq screens, we evaluated a training-mean baseline predicting the average observed shift across training perturbations ($\hat{\Delta}_g^{\text{mean}} = \frac{1}{|\mathcal{G}_{\text{train}}|} \sum_{g' \in \mathcal{G}_{\text{train}}} \Delta_{g'}$). On Norman held-out two-gene combinations $(a + b)$, we additionally evaluated a single-gene additive predictor summing the observed mean shifts of the two constituent single-gene perturbations ($\hat{\Delta}_{(a+b),c}^{\text{add}} = \Delta_{a,c}^{\text{train}} + \Delta_{b,c}^{\text{train}}$; Table S25).

#### 7. Composite ranking and rank-correlation diagnostics

**Average-tie block ranking (range $[1, 16]$, ↓).** Within each (dataset $d$, seed $s$, metric $k$) evaluation block across all $16$ methods, lower-is-better metrics ($\text{MAE}_{\text{expr}}$ and $\text{MAE}_\Delta$) are negated prior to descending sort. If exact floating-point ties occupy sorted positions $a$ through $b$ ($1 \leq a \leq b \leq 16$), each tied method $m$ receives the midpoint rank

$$r_{m,d,s,k} = \frac{a + b}{2} \in [1, 16], \tag{S33}$$

where lower ranks indicate stronger performance. Dataset-level task ranks average $r_{m,d,s,k}$ equally across the three random seeds ($s \in \{41, 42, 43\}$) and four primary metrics ($k \in \{1, \ldots, 4\}$); panel and 12-cohort summary ranks average dataset-level ranks equally across their constituent cohorts.

**Cross-model Spearman rank correlation (range $[-1, 1]$).** Associations between two summary vectors $\mathbf{u}, \mathbf{v}$ across methods are measured descriptively (without parametric hypothesis tests) using $\text{corr}(\cdot, \cdot)$ from Equation (S17):

$$\rho_{\text{rank}}(\mathbf{u}, \mathbf{v}) = \text{corr}\big(\text{rank}(\mathbf{u}), \text{rank}(\mathbf{v})\big). \tag{S34}$$

This statistic is applied to cross-task biological channel concordance across the $10$ scFMs with clustering $\mathrm{uARI} > 0.25$ (Table S23), within-block pairwise metric-rank agreement averaged over all $12$ cohorts and three seeds (Figure S21), model-attribute versus task-rank correlations (Table S26), and 52 leave-one-dataset-out (LODO) / 52 leave-one-metric-out (LOMO) rank stability checks across all static and perturbation panels (Table S28).

### NOTES ON SUPPLEMENTAL FIGURES

The 3-by-4 grids in Figures S1–S16 unpack the four static tasks in main-text Figures 2–5 by metric and dataset. Rows are human organs, human diseases, and mouse organs; columns are the four datasets within each panel. Each panel is drawn with its own $y$-axis scaled to the observed method range so contrasts remain visible; in Figures S13–S16, reconstruction metrics are scored on the $\log(1+\mathrm{CP10k})$ scale (with coverage-penalized Pearson $r$ and Spearman $\rho$, and non-zero top-50 expressed-HVG overlap in Figure S16). Each box summarizes random seeds 41, 42, and 43 using interpolated quartiles, a median line, and observed minimum–maximum whiskers, with individual seed markers and outlier markers hidden. These represent repeated runs or donor splits rather than independent biological replicates or confidence intervals, and no hypothesis tests are displayed.

#### Perturbation prediction

**Perturbation metrics (Figure S17).** Shows the four primary perturbation metrics across Norman, Adamson, Replogle K562, and Replogle RPE1: (a) Pearson $\Delta$, (b) Spearman $\Delta$, (c) MAE $\Delta$ (lower-is-better), and (d) differential-expression overlap score (DES, averaging only groups with a defined reference significant-gene set). Each box uses seeds 41–43 with interpolated quartiles, a median line, and min–max whiskers on screen-specific $y$-axes, with seed markers hidden (repeated runs rather than biological replicates). Screen-level zero-change, training-mean, and single-gene additive diagnostics (genome-wide, DE-20, and DE-100) are reported in Table S25.

#### Model scale and rank stability

**Parameter and corpus scale (Figures S18 and S19).** Figure S18 plots audited instantiated-model parameters versus mean task rank, averaged equally over twelve datasets, three seeds, and four primary metrics across all thirteen pretrained scFMs. Parameters include frozen weights and do not measure active compute; no scaling-law fit, confidence interval, or hypothesis test is shown, and dataset-fitted baselines remain in the ranks while excluded from the attribute axis. Figure S19 plots reported corpus size versus mean task rank on the same averaging across ten models with recorded estimates (CellPLM, Tabula, and scPRINT2 lack comparable verified model-specific values; LangCell uses cell–text pairs, whereas other values primarily count cells, and some estimates are family-level or lower bounds). Spearman values are reported in Table S26.

**Seed-to-seed rank stability (Figure S20).** Shows the maximum minus minimum seed-specific mean metric rank for each method and dataset. Each seed rank averages four within-dataset primary-metric ranks over seeds 41–43. Higher values indicate more variation across splits; this is not independent-study replication or a confidence interval.

**Pairwise metric-rank agreement (Figure S21).** Shows the Spearman correlation of method rankings between every pair of primary metrics, calculated within each dataset–seed block and then averaged equally across twelve datasets and three seeds. MAE is oriented lower-is-better before ranking. This is descriptive agreement, with no hypothesis test.

### NOTES ON SUPPLEMENTAL TABLES

| Display items | Analytical block | Summary of contents | Page |
|---|---|---|---|
| Tables S1–S4 | Experimental design | Donor-level split counts (Table S1), shared probes (Table S2), model specifications (Table S3), and evaluation cohorts (Table S4). | 32–35 |
| Tables S5–S7 | Panel composite ranks | Composite mean ranks across human organs (Table S5), human diseases (Table S6), and mouse organs (Table S7). | 35–36 |
| Tables S8–S11 | Human-organ metrics | Three-seed mean primary metrics for clustering, annotation, native batch assessment, and $\log(1 + \mathrm{CP10k})$ reconstruction. | 37–43 |
| Tables S12–S15 | Human-disease metrics | Three-seed mean primary metrics for clustering, annotation, native batch assessment, and $\log(1 + \mathrm{CP10k})$ reconstruction. | 45–51 |
| Tables S16–S19 | Mouse-organ metrics | Three-seed mean primary metrics for clustering, annotation, native batch assessment, and $\log(1 + \mathrm{CP10k})$ reconstruction. | 53–59 |
| Tables S20–S23 | Channels and probes | Biological channel profiles (Table S20), scFoundation-minus-scVI deltas (Table S21), linear probes and 6-regime/clean-ISG audits (Table S22), and channel concordance (Table S23). | 61–62 |
| Tables S24–S26 | Shifts and scale | Mouse-minus-human shifts (Table S24), Perturb-seq reference-baseline skill (Table S25), and scale correlations (Table S26). | 63–64 |
| Tables S27–S29 | Provenance and stability | 208-pair provenance audit (Table S27), 52 LODO/LOMO stability checks (Table S28), and leave-study-out ranks (Table S29). | 65–66 |

#### Model architecture and parameter audit

Table S3 lists architecture, input codes, audited parameter counts, input extent, reported pre-training corpus, and species. Input codes may overlap: A, gene-aligned; V, gene identity and values; R, ranked genes; C, context; S, sampled genes; P, protein-derived features; F, fixed gene panel; B, expression bins. Input extent is sequence length or gene-panel size. Superscript c marks family-level corpus or species estimates that were not verified for the evaluated model. Parameter counts include all unique registered tensors, trainable or frozen, and exclude buffers: GeneCompass's 155.4M regulatory-prior buffer elements are excluded, whereas UCE's 744.8M ESM-2 protein lookup parameters and scPRINT2's 85.8M frozen gene-encoder parameters are included (scPRINT2 also has 16.1M trainable parameters). For mouse cohorts, fixed one-to-one mouse-to-human orthology tables used for human-vocabulary models retained $80.4\%$ of genes and $75.2\%$ to $87.5\%$ of total expression across the four mouse tissues; note also that GeneCompass's released gene-median dictionary lacks mouse gene keys, leaving mouse inputs unscaled. Corpus sources: CellFM (Nat. Commun.), CellPLM (ICLR), GeneCompass (Cell Res.), Geneformer (Nature), LangCell (PMLR), Nicheformer (Nat. Methods), Tabula (NeurIPS), TranscriptFormer (GitHub), UCE (DOI), scFoundation (Nat. Methods), scGPT (GitHub), scLong (Nat. Commun.), and scPRINT2 (GitHub).

#### Leave-study-out overlap sensitivity and provenance audit

Table S29 reports full-panel and leave-study-out annotation ranks for all sixteen methods: *Organs full* uses four human-organ datasets, *Heart excluded* drops the adult-heart cohort, *Diseases full* uses four human-disease datasets, and *MS excluded* drops the MS cohort, with remaining datasets equally weighted. The 208-pair provenance audit (Table S27) classified every model–dataset pair into four mutually exclusive categories: 154 $(74.0\%)$ unresolved, 28 $(13.5\%)$ excluded by documented species scope (the seven strictly human-pretrained models evaluated on the four mouse cohorts), 24 $(11.5\%)$ involving shared public repositories (CZ CELLxGENE Census and scBaseCount) without cell-level logs, and 2 $(1.0\%)$ confirmed sample or donor identifier matches (AH and MS for scFoundation). Because confirmed matches reflect sample or donor identifiers rather than verified cell-barcode overlap, these exclusions do not establish that the remaining human cohorts were absent from all pretraining corpora.

### SUPPLEMENTAL FIGURES

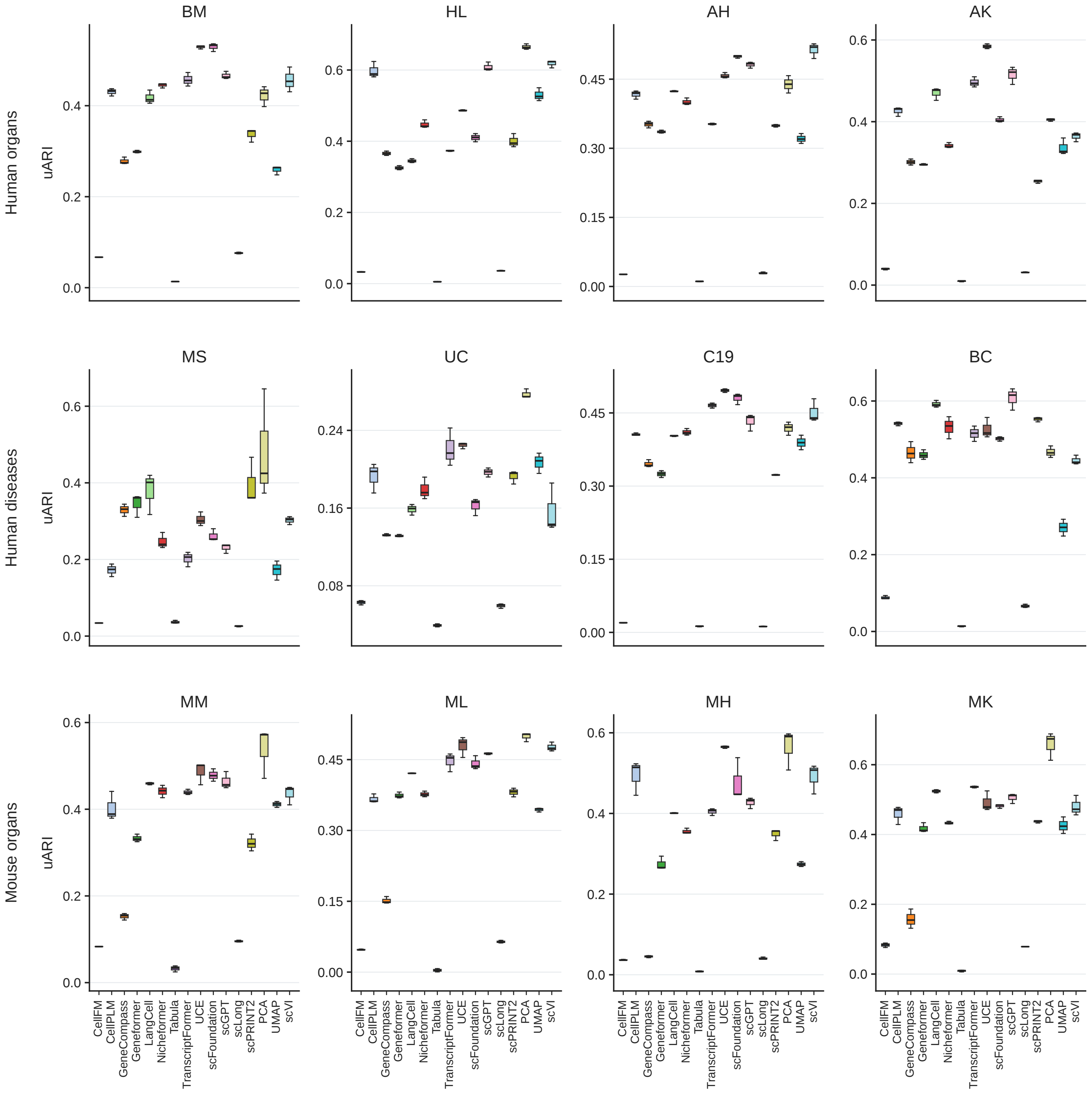


Figure S1: **Unsupervised clustering, uARI.**

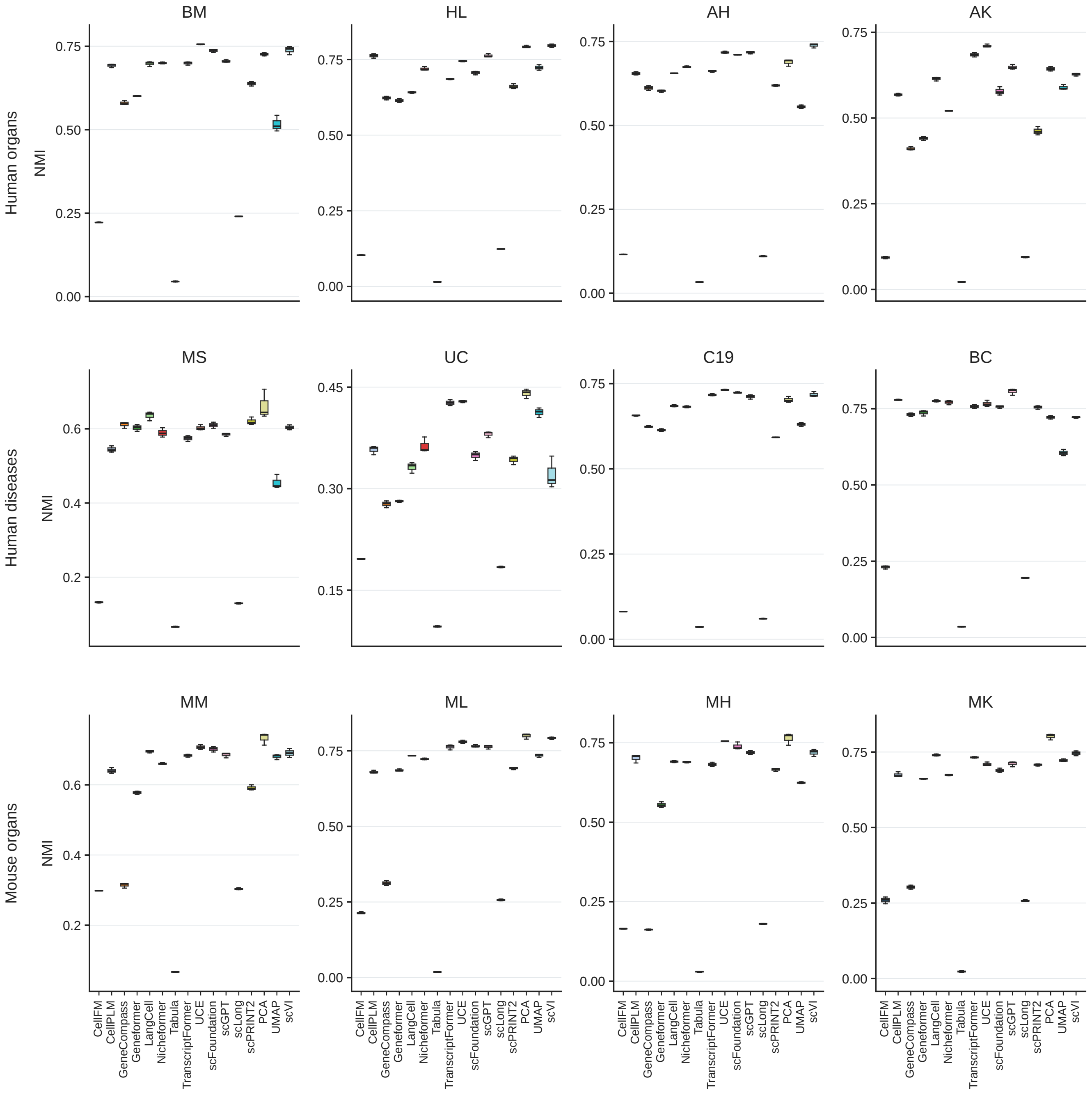


Figure S2: **Unsupervised clustering, NMI.**

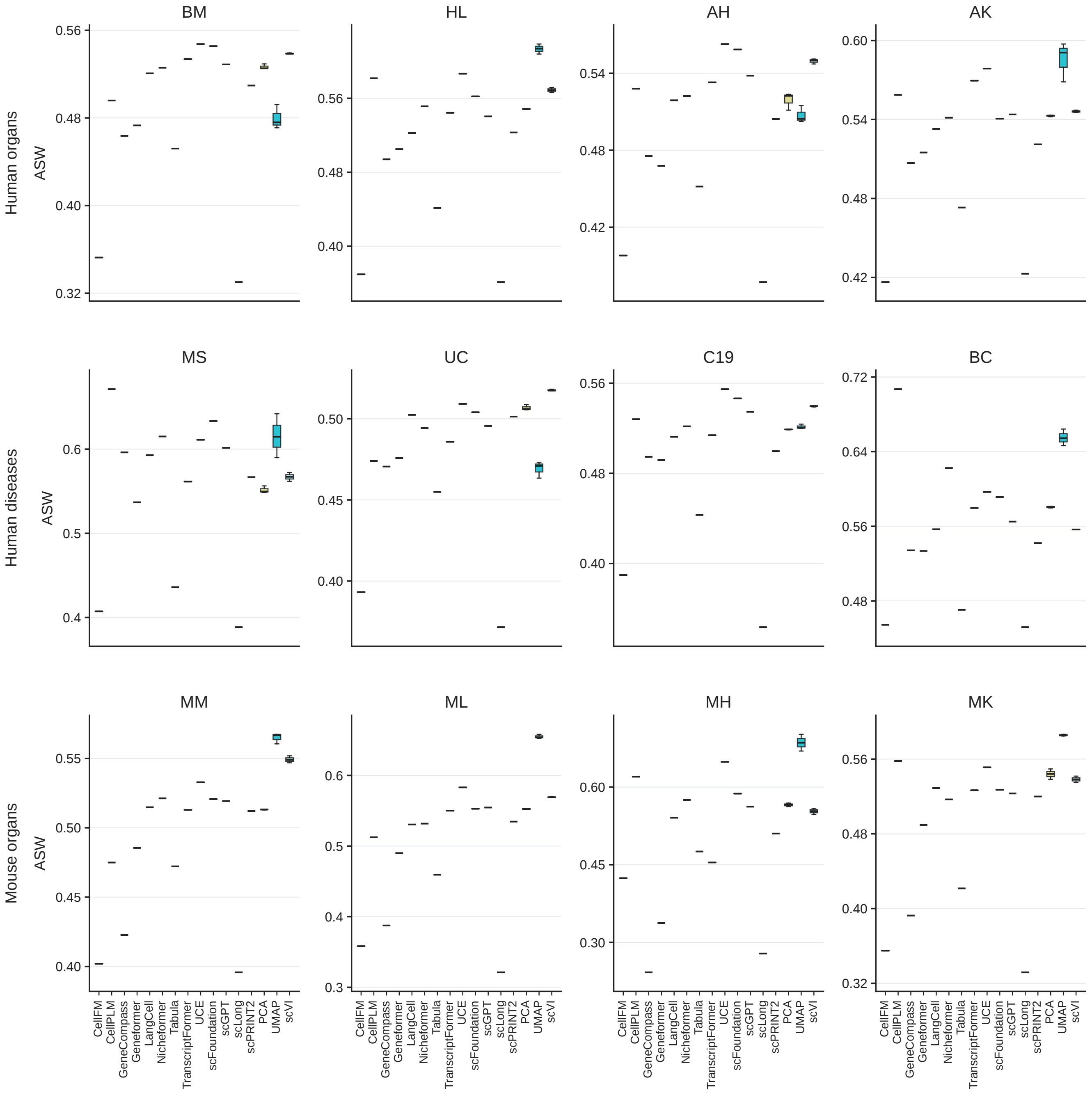


Figure S3: **Unsupervised clustering, ASW.**

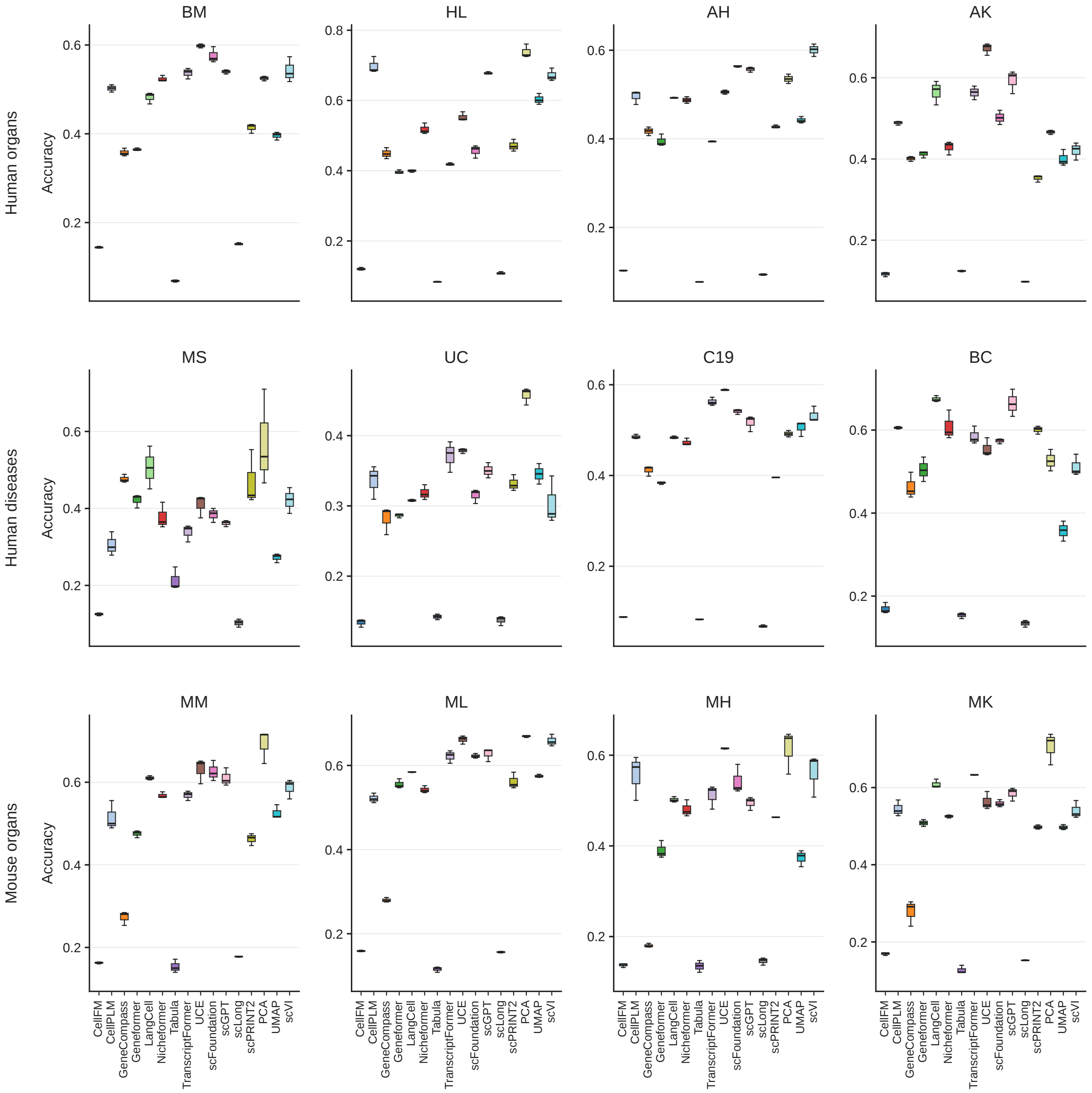


Figure S4: **Unsupervised clustering, accuracy.**

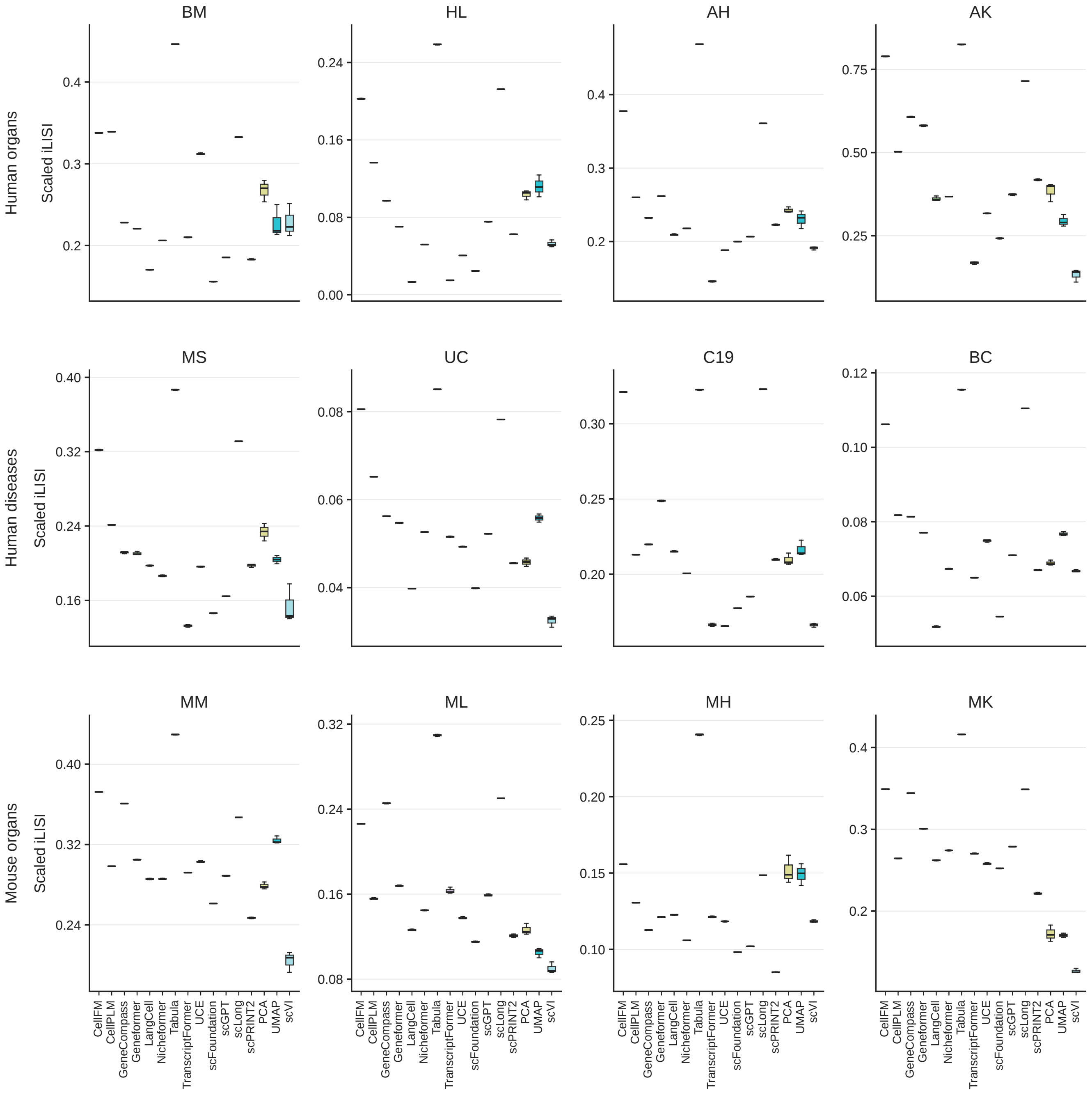


Figure S5: **Native batch assessment, scaled iLISI.**

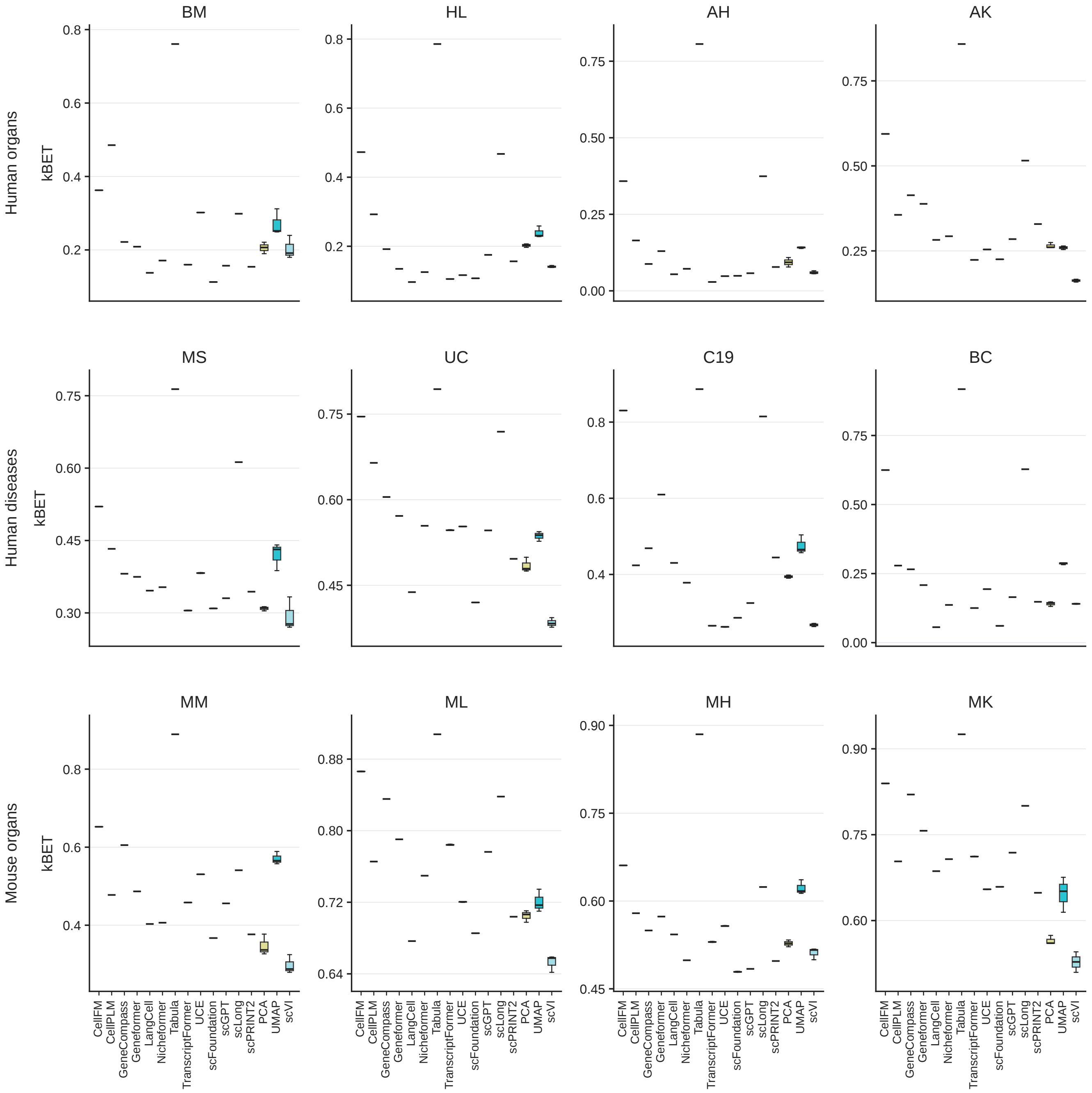


Figure S6: **Native batch assessment, kBET.**

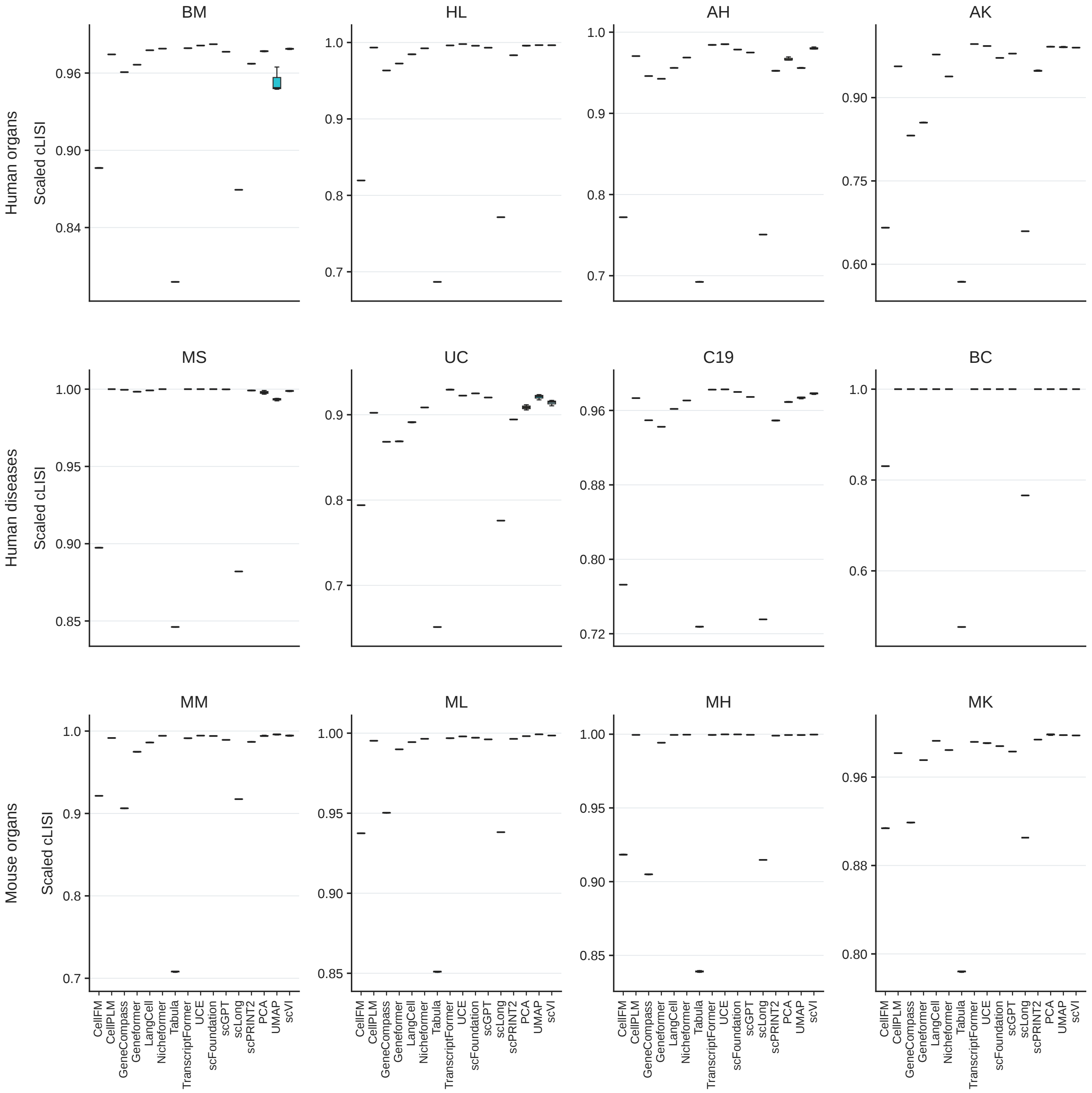


Figure S7: **Native batch assessment, scaled cLISI.**

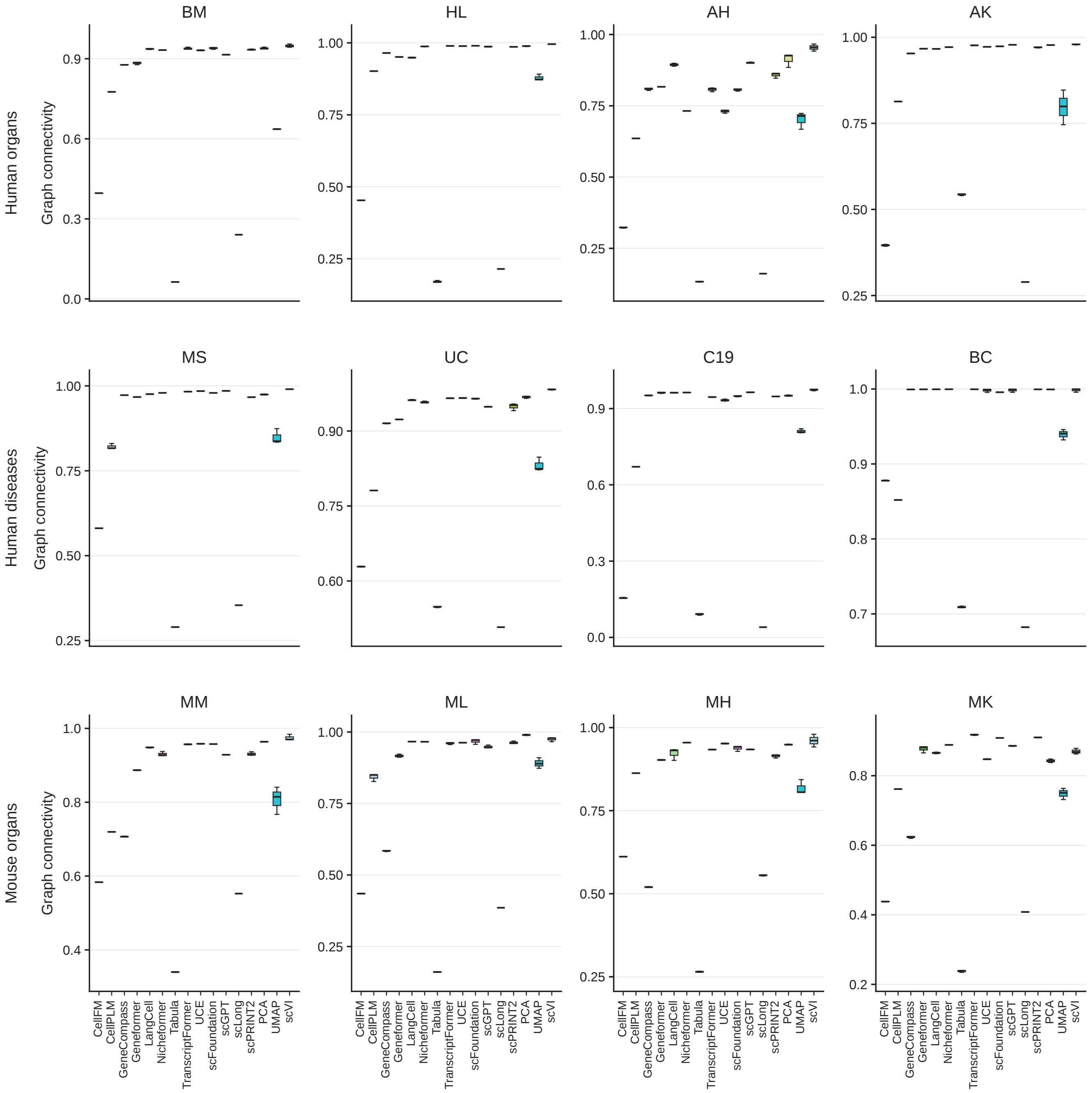


Figure S8: **Native batch assessment, graph connectivity.**

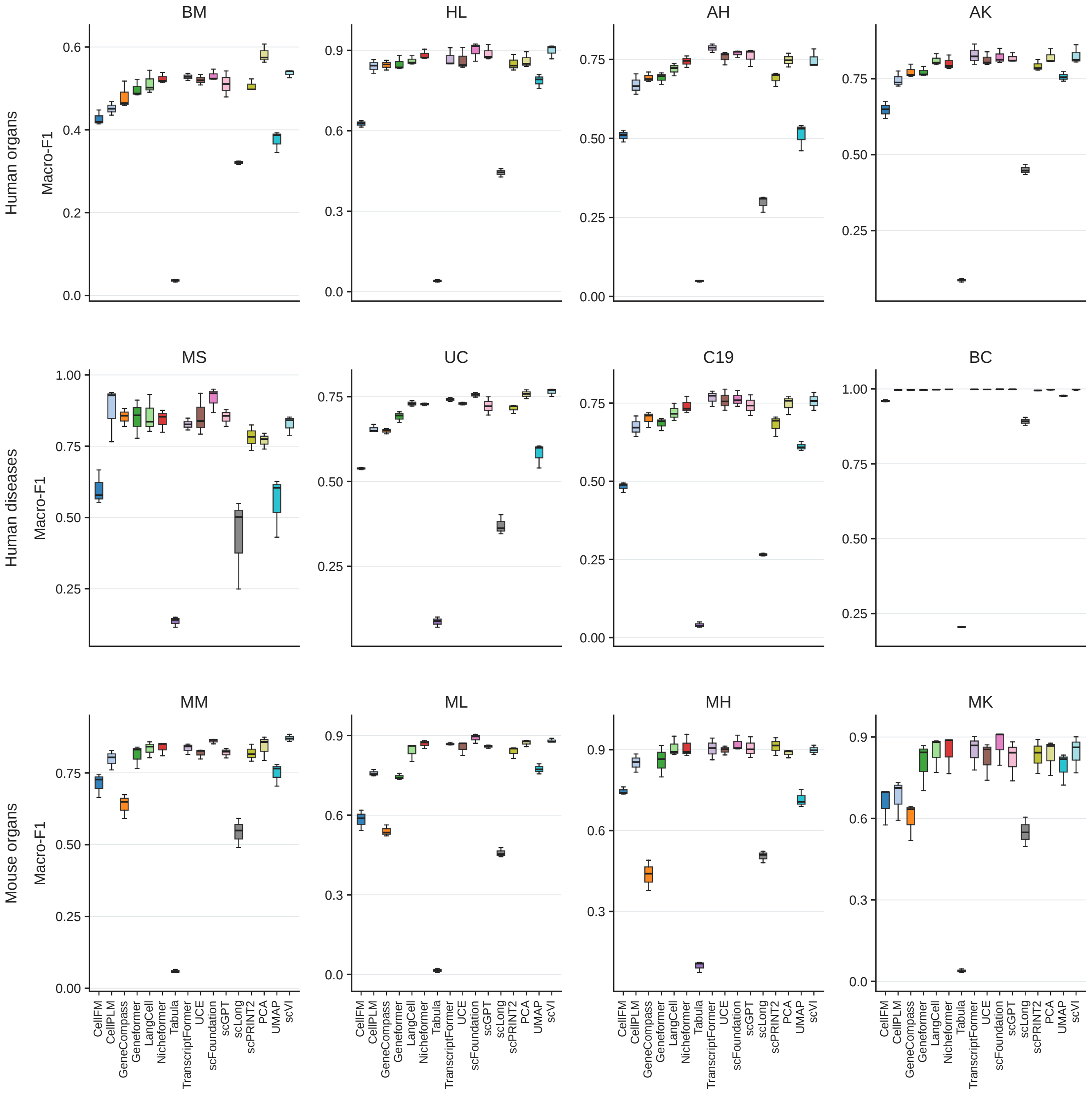


Figure S9: **Cell-type annotation, macro-F1.**

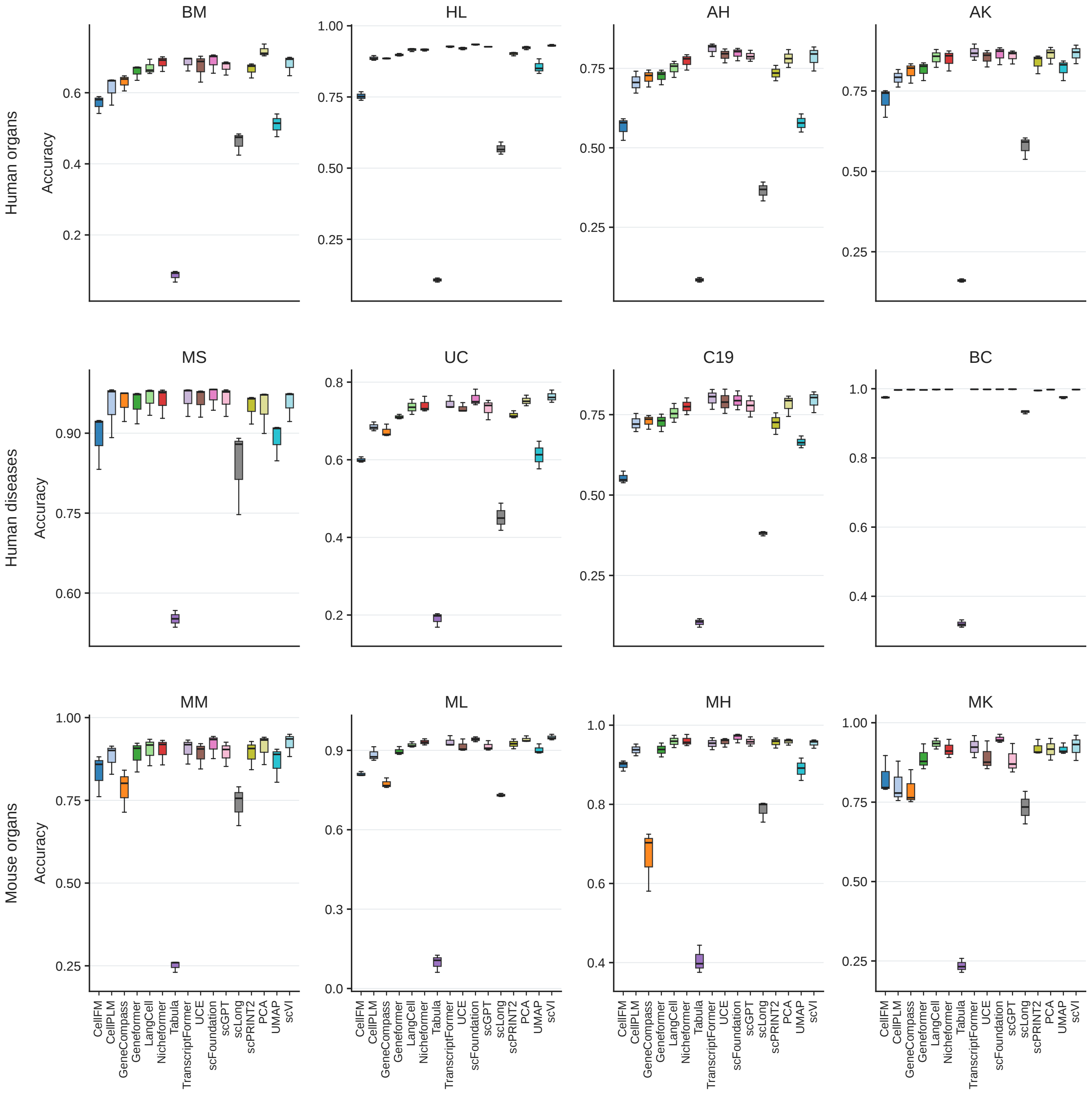


Figure S10: **Cell-type annotation, accuracy.**

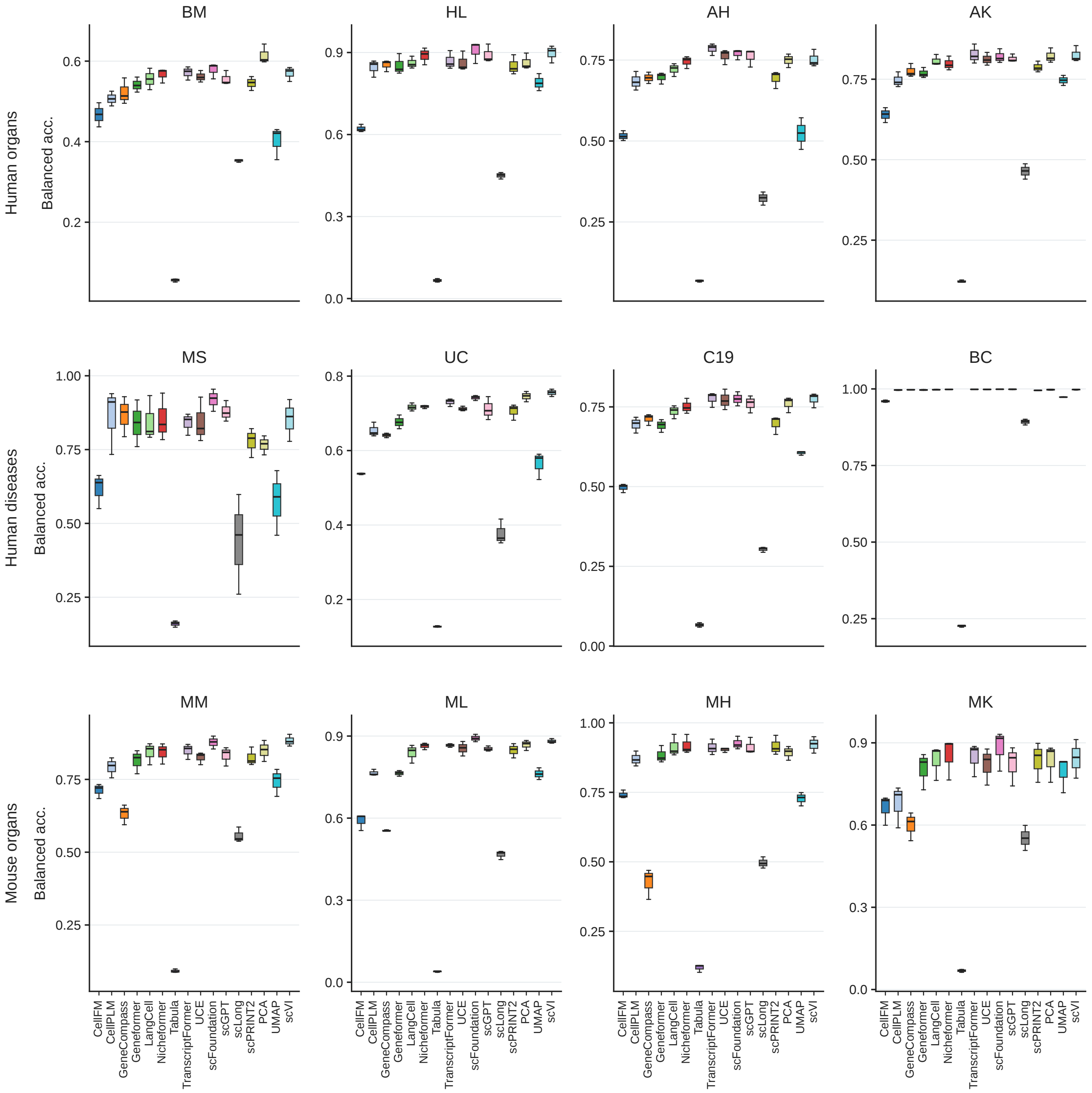


Figure S11: **Cell-type annotation, balanced accuracy.**

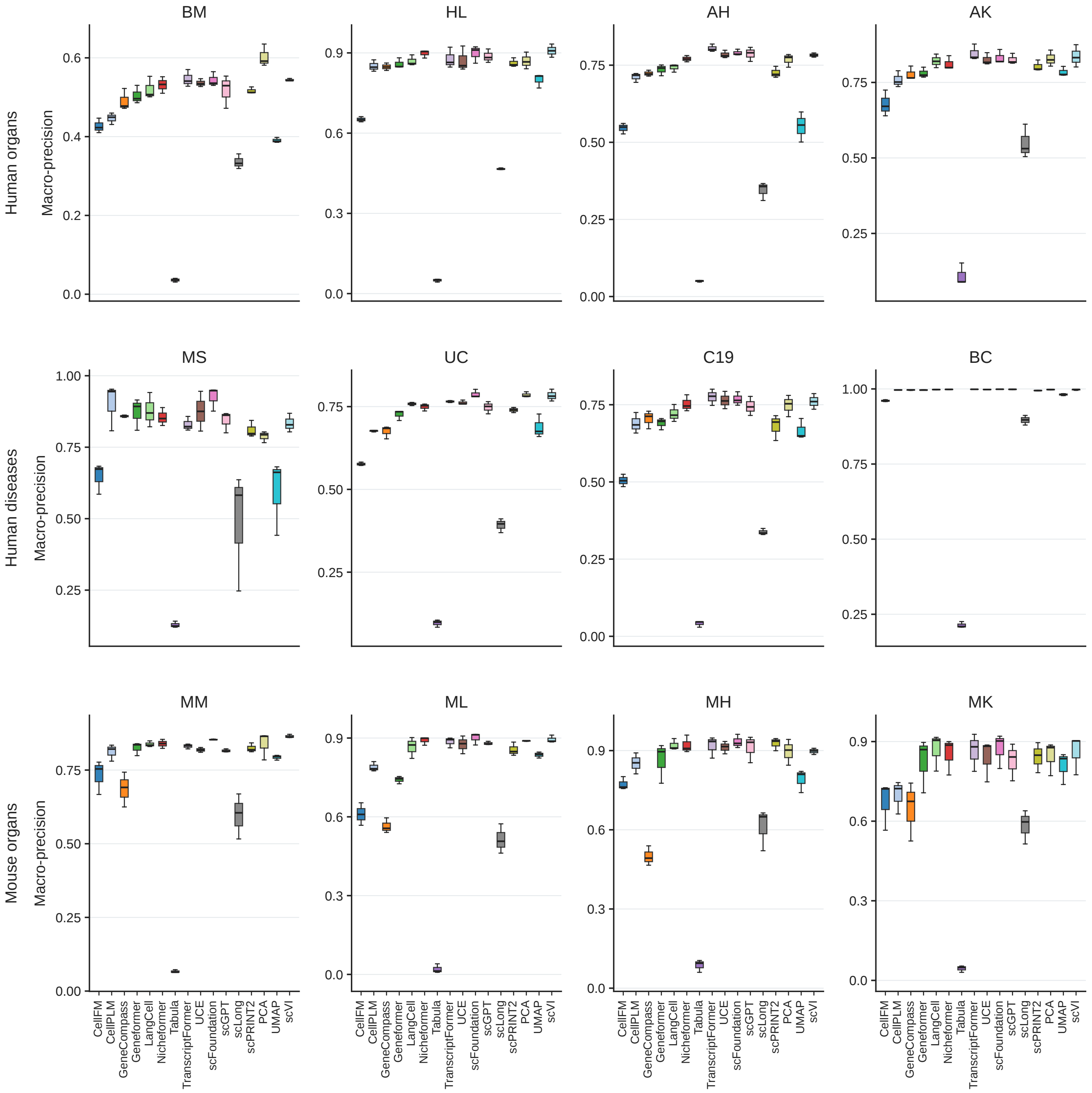


Figure S12: **Cell-type annotation, macro-precision.**

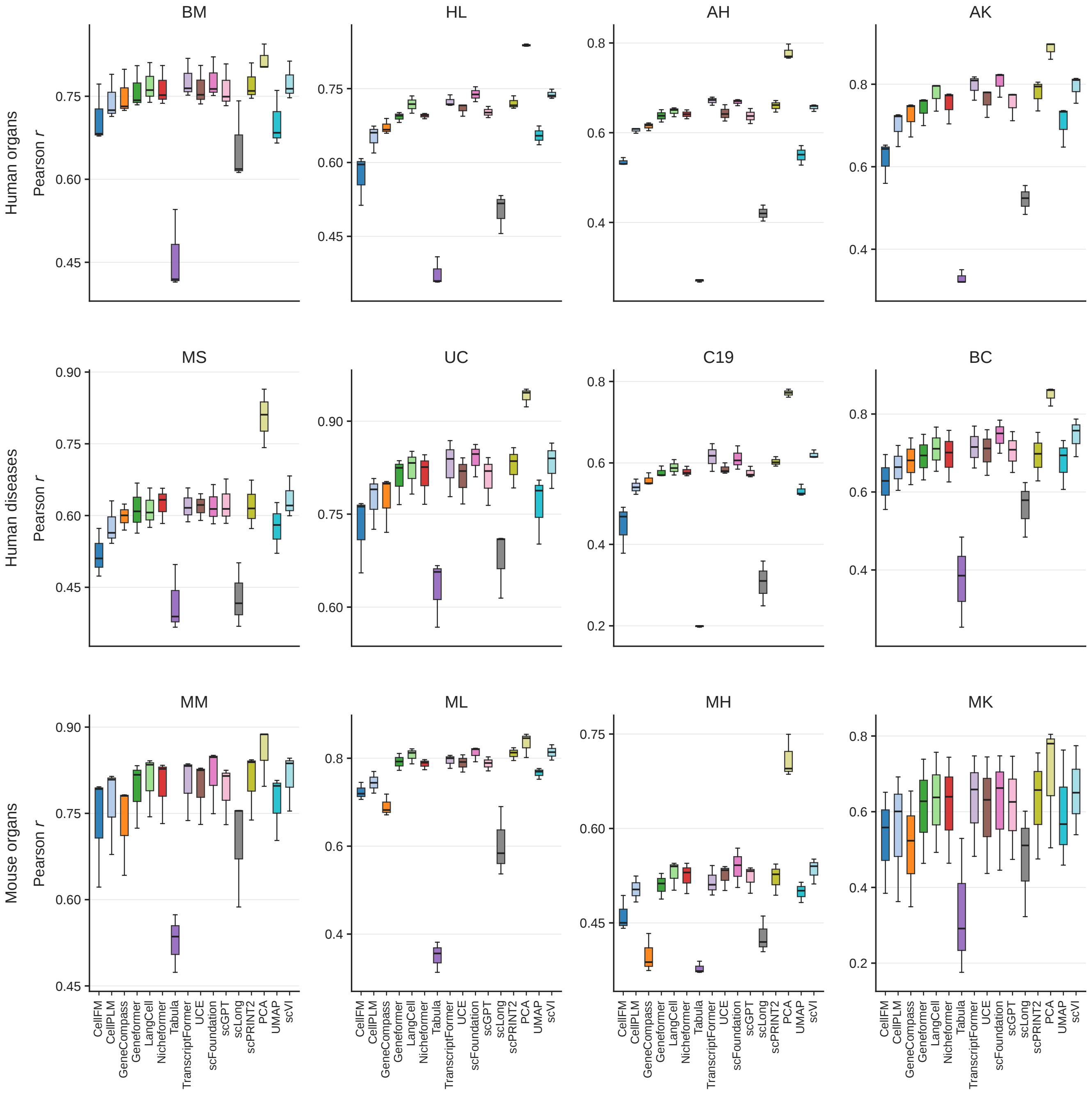


Figure S13: **HVG reconstruction on** $\log(1 + \mathrm{CP10k})$**, coverage-penalized Pearson** $r$**.**

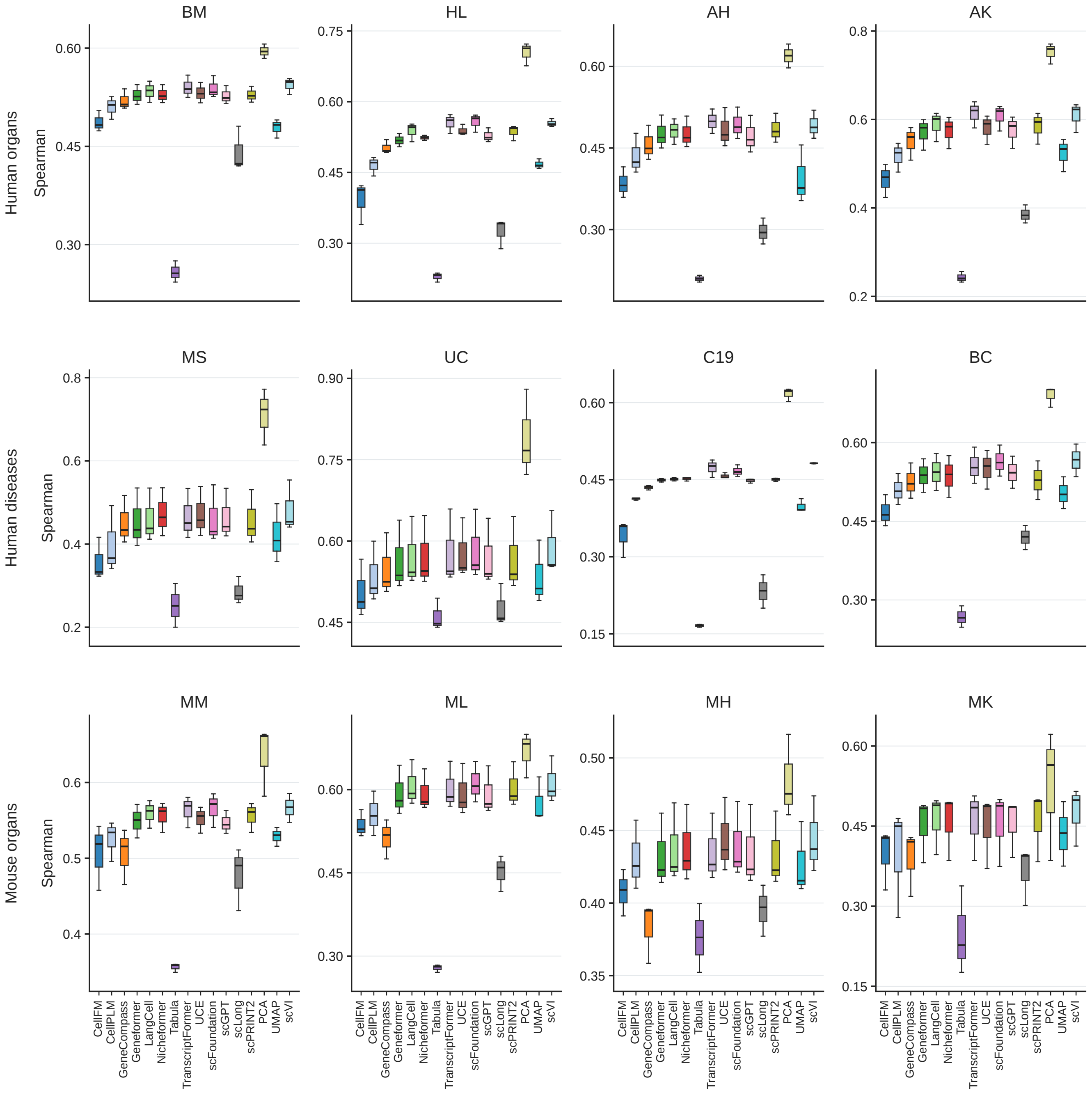


Figure S14: **HVG reconstruction on** $\log(1 + \mathrm{CP10k})$**, coverage-penalized Spearman** $\rho$**.**

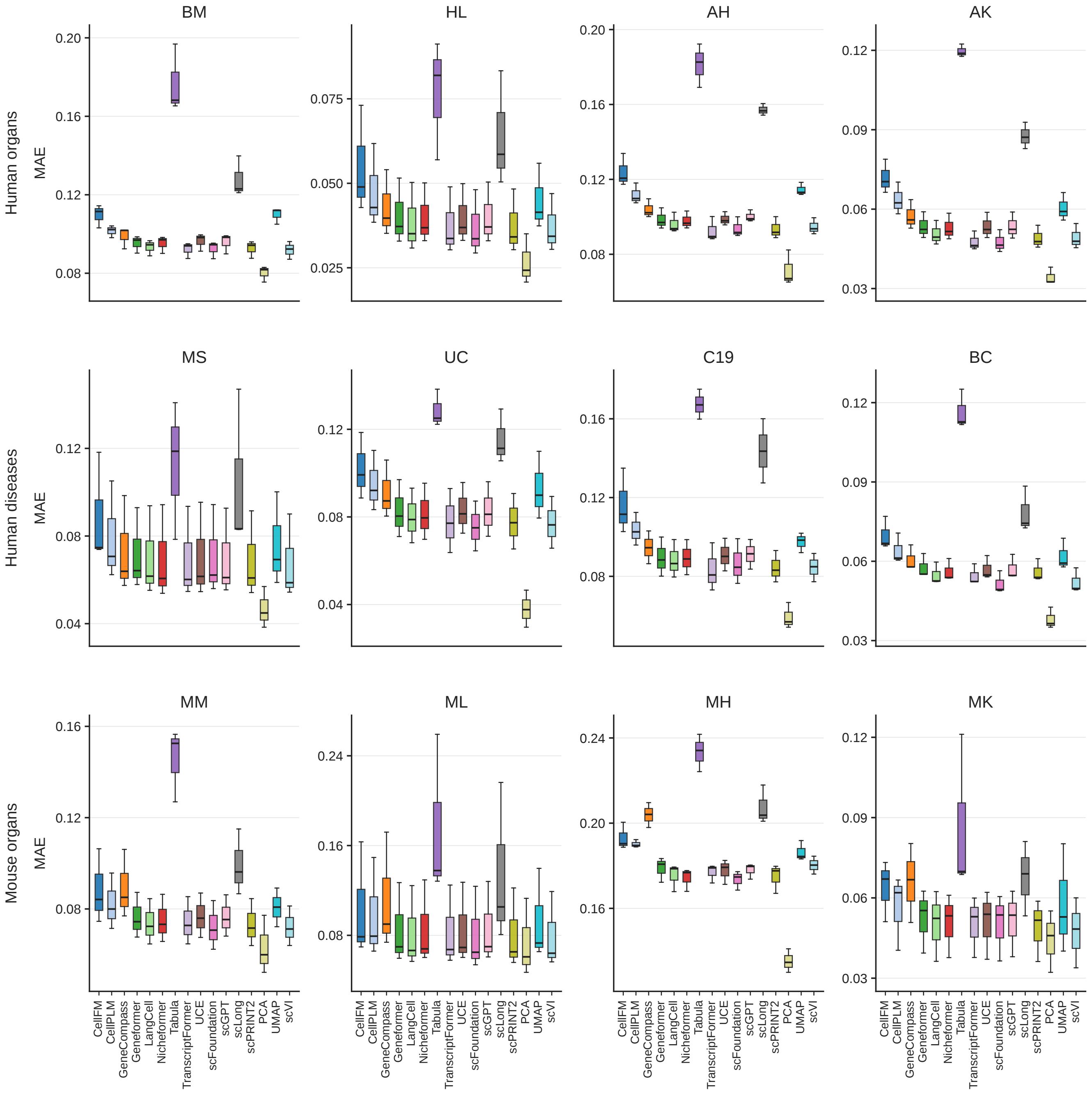


Figure S15: **HVG reconstruction on** $\log(1 + \mathrm{CP10k})$**, MAE.**

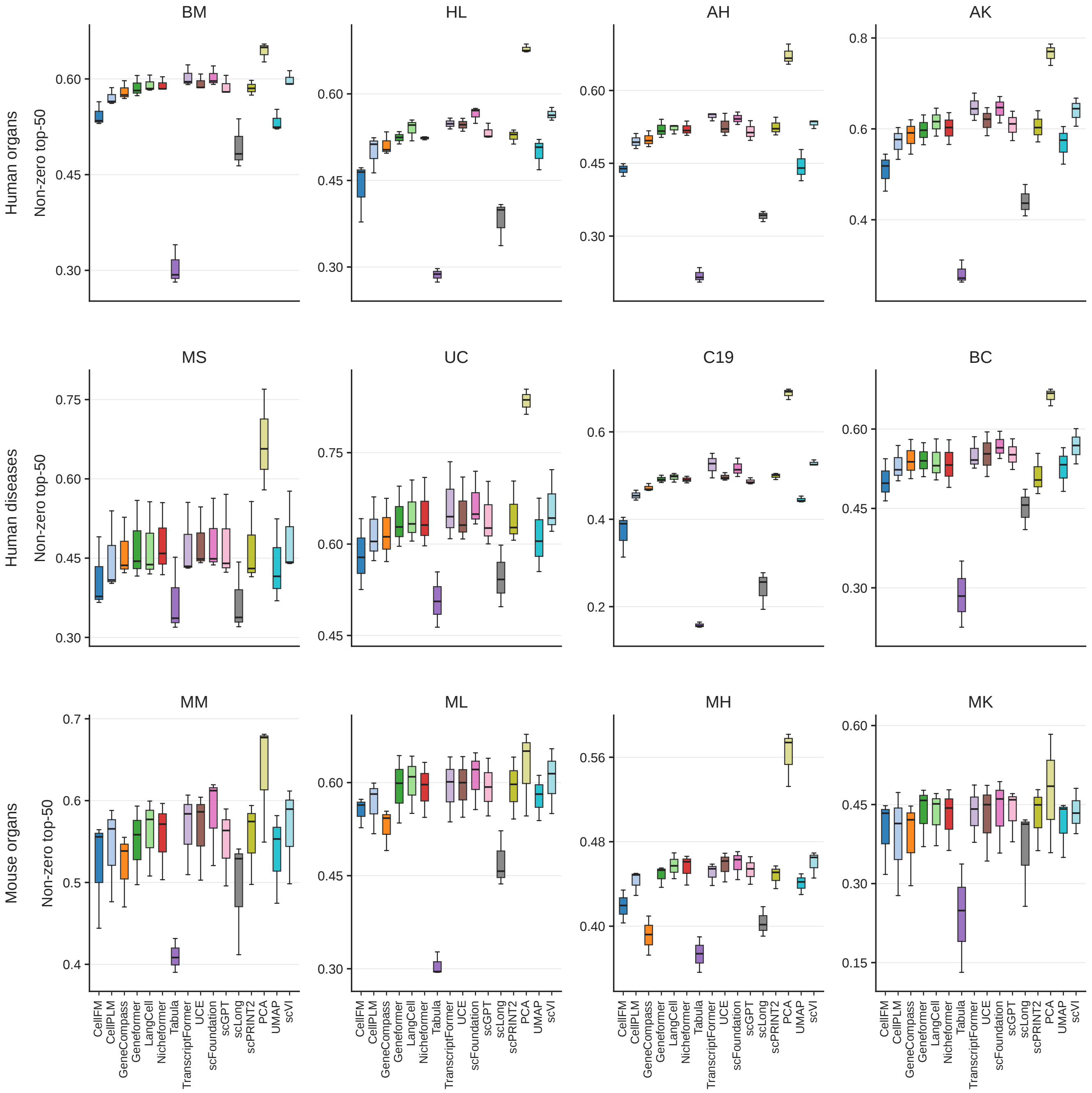


Figure S16: **HVG reconstruction on** $\log(1 + \mathrm{CP10k})$**, non-zero top-50 overlap.**

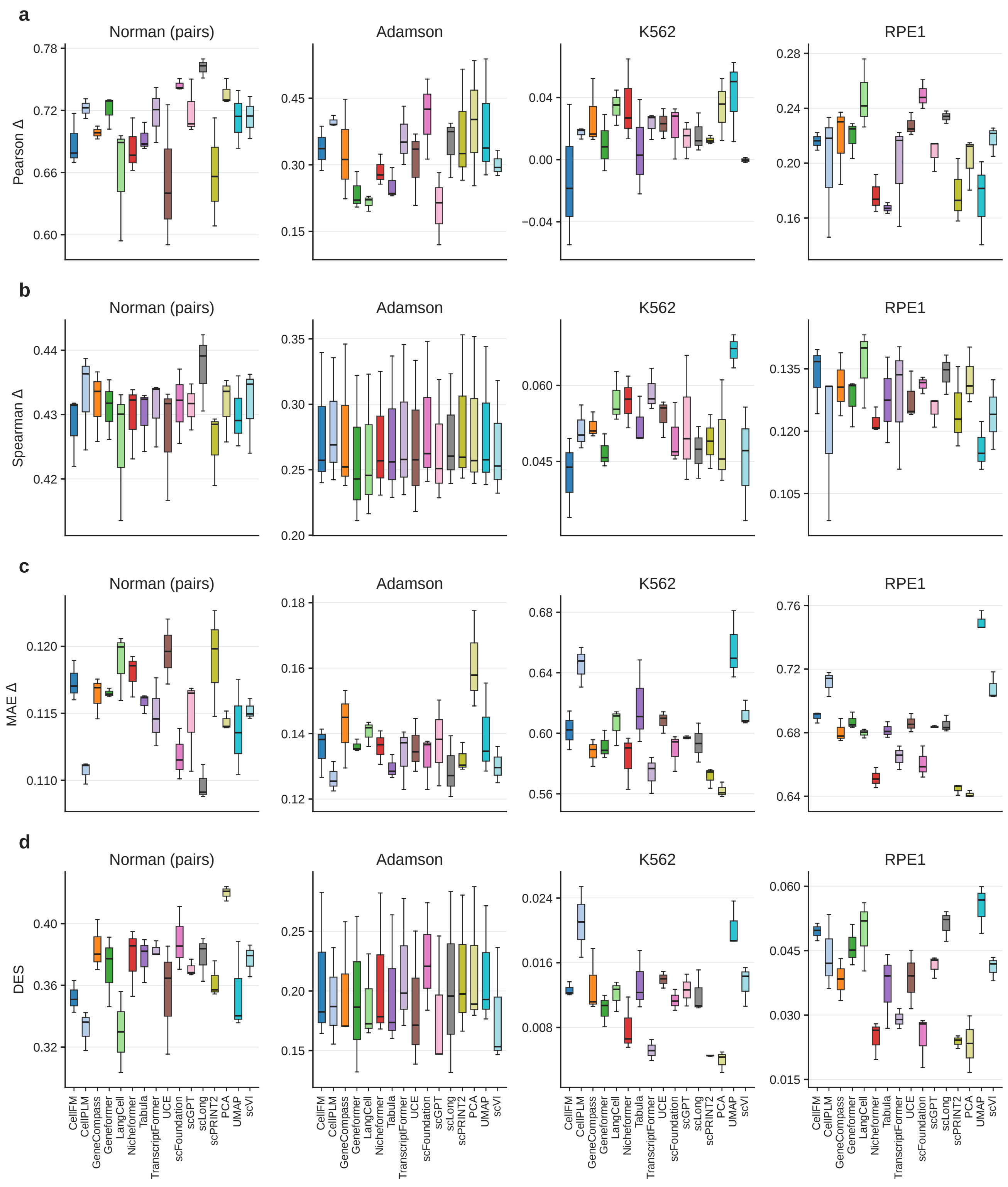


Figure S17: **Perturbation prediction.** (a) Pearson Δ. (b) Spearman Δ. (c) MAE Δ. (d) DES.

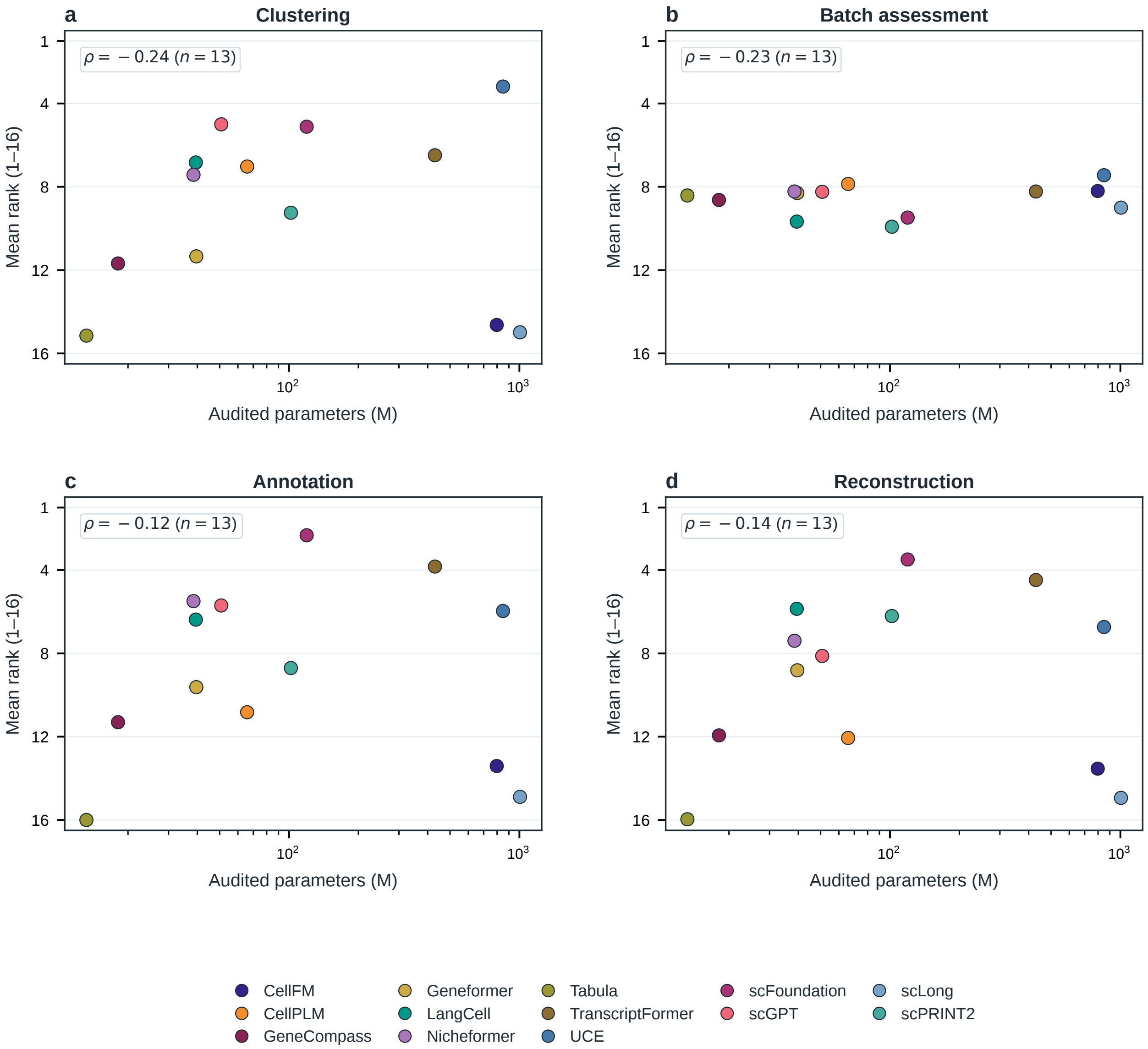


Figure S18: **Parameter count versus task rank.**

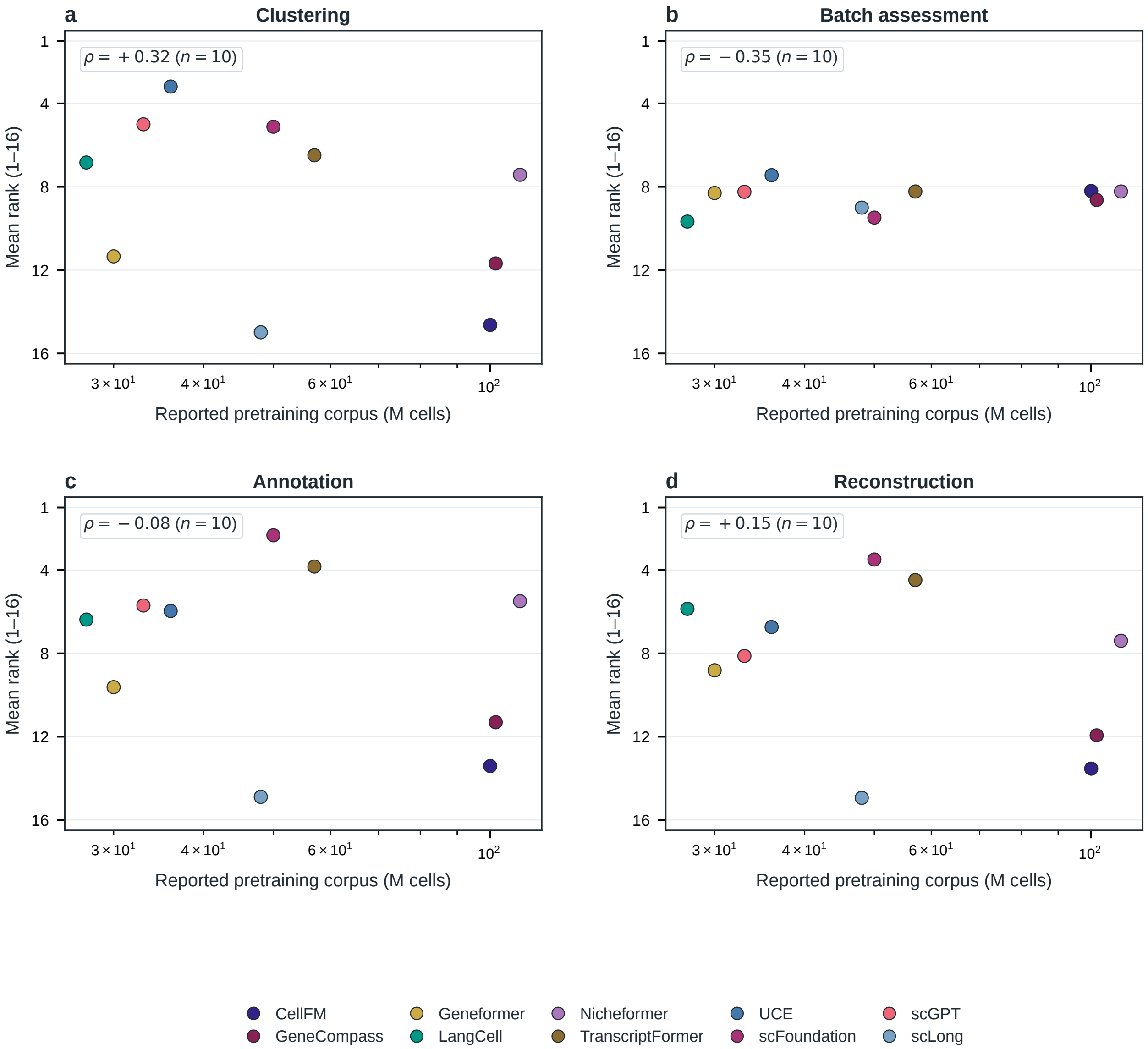


Figure S19: **Corpus size versus task rank.**

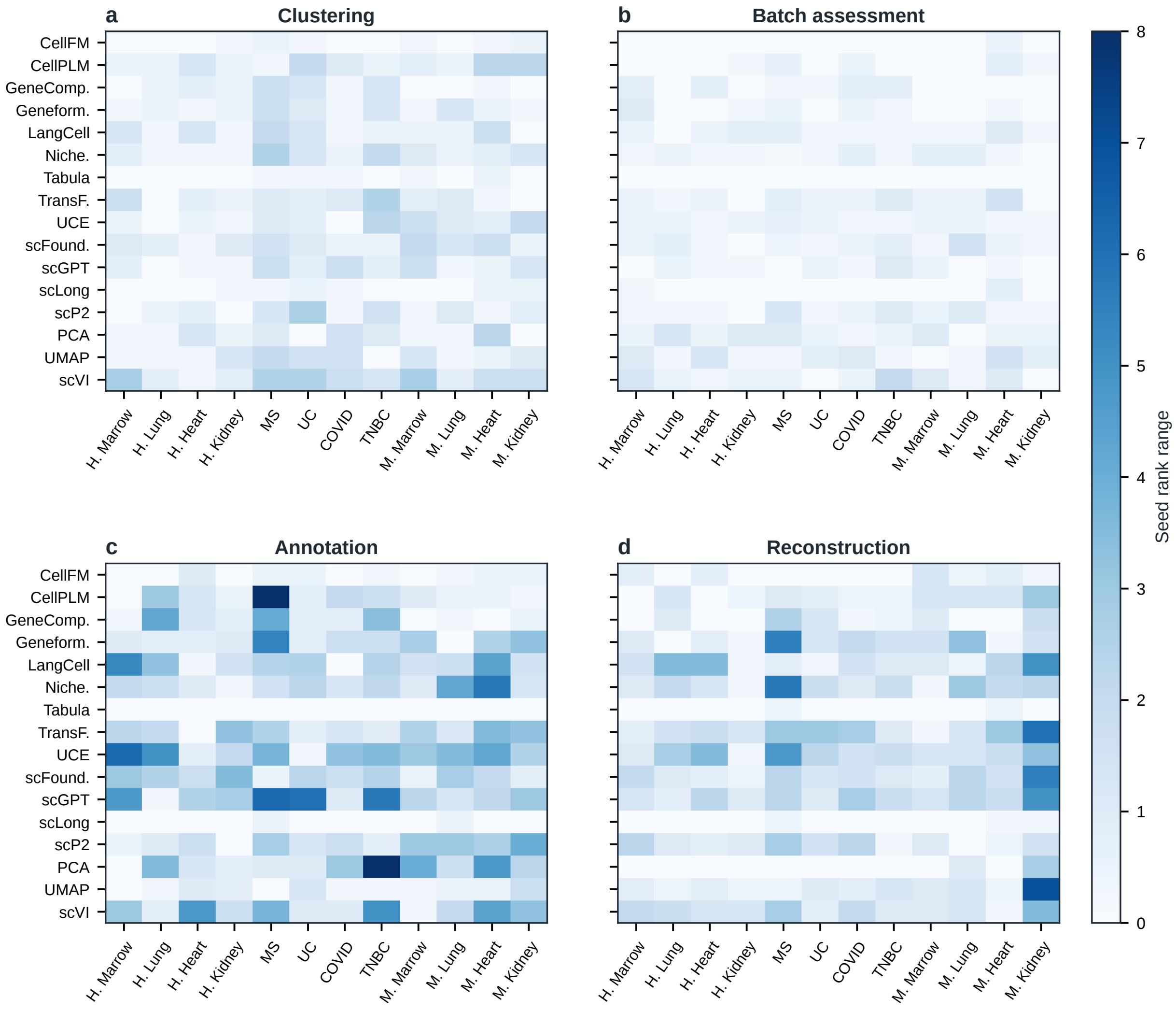


Figure S20: **Seed-to-seed rank range.**

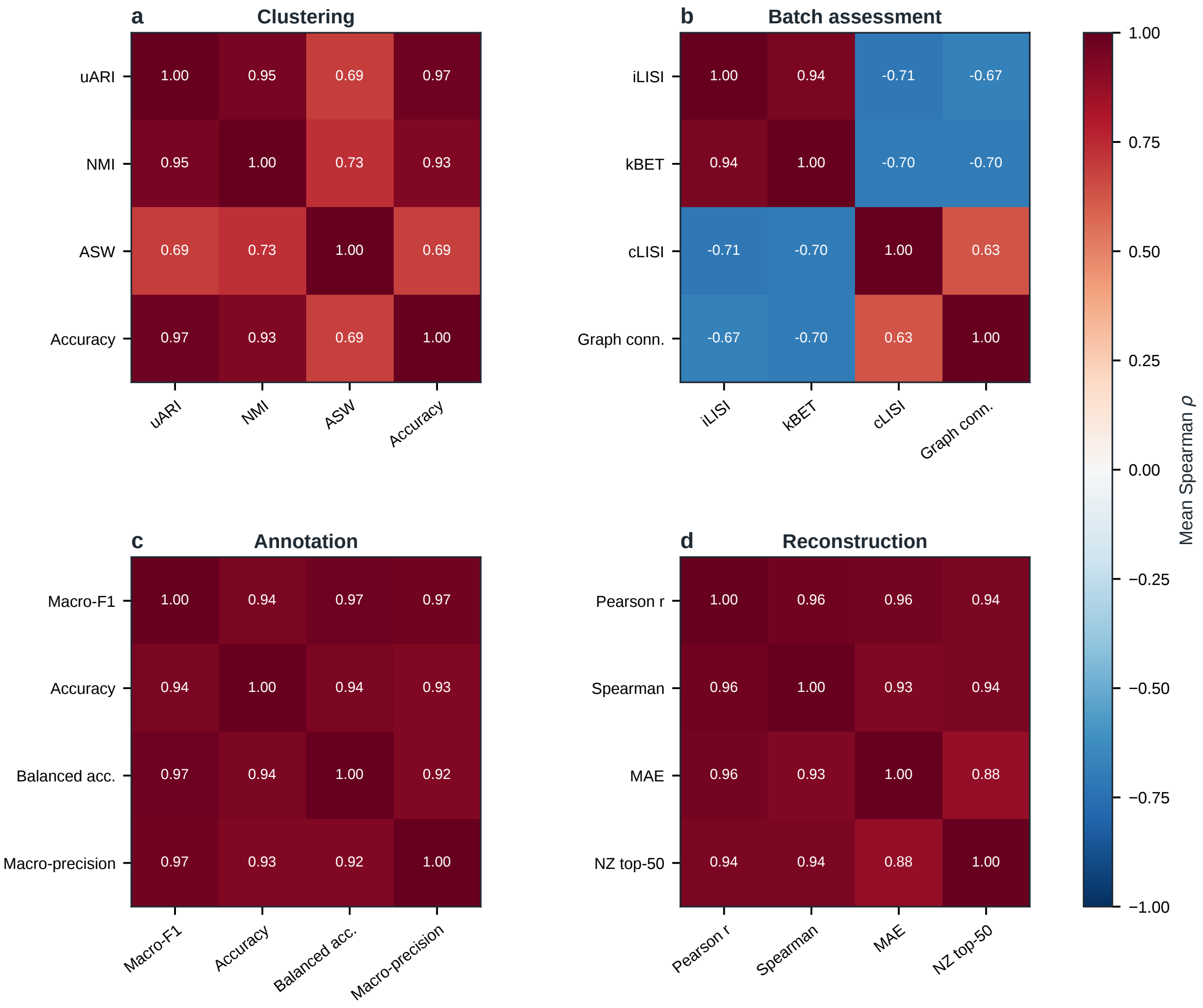


Figure S21: **Pairwise metric-rank agreement.**

### SUPPLEMENTAL TABLES

Table S1: **Donor-level split counts.**

| Panel | Dataset | Seed | Cells: train/val/test | Unit | Units: train/val/test |
|---|---|---|---|---|---|
| Human organs | bmmc | 41 | 21,378/2,534/6,004 | donors | 6/1/2 |
| Human organs | bmmc | 42 | 20,515/2,569/6,874 | donors | 6/1/2 |
| Human organs | bmmc | 43 | 20,569/2,564/6,867 | donors | 6/1/2 |
| Human organs | hlca | 41 | 20,159/3,664/6,177 | donors | 75/11/21 |
| Human organs | hlca | 42 | 21,079/3,285/5,636 | donors | 75/11/21 |
| Human organs | hlca | 43 | 21,347/3,702/4,951 | donors | 75/11/21 |
| Human organs | heart | 41 | 21,798/2,274/5,928 | donors | 10/1/3 |
| Human organs | heart | 42 | 20,092/2,145/7,763 | donors | 10/1/3 |
| Human organs | heart | 43 | 21,616/2,067/6,317 | donors | 10/1/3 |
| Human organs | kidney | 41 | 21,088/2,711/6,201 | donors | 32/4/9 |
| Human organs | kidney | 42 | 20,968/3,103/5,929 | donors | 32/4/9 |
| Human organs | kidney | 43 | 21,484/2,829/5,687 | donors | 32/4/9 |
| Human diseases | ms | 41 | 14,598/957/2,244 | donors | 6/1/2 |
| Human diseases | ms | 42 | 13,558/959/3,282 | donors | 6/1/2 |
| Human diseases | ms | 43 | 9,796/5,685/2,318 | donors | 6/1/2 |
| Human diseases | uc_epi | 41 | 21,676/3,047/5,277 | donors | 21/3/6 |
| Human diseases | uc_epi | 42 | 21,749/1,894/6,357 | donors | 21/3/6 |
| Human diseases | uc_epi | 43 | 22,237/2,840/4,923 | donors | 21/3/6 |
| Human diseases | covid_lung | 41 | 20,788/3,375/5,837 | donors | 19/3/5 |
| Human diseases | covid_lung | 42 | 21,193/3,542/5,265 | donors | 19/3/5 |
| Human diseases | covid_lung | 43 | 20,448/4,078/5,474 | donors | 19/3/5 |
| Human diseases | tnbc | 41 | 21,252/3,218/5,530 | donors | 71/10/20 |
| Human diseases | tnbc | 42 | 21,370/3,066/5,564 | donors | 71/10/20 |
| Human diseases | tnbc | 43 | 20,668/3,042/6,290 | donors | 71/10/20 |
| Mouse organs | mouse_marrow | 41 | 21,958/2,560/5,482 | donors | 12/2/3 |
| Mouse organs | mouse_marrow | 42 | 20,922/3,748/5,330 | donors | 12/2/3 |
| Mouse organs | mouse_marrow | 43 | 19,967/3,485/6,548 | donors | 12/2/3 |
| Mouse organs | mouse_lung | 41 | 20,371/1,870/2,299 | donors | 11/2/3 |
| Mouse organs | mouse_lung | 42 | 19,418/1,587/3,531 | donors | 11/2/3 |
| Mouse organs | mouse_lung | 43 | 19,522/2,167/2,851 | donors | 11/2/3 |
| Mouse organs | mouse_kidney | 41 | 14,611/1,691/5,345 | donors | 11/2/3 |
| Mouse organs | mouse_kidney | 42 | 17,213/1,561/2,745 | donors | 11/2/3 |
| Mouse organs | mouse_kidney | 43 | 14,834/1,699/5,111 | donors | 11/2/3 |
| Mouse organs | mouse_heart | 41 | 13,075/2,212/2,995 | donors | 19/3/5 |
| Mouse organs | mouse_heart | 42 | 15,214/1,685/1,383 | donors | 19/3/5 |
| Mouse organs | mouse_heart | 43 | 14,241/1,620/2,421 | donors | 19/3/5 |

Table S2: **Shared downstream probes.**

| Task | Probe | Shared architecture |
|---|---|---|
| Clustering and native batch assessment | Unsupervised graph | $\mathrm{PCA}(\min(256, n_{\mathrm{cells}}, d))$, 15-nearest-neighbor graph, Louvain clustering (`sc.tl.louvain`) at resolution 1.0, no post-hoc batch alignment |
| Cell-type annotation | Supervised MLP | $d \to 256 \to 256 \to C$ with batch normalization, ReLU, dropout 0.5, and $L_2$-normalized logits; Adam, $\mathrm{lr} = 10^{-3}$, batch size 64, up to 500 epochs, selected by minimum validation cross-entropy |
| HVG reconstruction | Supervised MLP | $d \to 512 \to 1024 \to 512 \to M$ ($M \leq 400$ training-split HVGs) with batch normalization, GELU, dropout 0.5, and nonnegative ReLU; Adam, $\mathrm{lr} = 2 \times 10^{-3}$, adaptive batch size 256–2,048, up to 150 epochs, MSE on $\log(1 + \mathrm{CP10k})$ expression, validation every 5 epochs with patience 4 |
| Perturbation prediction | Shared residual decoder | Unstandardized control-cell embedding concatenated with a 128-dimensional condition vector; $d + 128 \to 512 \to 1024 \to 512 \to G$ with batch normalization, GELU, and dropout 0.5; residual addition onto matched control expression, then nonnegative ReLU |

Table S3: **Checkpoint architecture and pretraining scope.**

| Checkpoint | Input[a] | Architecture / prior | Params (M) | Input extent[b] | Reported pretraining data[c,d] | Species |
|---|---|---|---|---|---|---|
| CellFM | A | ERetNet with linear retention | 797.4 | 2,048 | ∼100M cells[c] | Human |
| CellPLM | V | OmicsFormer with Gaussian-mixture prior | 65.8 | Full panel | >9M scRNA + >2M spatial[c] | Human[c] |
| GeneCompass | R+V | 6-layer BERT with regulatory-prior buffers | 18.1 | 2,048 | 101.8M cells (post-QC) | Human, mouse |
| Geneformer | R | 12-layer BERT with masked language modeling | 39.6 | 2,048 | ∼30M cells | Human |
| LangCell | R | Geneformer-based BERT with biomedical text head | 39.4 | 2,048 | ∼27.5M cell–text pairs | Human |
| Nicheformer | R+C | 12-layer transformer with scRNA and spatial context | 38.5 | 1,500 | >110M cells | Human, mouse |
| Tabula | V | FlashAttention-based tabular transformer (pancreas) | 13.2 | Tabular | 15M profiles[c] | Human[c] |
| TranscriptFormer | V | Autoregressive transformer with ESM-2 tokens | 429.9 | 1,023 | 57M cells | Human |
| UCE | S+P | 4-layer transformer with ESM-2 protein lookup table | 848.9 | 1,536 | >36M cells[c] | 8 species[c] |
| scFoundation | F+V | Asymmetric masked autoencoder with read-depth information | 119.3 | 19,264 | >50M cells | Human |
| scGPT | B | Generative masked transformer with 51 expression bins | 50.8 | 1,200 | ∼33M cells | Human |
| scLong | F+V | Dual Performer with Gene Ontology graph convolution | 1,006.1 | 27,875 | 48.0M unique cells | Human |
| scPRINT2 | A+C | Transformer with frozen gene encoder | 101.9 | 2,000 | >350M cells[c] | 16 species[c] |

[a]Input codes: A, gene-aligned; V, gene identity and values; R, ranked genes; C, context; S, sampled genes; P, protein-derived features; F, fixed gene panel; B, expression bins. [b]Sequence length or gene-panel size. [c]Family-level corpus or species estimate not separately verified for the evaluated checkpoint. [d]Primary corpus and checkpoint sources are listed in Notes on supplemental tables.

Table S4: **Evaluation datasets.**

| Panel | Code | Dataset | Cells | Genes | Source |
|---|---|---|---|---|---|
| Human organs | BM | bmmc | 30,000 | 13,953 | source |
| Human organs | HL | hlca | 30,000 | 27,402 | source |
| Human organs | AH | heart | 30,000 | 32,383 | source |
| Human organs | AK | kidney | 30,000 | 26,837 | source |
| Human diseases | MS | ms | 17,799 | 20,110 | source |
| Human diseases | UC | uc_epi | 30,000 | 20,028 | source |
| Human diseases | C19 | covid_lung | 30,000 | 33,523 | source |
| Human diseases | BC | tnbc | 30,000 | 32,354 | source |
| Mouse organs | MM | mouse_marrow | 30,000 | 17,943 | source |
| Mouse organs | ML | mouse_lung | 24,540 | 17,943 | source |
| Mouse organs | MH | mouse_heart | 18,282 | 16,669 | source |
| Mouse organs | MK | mouse_kidney | 21,647 | 17,943 | source |

Table S5: **Human-organ mean ranks.**

| Method | Clustering | Annotation | Batch Assessment | Reconstruction |
|---|---|---|---|---|
| CellFM | 14.771 | 13.646 | 8.312 | 13.729 |
| CellPLM | 6.104 | 11.792 | 7.083 | 12.292 |
| GeneCompass | 11.708 | 10.604 | 8.833 | 11.042 |
| Geneformer | 11.979 | 9.896 | 8.812 | 9.062 |
| LangCell | 8.208 | 7.125 | 10.438 | 6.125 |
| Nicheformer | 7.229 | 6.104 | 9.125 | 8.188 |
| Tabula | 15.375 | 16.000 | 8.375 | 16.000 |
| TranscriptFormer | 6.542 | 3.167 | 9.042 | 3.021 |
| UCE | 2.604 | 5.667 | 7.792 | 6.750 |
| scFoundation | 4.812 | 2.854 | 9.521 | 2.896 |
| scGPT | 3.917 | 4.562 | 8.167 | 8.521 |
| scLong | 14.854 | 15.000 | 9.146 | 15.000 |
| scPRINT2 | 10.333 | 9.146 | 9.229 | 5.833 |
| PCA | 5.083 | 4.146 | 6.042 | 1.000 |
| UMAP | 8.812 | 12.958 | 8.833 | 12.917 |
| scVI | 3.667 | 3.333 | 7.250 | 3.625 |

Table S6: **Human-disease mean ranks.**

| Method | Clustering | Annotation | Batch Assessment | Reconstruction |
|---|---|---|---|---|
| CellFM | 14.479 | 13.771 | 8.250 | 14.000 |
| CellPLM | 6.771 | 9.479 | 7.594 | 12.312 |
| GeneCompass | 9.812 | 9.083 | 7.688 | 10.812 |
| Geneformer | 10.604 | 9.323 | 7.875 | 8.688 |
| LangCell | 6.250 | 6.646 | 9.427 | 6.708 |
| Nicheformer | 6.583 | 5.698 | 7.760 | 7.062 |
| Tabula | 15.000 | 16.000 | 8.375 | 15.938 |
| TranscriptFormer | 6.500 | 3.552 | 8.719 | 4.333 |
| UCE | 3.771 | 5.000 | 7.625 | 6.333 |
| scFoundation | 5.740 | 2.156 | 9.688 | 4.000 |
| scGPT | 5.604 | 5.260 | 8.073 | 7.583 |
| scLong | 15.521 | 14.958 | 8.812 | 15.062 |
| scPRINT2 | 7.594 | 10.604 | 10.042 | 7.021 |
| PCA | 5.375 | 6.292 | 8.865 | 1.000 |
| UMAP | 8.979 | 13.146 | 8.052 | 12.229 |
| scVI | 7.417 | 5.031 | 9.156 | 2.917 |

Table S7: **Mouse-organ mean ranks.**

| Method | Clustering | Annotation | Batch Assessment | Reconstruction |
|---|---|---|---|---|
| CellFM | 14.646 | 12.812 | 8.042 | 12.875 |
| CellPLM | 8.208 | 11.208 | 8.917 | 11.583 |
| GeneCompass | 13.521 | 14.229 | 9.375 | 13.958 |
| Geneformer | 11.438 | 9.646 | 8.208 | 8.688 |
| LangCell | 6.042 | 5.365 | 9.146 | 4.750 |
| Nicheformer | 8.458 | 4.677 | 7.792 | 6.938 |
| Tabula | 15.062 | 16.000 | 8.500 | 15.958 |
| TranscriptFormer | 6.417 | 4.781 | 6.917 | 6.083 |
| UCE | 3.188 | 7.229 | 6.917 | 7.125 |
| scFoundation | 4.792 | 1.979 | 9.229 | 3.583 |
| scGPT | 5.479 | 7.281 | 8.479 | 8.271 |
| scLong | 14.583 | 14.708 | 9.042 | 14.750 |
| scPRINT2 | 9.812 | 6.365 | 10.479 | 5.771 |
| PCA | 2.500 | 5.188 | 8.083 | 1.312 |
| UMAP | 7.396 | 11.104 | 7.979 | 10.938 |
| scVI | 4.458 | 3.427 | 8.896 | 3.417 |

Table S8: **Human organs, clustering metrics.**

| Dataset | Method | uARI ↑ | NMI ↑ | ASW ↑ | ACC ↑ |
|---|---|---|---|---|---|
| bmmc | CellFM | 0.0670 | 0.2224 | 0.3525 | 0.1442 |
| bmmc | CellPLM | 0.4303 | 0.6921 | 0.4959 | 0.5025 |
| bmmc | GeneCompass | 0.2782 | 0.5802 | 0.4636 | 0.3577 |
| bmmc | Geneformer | 0.2989 | 0.6007 | 0.4732 | 0.3647 |
| bmmc | LangCell | 0.4176 | 0.6976 | 0.5207 | 0.4820 |
| bmmc | Nicheformer | 0.4447 | 0.6997 | 0.5258 | 0.5237 |
| bmmc | Tabula | 0.0133 | 0.0452 | 0.4520 | 0.0685 |
| bmmc | TranscriptFormer | 0.4572 | 0.6992 | 0.5336 | 0.5368 |
| bmmc | UCE | 0.5288 | 0.7558 | 0.5474 | 0.5979 |
| bmmc | scFoundation | 0.5293 | 0.7369 | 0.5456 | 0.5759 |
| bmmc | scGPT | 0.4661 | 0.7056 | 0.5288 | 0.5398 |
| bmmc | scLong | 0.0761 | 0.2404 | 0.3302 | 0.1520 |
| bmmc | scPRINT2 | 0.3366 | 0.6383 | 0.5096 | 0.4134 |
| bmmc | PCA | 0.4223 | 0.7260 | 0.5266 | 0.5248 |
| bmmc | UMAP | 0.2589 | 0.5166 | 0.4797 | 0.3958 |
| bmmc | scVI | 0.4564 | 0.7387 | 0.5388 | 0.5422 |
| hlca | CellFM | 0.0331 | 0.1035 | 0.3696 | 0.1203 |
| hlca | CellPLM | 0.5979 | 0.7624 | 0.5817 | 0.6982 |
| hlca | GeneCompass | 0.3659 | 0.6227 | 0.4940 | 0.4491 |
| hlca | Geneformer | 0.3252 | 0.6143 | 0.5051 | 0.3963 |
| hlca | LangCell | 0.3449 | 0.6415 | 0.5224 | 0.3995 |
| hlca | Nicheformer | 0.4471 | 0.7194 | 0.5513 | 0.5181 |
| hlca | Tabula | 0.0054 | 0.0150 | 0.4413 | 0.0838 |
| hlca | TranscriptFormer | 0.3732 | 0.6853 | 0.5443 | 0.4185 |
| hlca | UCE | 0.4866 | 0.7445 | 0.5866 | 0.5528 |
| hlca | scFoundation | 0.4102 | 0.7061 | 0.5621 | 0.4563 |
| hlca | scGPT | 0.6083 | 0.7630 | 0.5404 | 0.6780 |
| hlca | scLong | 0.0362 | 0.1236 | 0.3612 | 0.1084 |
| hlca | scPRINT2 | 0.4002 | 0.6609 | 0.5230 | 0.4712 |
| hlca | PCA | 0.6650 | 0.7925 | 0.5485 | 0.7383 |
| hlca | UMAP | 0.5300 | 0.7234 | 0.6134 | 0.6032 |
| hlca | scVI | 0.6180 | 0.7956 | 0.5689 | 0.6718 |

*Continued on next page*

Table S8: **Human organs, clustering metrics.** *(continued)*

| Dataset | Method | uARI ↑ | NMI ↑ | ASW ↑ | ACC ↑ |
|---|---|---|---|---|---|
| heart | CellFM | 0.0263 | 0.1157 | 0.3979 | 0.1027 |
| heart | CellPLM | 0.4168 | 0.6551 | 0.5279 | 0.4957 |
| heart | GeneCompass | 0.3520 | 0.6117 | 0.4755 | 0.4174 |
| heart | Geneformer | 0.3359 | 0.6028 | 0.4678 | 0.3950 |
| heart | LangCell | 0.4240 | 0.6555 | 0.5189 | 0.4927 |
| heart | Nicheformer | 0.4006 | 0.6746 | 0.5222 | 0.4875 |
| heart | Tabula | 0.0108 | 0.0332 | 0.4517 | 0.0771 |
| heart | TranscriptFormer | 0.3527 | 0.6618 | 0.5329 | 0.3943 |
| heart | UCE | 0.4575 | 0.7178 | 0.5628 | 0.5055 |
| heart | scFoundation | 0.4991 | 0.7105 | 0.5585 | 0.5636 |
| heart | scGPT | 0.4814 | 0.7173 | 0.5381 | 0.5568 |
| heart | scLong | 0.0292 | 0.1097 | 0.3772 | 0.0937 |
| heart | scPRINT2 | 0.3490 | 0.6194 | 0.5043 | 0.4274 |
| heart | PCA | 0.4389 | 0.6883 | 0.5190 | 0.5353 |
| heart | UMAP | 0.3208 | 0.5557 | 0.5072 | 0.4422 |
| heart | scVI | 0.5137 | 0.7385 | 0.5494 | 0.6007 |
| kidney | CellFM | 0.0395 | 0.0932 | 0.4165 | 0.1164 |
| kidney | CellPLM | 0.4258 | 0.5682 | 0.5587 | 0.4888 |
| kidney | GeneCompass | 0.3012 | 0.4109 | 0.5069 | 0.4008 |
| kidney | Geneformer | 0.2952 | 0.4406 | 0.5149 | 0.4121 |
| kidney | LangCell | 0.4698 | 0.6146 | 0.5328 | 0.5654 |
| kidney | Nicheformer | 0.3416 | 0.5211 | 0.5414 | 0.4288 |
| kidney | Tabula | 0.0095 | 0.0219 | 0.4731 | 0.1241 |
| kidney | TranscriptFormer | 0.4963 | 0.6844 | 0.5695 | 0.5634 |
| kidney | UCE | 0.5844 | 0.7106 | 0.5787 | 0.6719 |
| kidney | scFoundation | 0.4046 | 0.5780 | 0.5406 | 0.5020 |
| kidney | scGPT | 0.5152 | 0.6482 | 0.5438 | 0.5934 |
| kidney | scLong | 0.0312 | 0.0945 | 0.4228 | 0.0978 |
| kidney | scPRINT2 | 0.2537 | 0.4619 | 0.5211 | 0.3528 |
| kidney | PCA | 0.4046 | 0.6433 | 0.5427 | 0.4660 |
| kidney | UMAP | 0.3365 | 0.5891 | 0.5856 | 0.4005 |
| kidney | scVI | 0.3637 | 0.6270 | 0.5461 | 0.4206 |

Table S9: **Human organs, annotation metrics.**

| Dataset | Method | ACC ↑ | Macro-F1 ↑ | Balanced ACC ↑ | Macro-precision ↑ |
|---|---|---|---|---|---|
| bmmc | CellFM | 0.5705 | 0.4272 | 0.4671 | 0.4266 |
| bmmc | CellPLM | 0.6116 | 0.4515 | 0.5068 | 0.4466 |
| bmmc | GeneCompass | 0.6304 | 0.4802 | 0.5224 | 0.4906 |
| bmmc | Geneformer | 0.6593 | 0.4981 | 0.5410 | 0.5042 |
| bmmc | LangCell | 0.6707 | 0.5123 | 0.5557 | 0.5203 |
| bmmc | Nicheformer | 0.6843 | 0.5235 | 0.5664 | 0.5316 |
| bmmc | Tabula | 0.0859 | 0.0361 | 0.0559 | 0.0359 |
| bmmc | TranscriptFormer | 0.6847 | 0.5280 | 0.5712 | 0.5464 |
| bmmc | UCE | 0.6736 | 0.5206 | 0.5614 | 0.5361 |
| bmmc | scFoundation | 0.6878 | 0.5309 | 0.5783 | 0.5435 |
| bmmc | scGPT | 0.6728 | 0.5107 | 0.5560 | 0.5183 |
| bmmc | scLong | 0.4611 | 0.3210 | 0.3529 | 0.3358 |
| bmmc | scPRINT2 | 0.6657 | 0.5059 | 0.5453 | 0.5165 |
| bmmc | PCA | 0.7172 | 0.5818 | 0.6146 | 0.6027 |
| bmmc | UMAP | 0.5103 | 0.3748 | 0.4021 | 0.3906 |
| bmmc | scVI | 0.6806 | 0.5365 | 0.5698 | 0.5440 |
| hlca | CellFM | 0.7527 | 0.6265 | 0.6218 | 0.6513 |
| hlca | CellPLM | 0.8863 | 0.8403 | 0.8453 | 0.8507 |
| hlca | GeneCompass | 0.8854 | 0.8456 | 0.8541 | 0.8480 |
| hlca | Geneformer | 0.8980 | 0.8498 | 0.8531 | 0.8592 |
| hlca | LangCell | 0.9158 | 0.8610 | 0.8618 | 0.8695 |
| hlca | Nicheformer | 0.9157 | 0.8828 | 0.8887 | 0.8975 |
| hlca | Tabula | 0.1070 | 0.0403 | 0.0664 | 0.0495 |
| hlca | TranscriptFormer | 0.9269 | 0.8701 | 0.8691 | 0.8776 |
| hlca | UCE | 0.9205 | 0.8645 | 0.8637 | 0.8726 |
| hlca | scFoundation | 0.9342 | 0.8993 | 0.9058 | 0.8983 |
| hlca | scGPT | 0.9264 | 0.8885 | 0.8921 | 0.8874 |
| hlca | scLong | 0.5687 | 0.4438 | 0.4501 | 0.4663 |
| hlca | scPRINT2 | 0.9017 | 0.8513 | 0.8514 | 0.8624 |
| hlca | PCA | 0.9226 | 0.8616 | 0.8638 | 0.8700 |
| hlca | UMAP | 0.8557 | 0.7866 | 0.7902 | 0.7992 |
| hlca | scVI | 0.9310 | 0.8985 | 0.8973 | 0.9083 |

*Continued on next page*

Table S9: **Human organs, annotation metrics.** *(continued)*

| Dataset | Method | ACC ↑ | Macro-F1 ↑ | Balanced ACC ↑ | Macro-precision ↑ |
|---|---|---|---|---|---|
| heart | CellFM | 0.5646 | 0.5085 | 0.5156 | 0.5459 |
| heart | CellPLM | 0.7063 | 0.6697 | 0.6846 | 0.7115 |
| heart | GeneCompass | 0.7212 | 0.6926 | 0.6951 | 0.7234 |
| heart | Geneformer | 0.7250 | 0.6919 | 0.6961 | 0.7357 |
| heart | LangCell | 0.7501 | 0.7192 | 0.7212 | 0.7423 |
| heart | Nicheformer | 0.7725 | 0.7437 | 0.7451 | 0.7708 |
| heart | Tabula | 0.0845 | 0.0486 | 0.0680 | 0.0501 |
| heart | TranscriptFormer | 0.8108 | 0.7858 | 0.7845 | 0.8048 |
| heart | UCE | 0.7915 | 0.7566 | 0.7623 | 0.7845 |
| heart | scFoundation | 0.7965 | 0.7684 | 0.7691 | 0.7903 |
| heart | scGPT | 0.7887 | 0.7599 | 0.7605 | 0.7868 |
| heart | scLong | 0.3647 | 0.2965 | 0.3231 | 0.3448 |
| heart | scPRINT2 | 0.7352 | 0.6901 | 0.6930 | 0.7258 |
| heart | PCA | 0.7809 | 0.7477 | 0.7492 | 0.7677 |
| heart | UMAP | 0.5781 | 0.5107 | 0.5235 | 0.5517 |
| heart | scVI | 0.7848 | 0.7494 | 0.7523 | 0.7826 |
| kidney | CellFM | 0.7209 | 0.6476 | 0.6396 | 0.6784 |
| kidney | CellPLM | 0.7906 | 0.7463 | 0.7467 | 0.7588 |
| kidney | GeneCompass | 0.8100 | 0.7730 | 0.7751 | 0.7779 |
| kidney | Geneformer | 0.8157 | 0.7720 | 0.7687 | 0.7807 |
| kidney | LangCell | 0.8536 | 0.8103 | 0.8072 | 0.8208 |
| kidney | Nicheformer | 0.8488 | 0.8010 | 0.7981 | 0.8117 |
| kidney | Tabula | 0.1606 | 0.0867 | 0.1222 | 0.1103 |
| kidney | TranscriptFormer | 0.8697 | 0.8278 | 0.8269 | 0.8464 |
| kidney | UCE | 0.8539 | 0.8130 | 0.8122 | 0.8255 |
| kidney | scFoundation | 0.8634 | 0.8220 | 0.8204 | 0.8321 |
| kidney | scGPT | 0.8584 | 0.8176 | 0.8143 | 0.8259 |
| kidney | scLong | 0.5777 | 0.4500 | 0.4640 | 0.5490 |
| kidney | scPRINT2 | 0.8380 | 0.7921 | 0.7878 | 0.8033 |
| kidney | PCA | 0.8630 | 0.8213 | 0.8217 | 0.8286 |
| kidney | UMAP | 0.8191 | 0.7566 | 0.7465 | 0.7844 |
| kidney | scVI | 0.8661 | 0.8263 | 0.8253 | 0.8361 |

Table S10: **Human organs, batch-assessment metrics.**

| Dataset | Method | Scaled iLISI ↑ | kBET ↑ | Scaled cLISI ↑ | Graph conn. ↑ |
|---|---|---|---|---|---|
| bmmc | CellFM | 0.3378 | 0.3625 | 0.8863 | 0.3964 |
| bmmc | CellPLM | 0.3392 | 0.4854 | 0.9745 | 0.7759 |
| bmmc | GeneCompass | 0.2280 | 0.2217 | 0.9607 | 0.8772 |
| bmmc | Geneformer | 0.2206 | 0.2088 | 0.9665 | 0.8829 |
| bmmc | LangCell | 0.1703 | 0.1376 | 0.9777 | 0.9366 |
| bmmc | Nicheformer | 0.2063 | 0.1711 | 0.9790 | 0.9326 |
| bmmc | Tabula | 0.4464 | 0.7607 | 0.7978 | 0.0635 |
| bmmc | TranscriptFormer | 0.2101 | 0.1598 | 0.9793 | 0.9387 |
| bmmc | UCE | 0.3122 | 0.3019 | 0.9814 | 0.9318 |
| bmmc | scFoundation | 0.1557 | 0.1126 | 0.9824 | 0.9393 |
| bmmc | scGPT | 0.1854 | 0.1570 | 0.9765 | 0.9150 |
| bmmc | scLong | 0.3326 | 0.2986 | 0.8693 | 0.2406 |
| bmmc | scPRINT2 | 0.1831 | 0.1541 | 0.9672 | 0.9341 |
| bmmc | PCA | 0.2678 | 0.2058 | 0.9769 | 0.9395 |
| bmmc | UMAP | 0.2272 | 0.2709 | 0.9535 | 0.6365 |
| bmmc | scVI | 0.2288 | 0.2036 | 0.9788 | 0.9483 |
| hlca | CellFM | 0.2026 | 0.4727 | 0.8195 | 0.4530 |
| hlca | CellPLM | 0.1365 | 0.2924 | 0.9933 | 0.9018 |
| hlca | GeneCompass | 0.0971 | 0.1917 | 0.9634 | 0.9647 |
| hlca | Geneformer | 0.0703 | 0.1346 | 0.9725 | 0.9509 |
| hlca | LangCell | 0.0133 | 0.0964 | 0.9845 | 0.9486 |
| hlca | Nicheformer | 0.0518 | 0.1252 | 0.9924 | 0.9872 |
| hlca | Tabula | 0.2587 | 0.7856 | 0.6868 | 0.1709 |
| hlca | TranscriptFormer | 0.0148 | 0.1053 | 0.9961 | 0.9893 |
| hlca | UCE | 0.0406 | 0.1164 | 0.9979 | 0.9888 |
| hlca | scFoundation | 0.0246 | 0.1072 | 0.9958 | 0.9897 |
| hlca | scGPT | 0.0754 | 0.1751 | 0.9932 | 0.9865 |
| hlca | scLong | 0.2125 | 0.4674 | 0.7714 | 0.2145 |
| hlca | scPRINT2 | 0.0625 | 0.1564 | 0.9834 | 0.9859 |
| hlca | PCA | 0.1035 | 0.2023 | 0.9959 | 0.9887 |
| hlca | UMAP | 0.1121 | 0.2391 | 0.9966 | 0.8786 |
| hlca | scVI | 0.0526 | 0.1410 | 0.9964 | 0.9955 |
| heart | CellFM | 0.3775 | 0.3582 | 0.7720 | 0.3228 |
| heart | CellPLM | 0.2602 | 0.1646 | 0.9705 | 0.6356 |
| heart | GeneCompass | 0.2323 | 0.0877 | 0.9460 | 0.8085 |
| heart | Geneformer | 0.2619 | 0.1297 | 0.9425 | 0.8168 |
| heart | LangCell | 0.2094 | 0.0541 | 0.9560 | 0.8941 |
| heart | Nicheformer | 0.2179 | 0.0720 | 0.9687 | 0.7320 |
| heart | Tabula | 0.4688 | 0.8061 | 0.6925 | 0.1331 |
| heart | TranscriptFormer | 0.1458 | 0.0289 | 0.9843 | 0.8072 |
| heart | UCE | 0.1882 | 0.0481 | 0.9851 | 0.7305 |
| heart | scFoundation | 0.1999 | 0.0491 | 0.9786 | 0.8059 |
| heart | scGPT | 0.2068 | 0.0576 | 0.9749 | 0.9012 |
| heart | scLong | 0.3610 | 0.3742 | 0.7507 | 0.1613 |
| heart | scPRINT2 | 0.2230 | 0.0779 | 0.9524 | 0.8577 |
| heart | PCA | 0.2427 | 0.0933 | 0.9671 | 0.9128 |
| heart | UMAP | 0.2306 | 0.1412 | 0.9560 | 0.7017 |
| heart | scVI | 0.1910 | 0.0598 | 0.9802 | 0.9543 |

*Continued on next page*

Table S10: **Human organs, batch-assessment metrics.** *(continued)*

| Dataset | Method | Scaled iLISI ↑ | kBET ↑ | Scaled cLISI ↑ | Graph conn. ↑ |
|---|---|---|---|---|---|
| kidney | CellFM | 0.7891 | 0.5940 | 0.6659 | 0.3958 |
| kidney | CellPLM | 0.5025 | 0.3559 | 0.9563 | 0.8134 |
| kidney | GeneCompass | 0.6070 | 0.4137 | 0.8317 | 0.9527 |
| kidney | Geneformer | 0.5809 | 0.3883 | 0.8551 | 0.9668 |
| kidney | LangCell | 0.3616 | 0.2823 | 0.9777 | 0.9662 |
| kidney | Nicheformer | 0.3676 | 0.2931 | 0.9380 | 0.9715 |
| kidney | Tabula | 0.8250 | 0.8584 | 0.5684 | 0.5430 |
| kidney | TranscriptFormer | 0.1684 | 0.2235 | 0.9964 | 0.9766 |
| kidney | UCE | 0.3173 | 0.2542 | 0.9929 | 0.9721 |
| kidney | scFoundation | 0.2421 | 0.2253 | 0.9715 | 0.9737 |
| kidney | scGPT | 0.3736 | 0.2846 | 0.9792 | 0.9780 |
| kidney | scLong | 0.7149 | 0.5155 | 0.6594 | 0.2892 |
| kidney | scPRINT2 | 0.4182 | 0.3290 | 0.9484 | 0.9704 |
| kidney | PCA | 0.3849 | 0.2651 | 0.9916 | 0.9773 |
| kidney | UMAP | 0.2942 | 0.2594 | 0.9910 | 0.7972 |
| kidney | scVI | 0.1326 | 0.1626 | 0.9900 | 0.9792 |

Table S11: **Human organs, reconstruction metrics ($\log(1 + \mathrm{CP10k})$).**

| Dataset | Method | Pearson $r$ ↑ | Spearman ↑ | MAE ↓ | NZ top-50 ↑ |
|---|---|---|---|---|---|
| bmmc | CellFM | 0.7108 | 0.4869 | 0.1097 | 0.5429 |
| bmmc | CellPLM | 0.7427 | 0.5101 | 0.1015 | 0.5709 |
| bmmc | GeneCompass | 0.7516 | 0.5200 | 0.0988 | 0.5804 |
| bmmc | Geneformer | 0.7609 | 0.5281 | 0.0952 | 0.5870 |
| bmmc | LangCell | 0.7703 | 0.5340 | 0.0933 | 0.5911 |
| bmmc | Nicheformer | 0.7650 | 0.5292 | 0.0951 | 0.5907 |
| bmmc | Tabula | 0.4597 | 0.2580 | 0.1768 | 0.3049 |
| bmmc | TranscriptFormer | 0.7783 | 0.5403 | 0.0922 | 0.6029 |
| bmmc | UCE | 0.7649 | 0.5315 | 0.0963 | 0.5935 |
| bmmc | scFoundation | 0.7785 | 0.5387 | 0.0924 | 0.6028 |
| bmmc | scGPT | 0.7636 | 0.5271 | 0.0958 | 0.5883 |
| bmmc | scLong | 0.6575 | 0.4414 | 0.1280 | 0.4946 |
| bmmc | scPRINT2 | 0.7721 | 0.5288 | 0.0926 | 0.5859 |
| bmmc | PCA | 0.8168 | 0.5951 | 0.0801 | 0.6435 |
| bmmc | UMAP | 0.7034 | 0.4785 | 0.1098 | 0.5328 |
| bmmc | scVI | 0.7748 | 0.5434 | 0.0919 | 0.5989 |
| bmmc | *Train mean* | 0.4373 | 0.2391 | 0.2112 | 0.2801 |
| bmmc | *Train median* | 0.3638 | 0.2123 | 0.1303 | 0.0636 |
| bmmc | *All zero* | 0.0000 | 0.0000 | 0.1379 | 0.0000 |
| hlca | CellFM | 0.5723 | 0.3915 | 0.0549 | 0.4382 |
| hlca | CellPLM | 0.6512 | 0.4651 | 0.0477 | 0.5000 |
| hlca | GeneCompass | 0.6718 | 0.5026 | 0.0430 | 0.5115 |
| hlca | Geneformer | 0.6926 | 0.5182 | 0.0406 | 0.5241 |
| hlca | LangCell | 0.7179 | 0.5378 | 0.0387 | 0.5398 |
| hlca | Nicheformer | 0.6948 | 0.5237 | 0.0400 | 0.5234 |
| hlca | Tabula | 0.3750 | 0.2289 | 0.0767 | 0.2863 |
| hlca | TranscriptFormer | 0.7239 | 0.5552 | 0.0376 | 0.5485 |
| hlca | UCE | 0.7088 | 0.5387 | 0.0400 | 0.5465 |
| hlca | scFoundation | 0.7385 | 0.5572 | 0.0370 | 0.5650 |
| hlca | scGPT | 0.7022 | 0.5280 | 0.0401 | 0.5335 |
| hlca | scLong | 0.5018 | 0.3247 | 0.0641 | 0.3815 |
| hlca | scPRINT2 | 0.7206 | 0.5362 | 0.0376 | 0.5267 |
| hlca | PCA | 0.8382 | 0.7039 | 0.0267 | 0.6781 |
| hlca | UMAP | 0.6548 | 0.4675 | 0.0449 | 0.4990 |
| hlca | scVI | 0.7384 | 0.5548 | 0.0372 | 0.5646 |
| hlca | *Train mean* | 0.3818 | 0.1956 | 0.0933 | 0.2909 |
| hlca | *Train median* | 0.2343 | 0.2112 | 0.0540 | 0.0823 |
| hlca | *All zero* | 0.0000 | 0.0000 | 0.0551 | 0.0000 |

*Continued on next page*

Table S11: **Human organs, reconstruction metrics ($\log(1 + \mathrm{CP10k})$).** *(continued)*

| Dataset | Method | Pearson $r \uparrow$ | Spearman $\uparrow$ | MAE $\downarrow$ | NZ top-50 $\uparrow$ |
|---|---|---|---|---|---|
| heart | CellFM | 0.5349 | 0.3853 | 0.1239 | 0.4371 |
| heart | CellPLM | 0.6055 | 0.4356 | 0.1118 | 0.4951 |
| heart | GeneCompass | 0.6143 | 0.4568 | 0.1040 | 0.4993 |
| heart | Geneformer | 0.6376 | 0.4767 | 0.0986 | 0.5201 |
| heart | LangCell | 0.6472 | 0.4813 | 0.0962 | 0.5217 |
| heart | Nicheformer | 0.6411 | 0.4768 | 0.0979 | 0.5207 |
| heart | Tabula | 0.2703 | 0.2098 | 0.1813 | 0.2184 |
| heart | TranscriptFormer | 0.6711 | 0.4992 | 0.0926 | 0.5468 |
| heart | UCE | 0.6434 | 0.4843 | 0.0988 | 0.5270 |
| heart | scFoundation | 0.6681 | 0.4938 | 0.0939 | 0.5423 |
| heart | scGPT | 0.6372 | 0.4727 | 0.1002 | 0.5161 |
| heart | scLong | 0.4206 | 0.2965 | 0.1571 | 0.3412 |
| heart | scPRINT2 | 0.6598 | 0.4850 | 0.0936 | 0.5248 |
| heart | PCA | 0.7780 | 0.6194 | 0.0715 | 0.6721 |
| heart | UMAP | 0.5500 | 0.3951 | 0.1145 | 0.4442 |
| heart | scVI | 0.6563 | 0.4918 | 0.0947 | 0.5317 |
| heart | *Train mean* | 0.2436 | 0.1929 | 0.1966 | 0.1904 |
| heart | *Train median* | 0.0000 | 0.0000 | 0.1135 | 0.0000 |
| heart | *All zero* | 0.0000 | 0.0000 | 0.1135 | 0.0000 |
| kidney | CellFM | 0.6185 | 0.4640 | 0.0719 | 0.5086 |
| kidney | CellPLM | 0.6990 | 0.5174 | 0.0636 | 0.5708 |
| kidney | GeneCompass | 0.7228 | 0.5500 | 0.0575 | 0.5851 |
| kidney | Geneformer | 0.7407 | 0.5706 | 0.0535 | 0.5976 |
| kidney | LangCell | 0.7757 | 0.5884 | 0.0507 | 0.6152 |
| kidney | Nicheformer | 0.7510 | 0.5742 | 0.0529 | 0.6015 |
| kidney | Tabula | 0.3306 | 0.2432 | 0.1197 | 0.2820 |
| kidney | TranscriptFormer | 0.7960 | 0.6137 | 0.0477 | 0.6473 |
| kidney | UCE | 0.7603 | 0.5803 | 0.0535 | 0.6176 |
| kidney | scFoundation | 0.8048 | 0.6075 | 0.0475 | 0.6437 |
| kidney | scGPT | 0.7538 | 0.5752 | 0.0535 | 0.6080 |
| kidney | scLong | 0.5209 | 0.3854 | 0.0876 | 0.4409 |
| kidney | scPRINT2 | 0.7783 | 0.5843 | 0.0491 | 0.6047 |
| kidney | PCA | 0.8850 | 0.7519 | 0.0344 | 0.7655 |
| kidney | UMAP | 0.7057 | 0.5235 | 0.0604 | 0.5676 |
| kidney | scVI | 0.7927 | 0.6089 | 0.0493 | 0.6393 |
| kidney | *Train mean* | 0.3217 | 0.2220 | 0.1305 | 0.2787 |
| kidney | *Train median* | 0.0426 | 0.0444 | 0.0753 | 0.0142 |
| kidney | *All zero* | 0.0000 | 0.0000 | 0.0752 | 0.0000 |

Table S12: **Human diseases, clustering metrics.**

| Dataset | Method | uARI ↑ | NMI ↑ | ASW ↑ | ACC ↑ |
|---|---|---|---|---|---|
| ms | CellFM | 0.0341 | 0.1322 | 0.4073 | 0.1252 |
| ms | CellPLM | 0.1725 | 0.5448 | 0.6712 | 0.3058 |
| ms | GeneCompass | 0.3293 | 0.6111 | 0.5961 | 0.4767 |
| ms | Geneformer | 0.3449 | 0.6031 | 0.5369 | 0.4215 |
| ms | LangCell | 0.3794 | 0.6357 | 0.5928 | 0.5061 |
| ms | Nicheformer | 0.2470 | 0.5895 | 0.6150 | 0.3777 |
| ms | Tabula | 0.0367 | 0.0662 | 0.4361 | 0.2136 |
| ms | TranscriptFormer | 0.2019 | 0.5743 | 0.5614 | 0.3383 |
| ms | UCE | 0.3043 | 0.6029 | 0.6111 | 0.4099 |
| ms | scFoundation | 0.2616 | 0.6097 | 0.6334 | 0.3838 |
| ms | scGPT | 0.2301 | 0.5845 | 0.6015 | 0.3616 |
| ms | scLong | 0.0257 | 0.1296 | 0.3885 | 0.1026 |
| ms | scPRINT2 | 0.3961 | 0.6199 | 0.5668 | 0.4699 |
| ms | PCA | 0.4811 | 0.6618 | 0.5515 | 0.5703 |
| ms | UMAP | 0.1724 | 0.4550 | 0.6155 | 0.2723 |
| ms | scVI | 0.3023 | 0.6040 | 0.5670 | 0.4216 |
| uc_epi | CellFM | 0.0628 | 0.1962 | 0.3932 | 0.1338 |
| uc_epi | CellPLM | 0.1927 | 0.3576 | 0.4740 | 0.3360 |
| uc_epi | GeneCompass | 0.1322 | 0.2772 | 0.4706 | 0.2818 |
| uc_epi | Geneformer | 0.1314 | 0.2814 | 0.4758 | 0.2864 |
| uc_epi | LangCell | 0.1587 | 0.3319 | 0.5024 | 0.3078 |
| uc_epi | Nicheformer | 0.1791 | 0.3632 | 0.4943 | 0.3185 |
| uc_epi | Tabula | 0.0392 | 0.0964 | 0.4549 | 0.1421 |
| uc_epi | TranscriptFormer | 0.2211 | 0.4270 | 0.4858 | 0.3715 |
| uc_epi | UCE | 0.2246 | 0.4288 | 0.5092 | 0.3786 |
| uc_epi | scFoundation | 0.1623 | 0.3490 | 0.5041 | 0.3150 |
| uc_epi | scGPT | 0.1969 | 0.3806 | 0.4956 | 0.3505 |
| uc_epi | scLong | 0.0594 | 0.1842 | 0.3715 | 0.1372 |
| uc_epi | scPRINT2 | 0.1926 | 0.3428 | 0.5013 | 0.3318 |
| uc_epi | PCA | 0.2773 | 0.4408 | 0.5068 | 0.4576 |
| uc_epi | UMAP | 0.2069 | 0.4127 | 0.4692 | 0.3456 |
| uc_epi | scVI | 0.1564 | 0.3212 | 0.5177 | 0.3036 |

*Continued on next page*

Table S12: **Human diseases, clustering metrics.** *(continued)*

| Dataset | Method | uARI ↑ | NMI ↑ | ASW ↑ | ACC ↑ |
|---|---|---|---|---|---|
| covid_lung | CellFM | 0.0197 | 0.0813 | 0.3897 | 0.0880 |
| covid_lung | CellPLM | 0.4060 | 0.6562 | 0.5281 | 0.4853 |
| covid_lung | GeneCompass | 0.3455 | 0.6238 | 0.4946 | 0.4115 |
| covid_lung | Geneformer | 0.3247 | 0.6137 | 0.4918 | 0.3835 |
| covid_lung | LangCell | 0.4028 | 0.6847 | 0.5124 | 0.4838 |
| covid_lung | Nicheformer | 0.4106 | 0.6820 | 0.5217 | 0.4732 |
| covid_lung | Tabula | 0.0124 | 0.0362 | 0.4430 | 0.0827 |
| covid_lung | TranscriptFormer | 0.4653 | 0.7172 | 0.5139 | 0.5626 |
| covid_lung | UCE | 0.4959 | 0.7319 | 0.5547 | 0.5887 |
| covid_lung | scFoundation | 0.4800 | 0.7238 | 0.5465 | 0.5411 |
| covid_lung | scGPT | 0.4328 | 0.7117 | 0.5345 | 0.5167 |
| covid_lung | scLong | 0.0123 | 0.0605 | 0.3434 | 0.0678 |
| covid_lung | scPRINT2 | 0.3230 | 0.5924 | 0.4997 | 0.3958 |
| covid_lung | PCA | 0.4186 | 0.7026 | 0.5189 | 0.4920 |
| covid_lung | UMAP | 0.3893 | 0.6308 | 0.5213 | 0.5052 |
| covid_lung | scVI | 0.4513 | 0.7180 | 0.5395 | 0.5327 |
| tnbc | CellFM | 0.0885 | 0.2306 | 0.4543 | 0.1696 |
| tnbc | CellPLM | 0.5410 | 0.7791 | 0.7070 | 0.6056 |
| tnbc | GeneCompass | 0.4656 | 0.7309 | 0.5342 | 0.4632 |
| tnbc | Geneformer | 0.4597 | 0.7367 | 0.5335 | 0.5048 |
| tnbc | LangCell | 0.5919 | 0.7753 | 0.5568 | 0.6747 |
| tnbc | Nicheformer | 0.5319 | 0.7711 | 0.6225 | 0.6081 |
| tnbc | Tabula | 0.0136 | 0.0348 | 0.4704 | 0.1536 |
| tnbc | TranscriptFormer | 0.5152 | 0.7571 | 0.5796 | 0.5852 |
| tnbc | UCE | 0.5268 | 0.7664 | 0.5967 | 0.5555 |
| tnbc | scFoundation | 0.5017 | 0.7562 | 0.5914 | 0.5738 |
| tnbc | scGPT | 0.6077 | 0.8064 | 0.5650 | 0.6646 |
| tnbc | scLong | 0.0663 | 0.1952 | 0.4518 | 0.1341 |
| tnbc | scPRINT2 | 0.5525 | 0.7547 | 0.5420 | 0.6006 |
| tnbc | PCA | 0.4670 | 0.7219 | 0.5806 | 0.5267 |
| tnbc | UMAP | 0.2705 | 0.6059 | 0.6550 | 0.3572 |
| tnbc | scVI | 0.4453 | 0.7214 | 0.5566 | 0.5119 |

Table S13: **Human diseases, annotation metrics.**

| Dataset | Method | ACC ↑ | Macro-F1 ↑ | Balanced ACC ↑ | Macro-precision ↑ |
|---|---|---|---|---|---|
| ms | CellFM | 0.8924 | 0.5990 | 0.6169 | 0.6474 |
| ms | CellPLM | 0.9504 | 0.8775 | 0.8614 | 0.9017 |
| ms | GeneCompass | 0.9573 | 0.8530 | 0.8666 | 0.8585 |
| ms | Geneformer | 0.9549 | 0.8494 | 0.8401 | 0.8724 |
| ms | LangCell | 0.9645 | 0.8565 | 0.8453 | 0.8775 |
| ms | Nicheformer | 0.9617 | 0.8427 | 0.8534 | 0.8549 |
| ms | Tabula | 0.5516 | 0.1353 | 0.1599 | 0.1288 |
| ms | TranscriptFormer | 0.9642 | 0.8274 | 0.8402 | 0.8299 |
| ms | UCE | 0.9621 | 0.8554 | 0.8432 | 0.8758 |
| ms | scFoundation | 0.9692 | 0.9177 | 0.9195 | 0.9248 |
| ms | scGPT | 0.9634 | 0.8517 | 0.8787 | 0.8432 |
| ms | scLong | 0.8390 | 0.4333 | 0.4397 | 0.4884 |
| ms | scPRINT2 | 0.9496 | 0.7809 | 0.7775 | 0.8102 |
| ms | PCA | 0.9482 | 0.7700 | 0.7661 | 0.7877 |
| ms | UMAP | 0.8893 | 0.5538 | 0.5762 | 0.5951 |
| ms | scVI | 0.9565 | 0.8268 | 0.8531 | 0.8336 |
| uc_epi | CellFM | 0.5998 | 0.5382 | 0.5376 | 0.5767 |
| uc_epi | CellPLM | 0.6851 | 0.6547 | 0.6543 | 0.6763 |
| uc_epi | GeneCompass | 0.6732 | 0.6493 | 0.6411 | 0.6750 |
| uc_epi | Geneformer | 0.7106 | 0.6911 | 0.6767 | 0.7261 |
| uc_epi | LangCell | 0.7359 | 0.7300 | 0.7168 | 0.7581 |
| uc_epi | Nicheformer | 0.7403 | 0.7278 | 0.7178 | 0.7491 |
| uc_epi | Tabula | 0.1899 | 0.0863 | 0.1273 | 0.0960 |
| uc_epi | TranscriptFormer | 0.7454 | 0.7418 | 0.7300 | 0.7653 |
| uc_epi | UCE | 0.7330 | 0.7299 | 0.7123 | 0.7620 |
| uc_epi | scFoundation | 0.7578 | 0.7556 | 0.7418 | 0.7875 |
| uc_epi | scGPT | 0.7320 | 0.7226 | 0.7118 | 0.7479 |
| uc_epi | scLong | 0.4521 | 0.3699 | 0.3776 | 0.3921 |
| uc_epi | scPRINT2 | 0.7156 | 0.7154 | 0.7058 | 0.7398 |
| uc_epi | PCA | 0.7521 | 0.7575 | 0.7456 | 0.7854 |
| uc_epi | UMAP | 0.6125 | 0.5813 | 0.5641 | 0.6875 |
| uc_epi | scVI | 0.7628 | 0.7642 | 0.7555 | 0.7837 |

*Continued on next page*

Table S13: **Human diseases, annotation metrics.** *(continued)*

| Dataset | Method | ACC ↑ | Macro-F1 ↑ | Balanced ACC ↑ | Macro-precision ↑ |
|---|---|---|---|---|---|
| covid_lung | CellFM | 0.5534 | 0.4824 | 0.4967 | 0.5045 |
| covid_lung | CellPLM | 0.7240 | 0.6745 | 0.6948 | 0.6893 |
| covid_lung | GeneCompass | 0.7294 | 0.7003 | 0.7121 | 0.7045 |
| covid_lung | Geneformer | 0.7267 | 0.6846 | 0.6917 | 0.6901 |
| covid_lung | LangCell | 0.7547 | 0.7198 | 0.7354 | 0.7210 |
| covid_lung | Nicheformer | 0.7757 | 0.7410 | 0.7512 | 0.7529 |
| covid_lung | Tabula | 0.1035 | 0.0406 | 0.0662 | 0.0411 |
| covid_lung | TranscriptFormer | 0.8003 | 0.7668 | 0.7755 | 0.7752 |
| covid_lung | UCE | 0.7907 | 0.7590 | 0.7718 | 0.7642 |
| covid_lung | scFoundation | 0.7943 | 0.7630 | 0.7751 | 0.7682 |
| covid_lung | scGPT | 0.7764 | 0.7430 | 0.7601 | 0.7451 |
| covid_lung | scLong | 0.3806 | 0.2656 | 0.3032 | 0.3379 |
| covid_lung | scPRINT2 | 0.7234 | 0.6807 | 0.6968 | 0.6808 |
| covid_lung | PCA | 0.7819 | 0.7469 | 0.7599 | 0.7483 |
| covid_lung | UMAP | 0.6646 | 0.6115 | 0.6055 | 0.6667 |
| covid_lung | scVI | 0.7934 | 0.7558 | 0.7736 | 0.7605 |
| tnbc | CellFM | 0.9749 | 0.9601 | 0.9597 | 0.9608 |
| tnbc | CellPLM | 0.9964 | 0.9965 | 0.9964 | 0.9966 |
| tnbc | GeneCompass | 0.9972 | 0.9968 | 0.9972 | 0.9965 |
| tnbc | Geneformer | 0.9966 | 0.9965 | 0.9967 | 0.9964 |
| tnbc | LangCell | 0.9975 | 0.9977 | 0.9977 | 0.9977 |
| tnbc | Nicheformer | 0.9981 | 0.9983 | 0.9984 | 0.9981 |
| tnbc | Tabula | 0.3204 | 0.2054 | 0.2258 | 0.2139 |
| tnbc | TranscriptFormer | 0.9984 | 0.9987 | 0.9986 | 0.9987 |
| tnbc | UCE | 0.9981 | 0.9983 | 0.9984 | 0.9981 |
| tnbc | scFoundation | 0.9984 | 0.9988 | 0.9988 | 0.9988 |
| tnbc | scGPT | 0.9984 | 0.9986 | 0.9986 | 0.9985 |
| tnbc | scLong | 0.9329 | 0.8918 | 0.8924 | 0.8962 |
| tnbc | scPRINT2 | 0.9948 | 0.9948 | 0.9951 | 0.9945 |
| tnbc | PCA | 0.9976 | 0.9976 | 0.9974 | 0.9978 |
| tnbc | UMAP | 0.9750 | 0.9771 | 0.9734 | 0.9812 |
| tnbc | scVI | 0.9974 | 0.9974 | 0.9975 | 0.9972 |

Table S14: **Human diseases, batch-assessment metrics.**

| Dataset | Method | Scaled iLISI ↑ | kBET ↑ | Scaled cLISI ↑ | Graph conn. ↑ |
|---|---|---|---|---|---|
| ms | CellFM | 0.3218 | 0.5204 | 0.8974 | 0.5808 |
| ms | CellPLM | 0.2412 | 0.4327 | 1.0000 | 0.8210 |
| ms | GeneCompass | 0.2114 | 0.3812 | 0.9996 | 0.9728 |
| ms | Geneformer | 0.2106 | 0.3747 | 0.9984 | 0.9672 |
| ms | LangCell | 0.1974 | 0.3461 | 0.9991 | 0.9758 |
| ms | Nicheformer | 0.1864 | 0.3533 | 1.0000 | 0.9793 |
| ms | Tabula | 0.3867 | 0.7636 | 0.8461 | 0.2897 |
| ms | TranscriptFormer | 0.1327 | 0.3050 | 1.0000 | 0.9831 |
| ms | UCE | 0.1962 | 0.3826 | 1.0000 | 0.9847 |
| ms | scFoundation | 0.1462 | 0.3094 | 1.0000 | 0.9792 |
| ms | scGPT | 0.1645 | 0.3304 | 0.9999 | 0.9851 |
| ms | scLong | 0.3312 | 0.6124 | 0.8820 | 0.3540 |
| ms | scPRINT2 | 0.1974 | 0.3440 | 0.9991 | 0.9668 |
| ms | PCA | 0.2336 | 0.3091 | 0.9979 | 0.9746 |
| ms | UMAP | 0.2039 | 0.4200 | 0.9934 | 0.8485 |
| ms | scVI | 0.1537 | 0.2935 | 0.9988 | 0.9901 |
| uc_epi | CellFM | 0.0806 | 0.7457 | 0.7939 | 0.6287 |
| uc_epi | CellPLM | 0.0652 | 0.6645 | 0.9023 | 0.7810 |
| uc_epi | GeneCompass | 0.0563 | 0.6047 | 0.8682 | 0.9152 |
| uc_epi | Geneformer | 0.0547 | 0.5715 | 0.8689 | 0.9233 |
| uc_epi | LangCell | 0.0398 | 0.4380 | 0.8912 | 0.9619 |
| uc_epi | Nicheformer | 0.0526 | 0.5542 | 0.9085 | 0.9576 |
| uc_epi | Tabula | 0.0851 | 0.7937 | 0.6512 | 0.5479 |
| uc_epi | TranscriptFormer | 0.0516 | 0.5467 | 0.9295 | 0.9657 |
| uc_epi | UCE | 0.0493 | 0.5531 | 0.9224 | 0.9659 |
| uc_epi | scFoundation | 0.0399 | 0.4198 | 0.9250 | 0.9645 |
| uc_epi | scGPT | 0.0522 | 0.5462 | 0.9202 | 0.9485 |
| uc_epi | scLong | 0.0782 | 0.7192 | 0.7759 | 0.5075 |
| uc_epi | scPRINT2 | 0.0456 | 0.4963 | 0.8945 | 0.9489 |
| uc_epi | PCA | 0.0458 | 0.4844 | 0.9086 | 0.9678 |
| uc_epi | UMAP | 0.0558 | 0.5363 | 0.9209 | 0.8315 |
| uc_epi | scVI | 0.0325 | 0.3843 | 0.9141 | 0.9830 |
| covid_lung | CellFM | 0.3212 | 0.8304 | 0.7727 | 0.1551 |
| covid_lung | CellPLM | 0.2129 | 0.4239 | 0.9733 | 0.6711 |
| covid_lung | GeneCompass | 0.2198 | 0.4688 | 0.9494 | 0.9516 |
| covid_lung | Geneformer | 0.2488 | 0.6097 | 0.9424 | 0.9619 |
| covid_lung | LangCell | 0.2151 | 0.4303 | 0.9616 | 0.9620 |
| covid_lung | Nicheformer | 0.2006 | 0.3783 | 0.9707 | 0.9629 |
| covid_lung | Tabula | 0.3226 | 0.8865 | 0.7274 | 0.0909 |
| covid_lung | TranscriptFormer | 0.1663 | 0.2652 | 0.9823 | 0.9451 |
| covid_lung | UCE | 0.1655 | 0.2622 | 0.9827 | 0.9330 |
| covid_lung | scFoundation | 0.1775 | 0.2861 | 0.9798 | 0.9484 |
| covid_lung | scGPT | 0.1851 | 0.3249 | 0.9745 | 0.9639 |
| covid_lung | scLong | 0.3230 | 0.8148 | 0.7354 | 0.0401 |
| covid_lung | scPRINT2 | 0.2098 | 0.4445 | 0.9492 | 0.9477 |
| covid_lung | PCA | 0.2096 | 0.3942 | 0.9691 | 0.9510 |
| covid_lung | UMAP | 0.2166 | 0.4755 | 0.9735 | 0.8108 |
| covid_lung | scVI | 0.1661 | 0.2674 | 0.9781 | 0.9737 |

*Continued on next page*

Table S14: **Human diseases, batch-assessment metrics.** *(continued)*

| Dataset | Method | Scaled iLISI ↑ | kBET ↑ | Scaled cLISI ↑ | Graph conn. ↑ |
|---|---|---|---|---|---|
| tnbc | CellFM | 0.1062 | 0.6249 | 0.8304 | 0.8779 |
| tnbc | CellPLM | 0.0818 | 0.2790 | 1.0000 | 0.8521 |
| tnbc | GeneCompass | 0.0813 | 0.2655 | 1.0000 | 0.9994 |
| tnbc | Geneformer | 0.0770 | 0.2083 | 1.0000 | 0.9995 |
| tnbc | LangCell | 0.0518 | 0.0559 | 1.0000 | 0.9996 |
| tnbc | Nicheformer | 0.0673 | 0.1365 | 1.0000 | 0.9996 |
| tnbc | Tabula | 0.1155 | 0.9178 | 0.4764 | 0.7092 |
| tnbc | TranscriptFormer | 0.0650 | 0.1252 | 1.0000 | 0.9996 |
| tnbc | UCE | 0.0749 | 0.1937 | 1.0000 | 0.9979 |
| tnbc | scFoundation | 0.0545 | 0.0607 | 1.0000 | 0.9957 |
| tnbc | scGPT | 0.0710 | 0.1648 | 1.0000 | 0.9982 |
| tnbc | scLong | 0.1105 | 0.6278 | 0.7660 | 0.6824 |
| tnbc | scPRINT2 | 0.0670 | 0.1478 | 1.0000 | 0.9995 |
| tnbc | PCA | 0.0689 | 0.1405 | 1.0000 | 0.9994 |
| tnbc | UMAP | 0.0767 | 0.2863 | 1.0000 | 0.9395 |
| tnbc | scVI | 0.0668 | 0.1406 | 1.0000 | 0.9984 |

Table S15: **Human diseases, reconstruction metrics ($\log(1 + \mathrm{CP10k})$).**

| Dataset | Method | Pearson $r$ ↑ | Spearman ↑ | MAE ↓ | NZ top-50 ↑ |
|---|---|---|---|---|---|
| ms | CellFM | 0.5190 | 0.3573 | 0.0889 | 0.4113 |
| ms | CellPLM | 0.5788 | 0.3997 | 0.0794 | 0.4501 |
| ms | GeneCompass | 0.5980 | 0.4519 | 0.0733 | 0.4620 |
| ms | Geneformer | 0.6133 | 0.4551 | 0.0717 | 0.4732 |
| ms | LangCell | 0.6131 | 0.4613 | 0.0702 | 0.4715 |
| ms | Nicheformer | 0.6244 | 0.4732 | 0.0696 | 0.4775 |
| ms | Tabula | 0.4177 | 0.2522 | 0.1127 | 0.3691 |
| ms | TranscriptFormer | 0.6203 | 0.4668 | 0.0695 | 0.4737 |
| ms | UCE | 0.6191 | 0.4722 | 0.0705 | 0.4789 |
| ms | scFoundation | 0.6204 | 0.4622 | 0.0709 | 0.4830 |
| ms | scGPT | 0.6247 | 0.4652 | 0.0697 | 0.4781 |
| ms | scLong | 0.4288 | 0.2857 | 0.1044 | 0.3669 |
| ms | scPRINT2 | 0.6206 | 0.4576 | 0.0688 | 0.4674 |
| ms | PCA | 0.8057 | 0.7114 | 0.0467 | 0.6686 |
| ms | UMAP | 0.5762 | 0.4210 | 0.0761 | 0.4364 |
| ms | scVI | 0.6344 | 0.4829 | 0.0677 | 0.4863 |
| ms | *Train mean* | 0.4183 | 0.2197 | 0.1467 | 0.3711 |
| ms | *Train median* | 0.0876 | 0.0754 | 0.0787 | 0.0229 |
| ms | *All zero* | 0.0000 | 0.0000 | 0.0799 | 0.0000 |
| uc_epi | CellFM | 0.7281 | 0.5061 | 0.1021 | 0.5818 |
| uc_epi | CellPLM | 0.7743 | 0.5353 | 0.0953 | 0.6182 |
| uc_epi | GeneCompass | 0.7741 | 0.5490 | 0.0912 | 0.6195 |
| uc_epi | Geneformer | 0.8088 | 0.5643 | 0.0828 | 0.6398 |
| uc_epi | LangCell | 0.8221 | 0.5717 | 0.0801 | 0.6477 |
| uc_epi | Nicheformer | 0.8125 | 0.5725 | 0.0816 | 0.6459 |
| uc_epi | Tabula | 0.6304 | 0.4611 | 0.1286 | 0.5080 |
| uc_epi | TranscriptFormer | 0.8287 | 0.5789 | 0.0779 | 0.6630 |
| uc_epi | UCE | 0.8089 | 0.5784 | 0.0832 | 0.6499 |
| uc_epi | scFoundation | 0.8398 | 0.5842 | 0.0756 | 0.6672 |
| uc_epi | scGPT | 0.8083 | 0.5704 | 0.0828 | 0.6432 |
| uc_epi | scLong | 0.6782 | 0.4769 | 0.1154 | 0.5458 |
| uc_epi | scPRINT2 | 0.8283 | 0.5672 | 0.0778 | 0.6456 |
| uc_epi | PCA | 0.9402 | 0.7899 | 0.0380 | 0.8346 |
| uc_epi | UMAP | 0.7648 | 0.5347 | 0.0931 | 0.6116 |
| uc_epi | scVI | 0.8320 | 0.5884 | 0.0771 | 0.6619 |
| uc_epi | *Train mean* | 0.6308 | 0.3521 | 0.1532 | 0.5021 |
| uc_epi | *Train median* | 0.5516 | 0.4266 | 0.1081 | 0.2663 |
| uc_epi | *All zero* | 0.0000 | 0.0000 | 0.1283 | 0.0000 |

*Continued on next page*

Table S15: **Human diseases, reconstruction metrics ($\log(1 + \mathrm{CP10k})$).** *(continued)*

| Dataset | Method | Pearson $r$ ↑ | Spearman ↑ | MAE ↓ | NZ top-50 ↑ |
|---|---|---|---|---|---|
| covid_lung | CellFM | 0.4458 | 0.3402 | 0.1164 | 0.3692 |
| covid_lung | CellPLM | 0.5411 | 0.4127 | 0.1037 | 0.4548 |
| covid_lung | GeneCompass | 0.5581 | 0.4349 | 0.0947 | 0.4721 |
| covid_lung | Geneformer | 0.5771 | 0.4492 | 0.0895 | 0.4918 |
| covid_lung | LangCell | 0.5886 | 0.4512 | 0.0883 | 0.4959 |
| covid_lung | Nicheformer | 0.5787 | 0.4516 | 0.0894 | 0.4907 |
| covid_lung | Tabula | 0.1988 | 0.1660 | 0.1673 | 0.1579 |
| covid_lung | TranscriptFormer | 0.6147 | 0.4732 | 0.0836 | 0.5243 |
| covid_lung | UCE | 0.5850 | 0.4575 | 0.0907 | 0.4970 |
| covid_lung | scFoundation | 0.6110 | 0.4669 | 0.0867 | 0.5171 |
| covid_lung | scGPT | 0.5763 | 0.4482 | 0.0913 | 0.4869 |
| covid_lung | scLong | 0.3061 | 0.2330 | 0.1437 | 0.2426 |
| covid_lung | scPRINT2 | 0.6029 | 0.4504 | 0.0845 | 0.4989 |
| covid_lung | PCA | 0.7716 | 0.6170 | 0.0593 | 0.6879 |
| covid_lung | UMAP | 0.5312 | 0.3984 | 0.0974 | 0.4450 |
| covid_lung | scVI | 0.6202 | 0.4819 | 0.0846 | 0.5285 |
| covid_lung | *Train mean* | 0.1937 | 0.1624 | 0.1760 | 0.1502 |
| covid_lung | *Train median* | 0.0000 | 0.0000 | 0.0937 | 0.0000 |
| covid_lung | *All zero* | 0.0000 | 0.0000 | 0.0937 | 0.0000 |
| tnbc | CellFM | 0.6266 | 0.4682 | 0.0699 | 0.5019 |
| tnbc | CellPLM | 0.6624 | 0.5101 | 0.0641 | 0.5314 |
| tnbc | GeneCompass | 0.6795 | 0.5257 | 0.0606 | 0.5415 |
| tnbc | Geneformer | 0.6909 | 0.5375 | 0.0577 | 0.5414 |
| tnbc | LangCell | 0.7102 | 0.5440 | 0.0549 | 0.5387 |
| tnbc | Nicheformer | 0.6950 | 0.5365 | 0.0562 | 0.5339 |
| tnbc | Tabula | 0.3743 | 0.2676 | 0.1165 | 0.2869 |
| tnbc | TranscriptFormer | 0.7153 | 0.5555 | 0.0546 | 0.5510 |
| tnbc | UCE | 0.7047 | 0.5507 | 0.0570 | 0.5527 |
| tnbc | scFoundation | 0.7446 | 0.5646 | 0.0515 | 0.5682 |
| tnbc | scGPT | 0.7044 | 0.5433 | 0.0572 | 0.5522 |
| tnbc | scLong | 0.5624 | 0.4195 | 0.0785 | 0.4509 |
| tnbc | scPRINT2 | 0.6931 | 0.5284 | 0.0561 | 0.5119 |
| tnbc | PCA | 0.8485 | 0.6903 | 0.0381 | 0.6623 |
| tnbc | UMAP | 0.6776 | 0.5034 | 0.0620 | 0.5265 |
| tnbc | scVI | 0.7450 | 0.5666 | 0.0521 | 0.5680 |
| tnbc | *Train mean* | 0.3632 | 0.2413 | 0.1287 | 0.2830 |
| tnbc | *Train median* | 0.2750 | 0.2276 | 0.0757 | 0.1052 |
| tnbc | *All zero* | 0.0000 | 0.0000 | 0.0786 | 0.0000 |

Table S16: **Mouse organs, clustering metrics.**

| Dataset | Method | uARI ↑ | NMI ↑ | ASW ↑ | ACC ↑ |
|---|---|---|---|---|---|
| mouse_marrow | CellFM | 0.0831 | 0.2982 | 0.4019 | 0.1628 |
| mouse_marrow | CellPLM | 0.4031 | 0.6404 | 0.4749 | 0.5150 |
| mouse_marrow | GeneCompass | 0.1528 | 0.3142 | 0.4227 | 0.2728 |
| mouse_marrow | Geneformer | 0.3329 | 0.5776 | 0.4855 | 0.4753 |
| mouse_marrow | LangCell | 0.4593 | 0.6949 | 0.5148 | 0.6104 |
| mouse_marrow | Nicheformer | 0.4415 | 0.6604 | 0.5212 | 0.5680 |
| mouse_marrow | Tabula | 0.0324 | 0.0667 | 0.4722 | 0.1537 |
| mouse_marrow | TranscriptFormer | 0.4394 | 0.6834 | 0.5129 | 0.5684 |
| mouse_marrow | UCE | 0.4865 | 0.7075 | 0.5328 | 0.6311 |
| mouse_marrow | scFoundation | 0.4785 | 0.7019 | 0.5207 | 0.6260 |
| mouse_marrow | scGPT | 0.4644 | 0.6852 | 0.5193 | 0.6105 |
| mouse_marrow | scLong | 0.0956 | 0.3036 | 0.3958 | 0.1775 |
| mouse_marrow | scPRINT2 | 0.3223 | 0.5914 | 0.5121 | 0.4627 |
| mouse_marrow | PCA | 0.5390 | 0.7332 | 0.5131 | 0.6923 |
| mouse_marrow | UMAP | 0.4114 | 0.6803 | 0.5649 | 0.5258 |
| mouse_marrow | scVI | 0.4357 | 0.6907 | 0.5492 | 0.5866 |
| mouse_lung | CellFM | 0.0475 | 0.2141 | 0.3583 | 0.1590 |
| mouse_lung | CellPLM | 0.3667 | 0.6801 | 0.5125 | 0.5219 |
| mouse_lung | GeneCompass | 0.1515 | 0.3125 | 0.3875 | 0.2799 |
| mouse_lung | Geneformer | 0.3742 | 0.6858 | 0.4901 | 0.5555 |
| mouse_lung | LangCell | 0.4211 | 0.7339 | 0.5306 | 0.5841 |
| mouse_lung | Nicheformer | 0.3768 | 0.7227 | 0.5318 | 0.5423 |
| mouse_lung | Tabula | 0.0039 | 0.0185 | 0.4594 | 0.1156 |
| mouse_lung | TranscriptFormer | 0.4466 | 0.7628 | 0.5501 | 0.6217 |
| mouse_lung | UCE | 0.4794 | 0.7794 | 0.5831 | 0.6616 |
| mouse_lung | scFoundation | 0.4418 | 0.7656 | 0.5528 | 0.6224 |
| mouse_lung | scGPT | 0.4627 | 0.7630 | 0.5546 | 0.6272 |
| mouse_lung | scLong | 0.0643 | 0.2568 | 0.3212 | 0.1563 |
| mouse_lung | scPRINT2 | 0.3809 | 0.6922 | 0.5346 | 0.5617 |
| mouse_lung | PCA | 0.4987 | 0.7990 | 0.5526 | 0.6691 |
| mouse_lung | UMAP | 0.3439 | 0.7343 | 0.6550 | 0.5748 |
| mouse_lung | scVI | 0.4765 | 0.7920 | 0.5692 | 0.6587 |

*Continued on next page*

Table S16: **Mouse organs, clustering metrics.** *(continued)*

| Dataset | Method | uARI ↑ | NMI ↑ | ASW ↑ | ACC ↑ |
|---|---|---|---|---|---|
| mouse_kidney | CellFM | 0.0831 | 0.2594 | 0.3549 | 0.1691 |
| mouse_kidney | CellPLM | 0.4588 | 0.6750 | 0.5580 | 0.5447 |
| mouse_kidney | GeneCompass | 0.1575 | 0.3027 | 0.3925 | 0.2786 |
| mouse_kidney | Geneformer | 0.4180 | 0.6611 | 0.4895 | 0.5081 |
| mouse_kidney | LangCell | 0.5241 | 0.7399 | 0.5290 | 0.6086 |
| mouse_kidney | Nicheformer | 0.4332 | 0.6738 | 0.5168 | 0.5253 |
| mouse_kidney | Tabula | 0.0090 | 0.0232 | 0.4216 | 0.1274 |
| mouse_kidney | TranscriptFormer | 0.5363 | 0.7322 | 0.5267 | 0.6326 |
| mouse_kidney | UCE | 0.4918 | 0.7097 | 0.5512 | 0.5632 |
| mouse_kidney | scFoundation | 0.4814 | 0.6893 | 0.5271 | 0.5586 |
| mouse_kidney | scGPT | 0.5052 | 0.7112 | 0.5232 | 0.5845 |
| mouse_kidney | scLong | 0.0785 | 0.2579 | 0.3317 | 0.1526 |
| mouse_kidney | scPRINT2 | 0.4369 | 0.7076 | 0.5200 | 0.4975 |
| mouse_kidney | PCA | 0.6583 | 0.8014 | 0.5440 | 0.7061 |
| mouse_kidney | UMAP | 0.4257 | 0.7223 | 0.5855 | 0.4970 |
| mouse_kidney | scVI | 0.4802 | 0.7461 | 0.5383 | 0.5401 |
| mouse_heart | CellFM | 0.0367 | 0.1648 | 0.4241 | 0.1369 |
| mouse_heart | CellPLM | 0.4942 | 0.7014 | 0.6201 | 0.5564 |
| mouse_heart | GeneCompass | 0.0451 | 0.1621 | 0.2421 | 0.1802 |
| mouse_heart | Geneformer | 0.2747 | 0.5545 | 0.3373 | 0.3899 |
| mouse_heart | LangCell | 0.4010 | 0.6908 | 0.5408 | 0.5019 |
| mouse_heart | Nicheformer | 0.3557 | 0.6891 | 0.5753 | 0.4810 |
| mouse_heart | Tabula | 0.0082 | 0.0298 | 0.4756 | 0.1346 |
| mouse_heart | TranscriptFormer | 0.4045 | 0.6821 | 0.4544 | 0.5113 |
| mouse_heart | UCE | 0.5647 | 0.7551 | 0.6487 | 0.6149 |
| mouse_heart | scFoundation | 0.4777 | 0.7390 | 0.5874 | 0.5430 |
| mouse_heart | scGPT | 0.4272 | 0.7194 | 0.5622 | 0.4949 |
| mouse_heart | scLong | 0.0407 | 0.1805 | 0.2784 | 0.1459 |
| mouse_heart | scPRINT2 | 0.3486 | 0.6655 | 0.5102 | 0.4632 |
| mouse_heart | PCA | 0.5652 | 0.7639 | 0.5657 | 0.6140 |
| mouse_heart | UMAP | 0.2741 | 0.6243 | 0.6858 | 0.3739 |
| mouse_heart | scVI | 0.4910 | 0.7192 | 0.5533 | 0.5622 |

Table S17: **Mouse organs, annotation metrics.**

| Dataset | Method | ACC ↑ | Macro-F1 ↑ | Balanced ACC ↑ | Macro-precision ↑ |
|---|---|---|---|---|---|
| mouse_marrow | CellFM | 0.8338 | 0.7120 | 0.7125 | 0.7326 |
| mouse_marrow | CellPLM | 0.8811 | 0.7974 | 0.7924 | 0.8120 |
| mouse_marrow | GeneCompass | 0.7856 | 0.6379 | 0.6317 | 0.6863 |
| mouse_marrow | Geneformer | 0.8885 | 0.8118 | 0.8143 | 0.8247 |
| mouse_marrow | LangCell | 0.9021 | 0.8340 | 0.8427 | 0.8378 |
| mouse_marrow | Nicheformer | 0.9024 | 0.8371 | 0.8422 | 0.8393 |
| mouse_marrow | Tabula | 0.2501 | 0.0593 | 0.0920 | 0.0667 |
| mouse_marrow | TranscriptFormer | 0.9033 | 0.8353 | 0.8485 | 0.8310 |
| mouse_marrow | UCE | 0.8905 | 0.8178 | 0.8248 | 0.8185 |
| mouse_marrow | scFoundation | 0.9177 | 0.8608 | 0.8774 | 0.8535 |
| mouse_marrow | scGPT | 0.8940 | 0.8200 | 0.8325 | 0.8158 |
| mouse_marrow | scLong | 0.7402 | 0.5436 | 0.5566 | 0.5967 |
| mouse_marrow | scPRINT2 | 0.8924 | 0.8186 | 0.8251 | 0.8244 |
| mouse_marrow | PCA | 0.9103 | 0.8415 | 0.8494 | 0.8385 |
| mouse_marrow | UMAP | 0.8660 | 0.7495 | 0.7435 | 0.7932 |
| mouse_marrow | scVI | 0.9225 | 0.8711 | 0.8830 | 0.8646 |
| mouse_lung | CellFM | 0.8104 | 0.5831 | 0.5896 | 0.6101 |
| mouse_lung | CellPLM | 0.8839 | 0.7583 | 0.7653 | 0.7894 |
| mouse_lung | GeneCompass | 0.7742 | 0.5399 | 0.5544 | 0.5643 |
| mouse_lung | Geneformer | 0.8966 | 0.7443 | 0.7640 | 0.7413 |
| mouse_lung | LangCell | 0.9206 | 0.8418 | 0.8383 | 0.8659 |
| mouse_lung | Nicheformer | 0.9315 | 0.8686 | 0.8635 | 0.8907 |
| mouse_lung | Tabula | 0.0979 | 0.0156 | 0.0390 | 0.0204 |
| mouse_lung | TranscriptFormer | 0.9323 | 0.8686 | 0.8659 | 0.8855 |
| mouse_lung | UCE | 0.9160 | 0.8553 | 0.8549 | 0.8755 |
| mouse_lung | scFoundation | 0.9418 | 0.8907 | 0.8924 | 0.9000 |
| mouse_lung | scGPT | 0.9153 | 0.8589 | 0.8536 | 0.8800 |
| mouse_lung | scLong | 0.7308 | 0.4582 | 0.4666 | 0.5140 |
| mouse_lung | scPRINT2 | 0.9246 | 0.8394 | 0.8480 | 0.8558 |
| mouse_lung | PCA | 0.9415 | 0.8725 | 0.8679 | 0.8888 |
| mouse_lung | UMAP | 0.9028 | 0.7739 | 0.7620 | 0.8359 |
| mouse_lung | scVI | 0.9491 | 0.8807 | 0.8813 | 0.8947 |

*Continued on next page*

Table S17: **Mouse organs, annotation metrics.** *(continued)*

| Dataset | Method | ACC ↑ | Macro-F1 ↑ | Balanced ACC ↑ | Macro-precision ↑ |
|---|---|---|---|---|---|
| mouse_kidney | CellFM | 0.8277 | 0.6571 | 0.6624 | 0.6712 |
| mouse_kidney | CellPLM | 0.8043 | 0.6795 | 0.6785 | 0.6984 |
| mouse_kidney | GeneCompass | 0.7893 | 0.5995 | 0.6000 | 0.6478 |
| mouse_kidney | Geneformer | 0.8890 | 0.8045 | 0.8052 | 0.8244 |
| mouse_kidney | LangCell | 0.9342 | 0.8453 | 0.8359 | 0.8697 |
| mouse_kidney | Nicheformer | 0.9163 | 0.8474 | 0.8528 | 0.8535 |
| mouse_kidney | Tabula | 0.2348 | 0.0386 | 0.0684 | 0.0443 |
| mouse_kidney | TranscriptFormer | 0.9242 | 0.8497 | 0.8462 | 0.8652 |
| mouse_kidney | UCE | 0.8913 | 0.8221 | 0.8209 | 0.8392 |
| mouse_kidney | scFoundation | 0.9491 | 0.8719 | 0.8815 | 0.8737 |
| mouse_kidney | scGPT | 0.8832 | 0.8210 | 0.8233 | 0.8281 |
| mouse_kidney | scLong | 0.7334 | 0.5502 | 0.5528 | 0.5835 |
| mouse_kidney | scPRINT2 | 0.9199 | 0.8329 | 0.8362 | 0.8424 |
| mouse_kidney | PCA | 0.9167 | 0.8340 | 0.8353 | 0.8452 |
| mouse_kidney | UMAP | 0.9167 | 0.7918 | 0.7934 | 0.8081 |
| mouse_kidney | scVI | 0.9244 | 0.8436 | 0.8433 | 0.8603 |
| mouse_heart | CellFM | 0.8986 | 0.7460 | 0.7416 | 0.7731 |
| mouse_heart | CellPLM | 0.9376 | 0.8517 | 0.8701 | 0.8523 |
| mouse_heart | GeneCompass | 0.6694 | 0.4359 | 0.4272 | 0.4992 |
| mouse_heart | Geneformer | 0.9378 | 0.8600 | 0.8837 | 0.8637 |
| mouse_heart | LangCell | 0.9591 | 0.9080 | 0.9139 | 0.9205 |
| mouse_heart | Nicheformer | 0.9605 | 0.9093 | 0.9188 | 0.9211 |
| mouse_heart | Tabula | 0.4056 | 0.0972 | 0.1186 | 0.0865 |
| mouse_heart | TranscriptFormer | 0.9536 | 0.9039 | 0.9116 | 0.9179 |
| mouse_heart | UCE | 0.9575 | 0.8986 | 0.9029 | 0.9125 |
| mouse_heart | scFoundation | 0.9685 | 0.9211 | 0.9262 | 0.9336 |
| mouse_heart | scGPT | 0.9586 | 0.9068 | 0.9133 | 0.9121 |
| mouse_heart | scLong | 0.7858 | 0.5045 | 0.4964 | 0.6112 |
| mouse_heart | scPRINT2 | 0.9564 | 0.9128 | 0.9163 | 0.9269 |
| mouse_heart | PCA | 0.9584 | 0.8870 | 0.8927 | 0.8963 |
| mouse_heart | UMAP | 0.8896 | 0.7162 | 0.7273 | 0.7908 |
| mouse_heart | scVI | 0.9541 | 0.8989 | 0.9221 | 0.8976 |

Table S18: **Mouse organs, batch-assessment metrics.**

| Dataset | Method | Scaled iLISI ↑ | kBET ↑ | Scaled cLISI ↑ | Graph conn. ↑ |
|---|---|---|---|---|---|
| mouse_marrow | CellFM | 0.3723 | 0.6523 | 0.9214 | 0.5835 |
| mouse_marrow | CellPLM | 0.2984 | 0.4776 | 0.9916 | 0.7198 |
| mouse_marrow | GeneCompass | 0.3607 | 0.6053 | 0.9062 | 0.7071 |
| mouse_marrow | Geneformer | 0.3050 | 0.4867 | 0.9748 | 0.8868 |
| mouse_marrow | LangCell | 0.2856 | 0.4032 | 0.9862 | 0.9483 |
| mouse_marrow | Nicheformer | 0.2858 | 0.4064 | 0.9943 | 0.9305 |
| mouse_marrow | Tabula | 0.4295 | 0.8893 | 0.7080 | 0.3400 |
| mouse_marrow | TranscriptFormer | 0.2921 | 0.4581 | 0.9913 | 0.9568 |
| mouse_marrow | UCE | 0.3031 | 0.5305 | 0.9945 | 0.9581 |
| mouse_marrow | scFoundation | 0.2612 | 0.3671 | 0.9940 | 0.9576 |
| mouse_marrow | scGPT | 0.2888 | 0.4559 | 0.9893 | 0.9288 |
| mouse_marrow | scLong | 0.3470 | 0.5408 | 0.9174 | 0.5525 |
| mouse_marrow | scPRINT2 | 0.2469 | 0.3765 | 0.9869 | 0.9309 |
| mouse_marrow | PCA | 0.2789 | 0.3468 | 0.9943 | 0.9639 |
| mouse_marrow | UMAP | 0.3242 | 0.5707 | 0.9958 | 0.8075 |
| mouse_marrow | scVI | 0.2042 | 0.2972 | 0.9945 | 0.9746 |
| mouse_lung | CellFM | 0.2261 | 0.8662 | 0.9375 | 0.4350 |
| mouse_lung | CellPLM | 0.1558 | 0.7655 | 0.9952 | 0.8424 |
| mouse_lung | GeneCompass | 0.2455 | 0.8354 | 0.9502 | 0.5838 |
| mouse_lung | Geneformer | 0.1678 | 0.7903 | 0.9899 | 0.9166 |
| mouse_lung | LangCell | 0.1261 | 0.6766 | 0.9944 | 0.9664 |
| mouse_lung | Nicheformer | 0.1448 | 0.7497 | 0.9965 | 0.9657 |
| mouse_lung | Tabula | 0.3094 | 0.9077 | 0.8510 | 0.1608 |
| mouse_lung | TranscriptFormer | 0.1632 | 0.7843 | 0.9968 | 0.9599 |
| mouse_lung | UCE | 0.1377 | 0.7203 | 0.9980 | 0.9625 |
| mouse_lung | scFoundation | 0.1152 | 0.6854 | 0.9971 | 0.9672 |
| mouse_lung | scGPT | 0.1591 | 0.7763 | 0.9961 | 0.9483 |
| mouse_lung | scLong | 0.2502 | 0.8381 | 0.9381 | 0.3855 |
| mouse_lung | scPRINT2 | 0.1208 | 0.7038 | 0.9964 | 0.9630 |
| mouse_lung | PCA | 0.1265 | 0.7048 | 0.9981 | 0.9897 |
| mouse_lung | UMAP | 0.1051 | 0.7205 | 0.9992 | 0.8907 |
| mouse_lung | scVI | 0.0901 | 0.6527 | 0.9986 | 0.9746 |
| mouse_heart | CellFM | 0.1557 | 0.6608 | 0.9183 | 0.6115 |
| mouse_heart | CellPLM | 0.1306 | 0.5792 | 0.9995 | 0.8629 |
| mouse_heart | GeneCompass | 0.1126 | 0.5497 | 0.9049 | 0.5202 |
| mouse_heart | Geneformer | 0.1211 | 0.5734 | 0.9942 | 0.9026 |
| mouse_heart | LangCell | 0.1226 | 0.5429 | 0.9995 | 0.9221 |
| mouse_heart | Nicheformer | 0.1059 | 0.4988 | 0.9997 | 0.9548 |
| mouse_heart | Tabula | 0.2406 | 0.8848 | 0.8391 | 0.2651 |
| mouse_heart | TranscriptFormer | 0.1212 | 0.5301 | 0.9995 | 0.9340 |
| mouse_heart | UCE | 0.1183 | 0.5574 | 0.9999 | 0.9522 |
| mouse_heart | scFoundation | 0.0982 | 0.4790 | 0.9998 | 0.9379 |
| mouse_heart | scGPT | 0.1020 | 0.4841 | 0.9996 | 0.9345 |
| mouse_heart | scLong | 0.1485 | 0.6239 | 0.9148 | 0.5552 |
| mouse_heart | scPRINT2 | 0.0851 | 0.4975 | 0.9990 | 0.9145 |
| mouse_heart | PCA | 0.1515 | 0.5276 | 0.9995 | 0.9491 |
| mouse_heart | UMAP | 0.1492 | 0.6222 | 0.9994 | 0.8182 |
| mouse_heart | scVI | 0.1184 | 0.5113 | 0.9997 | 0.9609 |

*Continued on next page*

Table S18: **Mouse organs, batch-assessment metrics.** *(continued)*

| Dataset | Method | Scaled iLISI ↑ | kBET ↑ | Scaled cLISI ↑ | Graph conn. ↑ |
|---|---|---|---|---|---|
| mouse_kidney | CellFM | 0.3491 | 0.8395 | 0.9138 | 0.4381 |
| mouse_kidney | CellPLM | 0.2643 | 0.7033 | 0.9817 | 0.7616 |
| mouse_kidney | GeneCompass | 0.3442 | 0.8202 | 0.9188 | 0.6232 |
| mouse_kidney | Geneformer | 0.3006 | 0.7568 | 0.9754 | 0.8770 |
| mouse_kidney | LangCell | 0.2621 | 0.6863 | 0.9928 | 0.8655 |
| mouse_kidney | Nicheformer | 0.2742 | 0.7073 | 0.9844 | 0.8885 |
| mouse_kidney | Tabula | 0.4160 | 0.9253 | 0.7840 | 0.2380 |
| mouse_kidney | TranscriptFormer | 0.2704 | 0.7120 | 0.9918 | 0.9176 |
| mouse_kidney | UCE | 0.2580 | 0.6546 | 0.9907 | 0.8475 |
| mouse_kidney | scFoundation | 0.2521 | 0.6590 | 0.9880 | 0.9085 |
| mouse_kidney | scGPT | 0.2787 | 0.7186 | 0.9831 | 0.8855 |
| mouse_kidney | scLong | 0.3489 | 0.8004 | 0.9052 | 0.4082 |
| mouse_kidney | scPRINT2 | 0.2216 | 0.6485 | 0.9939 | 0.9098 |
| mouse_kidney | PCA | 0.1723 | 0.5648 | 0.9984 | 0.8428 |
| mouse_kidney | UMAP | 0.1703 | 0.6471 | 0.9979 | 0.7485 |
| mouse_kidney | scVI | 0.1268 | 0.5276 | 0.9977 | 0.8699 |

Table S19: **Mouse organs, reconstruction metrics ($\log(1+\mathrm{CP10k})$).**

| Dataset | Method | Pearson $r$ ↑ | Spearman ↑ | MAE ↓ | NZ top-50 ↑ |
|---|---|---|---|---|---|
| mouse_marrow | CellFM | 0.7368 | 0.5063 | 0.0884 | 0.5214 |
| mouse_marrow | CellPLM | 0.7671 | 0.5255 | 0.0824 | 0.5433 |
| mouse_marrow | GeneCompass | 0.7350 | 0.5060 | 0.0894 | 0.5213 |
| mouse_marrow | Geneformer | 0.7914 | 0.5494 | 0.0765 | 0.5497 |
| mouse_marrow | LangCell | 0.8067 | 0.5594 | 0.0738 | 0.5615 |
| mouse_marrow | Nicheformer | 0.7977 | 0.5561 | 0.0751 | 0.5571 |
| mouse_marrow | Tabula | 0.5277 | 0.3562 | 0.1453 | 0.4100 |
| mouse_marrow | TranscriptFormer | 0.8020 | 0.5633 | 0.0743 | 0.5667 |
| mouse_marrow | UCE | 0.7948 | 0.5521 | 0.0768 | 0.5644 |
| mouse_marrow | scFoundation | 0.8160 | 0.5657 | 0.0722 | 0.5841 |
| mouse_marrow | scGPT | 0.7901 | 0.5468 | 0.0766 | 0.5497 |
| mouse_marrow | scLong | 0.6988 | 0.4774 | 0.0993 | 0.4939 |
| mouse_marrow | scPRINT2 | 0.8068 | 0.5558 | 0.0734 | 0.5554 |
| mouse_marrow | PCA | 0.8573 | 0.6355 | 0.0631 | 0.6358 |
| mouse_marrow | UMAP | 0.7693 | 0.5290 | 0.0807 | 0.5365 |
| mouse_marrow | scVI | 0.8123 | 0.5668 | 0.0722 | 0.5666 |
| mouse_marrow | *Train mean* | 0.5101 | 0.3236 | 0.1697 | 0.3967 |
| mouse_marrow | *Train median* | 0.4129 | 0.2526 | 0.1174 | 0.1113 |
| mouse_marrow | *All zero* | 0.0000 | 0.0000 | 0.1221 | 0.0000 |
| mouse_lung | CellFM | 0.7237 | 0.5364 | 0.1039 | 0.5547 |
| mouse_lung | CellPLM | 0.7449 | 0.5557 | 0.0982 | 0.5661 |
| mouse_lung | GeneCompass | 0.6906 | 0.5129 | 0.1119 | 0.5290 |
| mouse_lung | Geneformer | 0.7919 | 0.5938 | 0.0854 | 0.5924 |
| mouse_lung | LangCell | 0.8069 | 0.6075 | 0.0825 | 0.6008 |
| mouse_lung | Nicheformer | 0.7869 | 0.5944 | 0.0859 | 0.5911 |
| mouse_lung | Tabula | 0.3501 | 0.2784 | 0.1750 | 0.3056 |
| mouse_lung | TranscriptFormer | 0.7945 | 0.6026 | 0.0833 | 0.5931 |
| mouse_lung | UCE | 0.7893 | 0.5942 | 0.0856 | 0.5953 |
| mouse_lung | scFoundation | 0.8117 | 0.6116 | 0.0808 | 0.6085 |
| mouse_lung | scGPT | 0.7878 | 0.5932 | 0.0862 | 0.5928 |
| mouse_lung | scLong | 0.6036 | 0.4519 | 0.1341 | 0.4721 |
| mouse_lung | scPRINT2 | 0.8105 | 0.6040 | 0.0811 | 0.5932 |
| mouse_lung | PCA | 0.8339 | 0.6677 | 0.0736 | 0.6249 |
| mouse_lung | UMAP | 0.7661 | 0.5761 | 0.0928 | 0.5773 |
| mouse_lung | scVI | 0.8134 | 0.6126 | 0.0798 | 0.6063 |
| mouse_lung | *Train mean* | 0.3458 | 0.2719 | 0.1958 | 0.3056 |
| mouse_lung | *Train median* | 0.2372 | 0.1502 | 0.1458 | 0.0700 |
| mouse_lung | *All zero* | 0.0000 | 0.0000 | 0.1441 | 0.0000 |

*Continued on next page*

Table S19: **Mouse organs, reconstruction metrics ($\log(1 + \mathrm{CP10k})$).** *(continued)*

| Dataset | Method | Pearson $r$ ↑ | Spearman ↑ | MAE ↓ | NZ top-50 ↑ |
|---|---|---|---|---|---|
| mouse_heart | CellFM | 0.4617 | 0.4077 | 0.1932 | 0.4190 |
| mouse_heart | CellPLM | 0.5036 | 0.4310 | 0.1902 | 0.4426 |
| mouse_heart | GeneCompass | 0.3984 | 0.3830 | 0.2039 | 0.3914 |
| mouse_heart | Geneformer | 0.5097 | 0.4330 | 0.1788 | 0.4483 |
| mouse_heart | LangCell | 0.5288 | 0.4375 | 0.1753 | 0.4572 |
| mouse_heart | Nicheformer | 0.5238 | 0.4379 | 0.1742 | 0.4554 |
| mouse_heart | Tabula | 0.3780 | 0.3760 | 0.2333 | 0.3734 |
| mouse_heart | TranscriptFormer | 0.5153 | 0.4354 | 0.1769 | 0.4505 |
| mouse_heart | UCE | 0.5249 | 0.4442 | 0.1777 | 0.4576 |
| mouse_heart | scFoundation | 0.5389 | 0.4399 | 0.1735 | 0.4592 |
| mouse_heart | scGPT | 0.5222 | 0.4356 | 0.1779 | 0.4533 |
| mouse_heart | scLong | 0.4283 | 0.3955 | 0.2075 | 0.4035 |
| mouse_heart | scPRINT2 | 0.5216 | 0.4337 | 0.1748 | 0.4479 |
| mouse_heart | PCA | 0.7103 | 0.4841 | 0.1352 | 0.5627 |
| mouse_heart | UMAP | 0.4993 | 0.4271 | 0.1865 | 0.4405 |
| mouse_heart | scVI | 0.5344 | 0.4445 | 0.1803 | 0.4600 |
| mouse_heart | *Train mean* | 0.3764 | 0.3708 | 0.2532 | 0.3713 |
| mouse_heart | *Train median* | 0.1762 | 0.2073 | 0.1777 | 0.1109 |
| mouse_heart | *All zero* | 0.0000 | 0.0000 | 0.1792 | 0.0000 |
| mouse_kidney | CellFM | 0.5314 | 0.3966 | 0.0638 | 0.3996 |
| mouse_kidney | CellPLM | 0.5517 | 0.3976 | 0.0564 | 0.3881 |
| mouse_kidney | GeneCompass | 0.5090 | 0.3892 | 0.0660 | 0.3881 |
| mouse_kidney | Geneformer | 0.6100 | 0.4511 | 0.0524 | 0.4350 |
| mouse_kidney | LangCell | 0.6291 | 0.4610 | 0.0503 | 0.4314 |
| mouse_kidney | Nicheformer | 0.6158 | 0.4574 | 0.0507 | 0.4281 |
| mouse_kidney | Tabula | 0.3322 | 0.2471 | 0.0866 | 0.2392 |
| mouse_kidney | TranscriptFormer | 0.6294 | 0.4589 | 0.0502 | 0.4357 |
| mouse_kidney | UCE | 0.6043 | 0.4494 | 0.0510 | 0.4264 |
| mouse_kidney | scFoundation | 0.6185 | 0.4540 | 0.0502 | 0.4374 |
| mouse_kidney | scGPT | 0.6154 | 0.4545 | 0.0514 | 0.4364 |
| mouse_kidney | scLong | 0.4784 | 0.3643 | 0.0678 | 0.3636 |
| mouse_kidney | scPRINT2 | 0.6292 | 0.4597 | 0.0489 | 0.4301 |
| mouse_kidney | PCA | 0.6963 | 0.5240 | 0.0444 | 0.4756 |
| mouse_kidney | UMAP | 0.5963 | 0.4359 | 0.0577 | 0.4132 |
| mouse_kidney | scVI | 0.6546 | 0.4755 | 0.0474 | 0.4365 |
| mouse_kidney | *Train mean* | 0.3327 | 0.2105 | 0.1173 | 0.2429 |
| mouse_kidney | *Train median* | 0.2231 | 0.1594 | 0.0655 | 0.0578 |
| mouse_kidney | *All zero* | 0.0000 | 0.0000 | 0.0679 | 0.0000 |

Table S20: **Twelve-cohort channel profiles.**

| Method | Acc | Mac-F1 | AUPRC | $\Delta F1_{\mathrm{PCA}}$ | uARI | $ASW_{\mathrm{ct}}$ | iLISI | $r_{\mathrm{gene}}$ | NZ top-50 | MAE |
|---|---|---|---|---|---|---|---|---|---|---|
| scFoundation | 0.8816 | 0.8334 | 0.8719 | +0.0234 | 0.4273 | 0.5558 | 0.1473 | 0.4077 | 0.5566 | 0.0777 |
| scVI | 0.8769 | 0.8208 | 0.8686 | +0.0108 | 0.4326 | 0.5495 | 0.1303 | 0.4093 | 0.5540 | 0.0779 |
| TranscriptFormer | 0.8762 | 0.8170 | 0.8598 | +0.0071 | 0.4091 | 0.5304 | 0.1501 | 0.3928 | 0.5503 | 0.0784 |
| Nicheformer | 0.8673 | 0.8103 | 0.8506 | +0.0003 | 0.3842 | 0.5449 | 0.1801 | 0.3894 | 0.5347 | 0.0807 |
| PCA | 0.8742 | 0.8099 | 0.8452 | 0.0000 | 0.4863 | 0.5392 | 0.1905 | 0.4141 | 0.6593 | 0.0593 |
| scGPT | 0.8640 | 0.8083 | 0.8503 | −0.0017 | 0.4498 | 0.5423 | 0.1786 | 0.3946 | 0.5365 | 0.0819 |
| UCE | 0.8649 | 0.8076 | 0.8580 | −0.0024 | 0.4693 | 0.5719 | 0.1801 | 0.3905 | 0.5422 | 0.0818 |
| LangCell | 0.8632 | 0.8030 | 0.8403 | −0.0070 | 0.4162 | 0.5312 | 0.1713 | 0.3876 | 0.5394 | 0.0795 |
| scPRINT2 | 0.8514 | 0.7846 | 0.8282 | −0.0254 | 0.3577 | 0.5204 | 0.1734 | 0.3795 | 0.5327 | 0.0782 |
| Geneformer | 0.8415 | 0.7712 | 0.8147 | −0.0388 | 0.3263 | 0.4834 | 0.2183 | 0.3758 | 0.5334 | 0.0819 |
| CellPLM | 0.8215 | 0.7498 | 0.8018 | −0.0601 | 0.4088 | 0.5592 | 0.2241 | 0.3601 | 0.5109 | 0.0912 |
| GeneCompass | 0.7852 | 0.6837 | 0.7280 | −0.1263 | 0.2564 | 0.4567 | 0.2330 | 0.3277 | 0.5084 | 0.0912 |
| UMAP | 0.7900 | 0.6820 | 0.7118 | −0.1280 | 0.3284 | 0.5782 | 0.1805 | 0.3457 | 0.5025 | 0.0897 |
| CellFM | 0.7500 | 0.6240 | 0.6763 | −0.1860 | 0.0518 | 0.3934 | 0.3033 | 0.2940 | 0.4738 | 0.0989 |
| scLong | 0.6306 | 0.4607 | 0.5318 | −0.3492 | 0.0513 | 0.3645 | 0.2965 | 0.2160 | 0.4164 | 0.1156 |
| Tabula | 0.2160 | 0.0742 | 0.0994 | −0.7358 | 0.0162 | 0.4543 | 0.3587 | 0.0386 | 0.3118 | 0.1433 |

Table S21: **scFoundation minus scVI annotation differences.**

| Panel | Dataset | Metric | Mean | Minimum | Maximum |
|---|---|---|---|---|---|
| Human diseases | covid_lung | Accuracy | 0.0008 | −0.0097 | 0.0089 |
| Human diseases | covid_lung | Macro-F1 | 0.0072 | 0.0023 | 0.0131 |
| Human diseases | ms | Accuracy | 0.0128 | 0.0080 | 0.0211 |
| Human diseases | ms | Macro-F1 | 0.0910 | 0.0153 | 0.1490 |
| Human diseases | tnbc | Accuracy | 0.0009 | 0.0005 | 0.0014 |
| Human diseases | tnbc | Macro-F1 | 0.0015 | 0.0006 | 0.0029 |
| Human diseases | uc_epi | Accuracy | −0.0050 | −0.0110 | 0.0023 |
| Human diseases | uc_epi | Macro-F1 | −0.0086 | −0.0212 | 0.0115 |
| Human organs | bmmc | Accuracy | 0.0072 | 0.0067 | 0.0079 |
| Human organs | bmmc | Macro-F1 | −0.0056 | −0.0179 | 0.0045 |
| Human organs | heart | Accuracy | 0.0117 | −0.0046 | 0.0322 |
| Human organs | heart | Macro-F1 | 0.0191 | −0.0072 | 0.0420 |
| Human organs | hlca | Accuracy | 0.0033 | 0.0002 | 0.0061 |
| Human organs | hlca | Macro-F1 | 0.0008 | −0.0088 | 0.0074 |
| Human organs | kidney | Accuracy | −0.0027 | −0.0081 | 0.0032 |
| Human organs | kidney | Macro-F1 | −0.0043 | −0.0120 | 0.0082 |
| Mouse organs | mouse_heart | Accuracy | 0.0144 | 0.0136 | 0.0152 |
| Mouse organs | mouse_heart | Macro-F1 | 0.0222 | −0.0136 | 0.0561 |
| Mouse organs | mouse_kidney | Accuracy | 0.0247 | 0.0031 | 0.0642 |
| Mouse organs | mouse_kidney | Macro-F1 | 0.0283 | 0.0093 | 0.0476 |
| Mouse organs | mouse_lung | Accuracy | −0.0073 | −0.0096 | −0.0049 |
| Mouse organs | mouse_lung | Macro-F1 | 0.0100 | −0.0187 | 0.0284 |
| Mouse organs | mouse_marrow | Accuracy | −0.0048 | −0.0064 | −0.0018 |
| Mouse organs | mouse_marrow | Macro-F1 | −0.0103 | −0.0173 | −0.0051 |

#### Table S22: **Linear-probe diagnostics.**

| Method | Depth $R^2$ | Cycle $R^2$ | IFN $R^2$ | IFN$_{\text{clean}}$ $R^2$ | Ribo $R^2$ | Mito $R^2$ | Rare F1 ($< 1\%$) | Abund. F1 ($\geq 5\%$) |
|---|---|---|---|---|---|---|---|---|
| scFoundation | 0.9903 | 0.3611 | 0.2957 | −0.1239 | 0.8377 | 0.8473 | 0.7353 | 0.8834 |
| scVI | 0.8415 | 0.2360 | 0.2779 | −0.0048 | 0.5753 | 0.5061 | 0.4713 | 0.8536 |
| TranscriptFormer | 0.9454 | 0.2250 | −0.3335 | −0.9430 | 0.5233 | −0.4076 | 0.7340 | 0.8433 |
| Nicheformer | 0.9306 | 0.3835 | 0.3162 | −0.1390 | 0.7740 | 0.4214 | 0.6150 | 0.8545 |
| PCA | 0.7071 | 0.3987 | 0.4394 | −0.0207 | 0.5159 | 0.3546 | 0.5357 | 0.8444 |
| scGPT | 0.9077 | 0.4122 | 0.2198 | −0.0928 | 0.8494 | 0.6003 | 0.5964 | 0.8653 |
| UCE | 0.8869 | 0.3698 | 0.0387 | −0.2700 | 0.6837 | 0.3183 | 0.7263 | 0.8705 |
| LangCell | 0.9275 | 0.3890 | 0.3517 | 0.0564 | 0.6936 | 0.6243 | 0.4979 | 0.8382 |
| scPRINT2 | 0.9627 | −0.0737 | 0.1197 | −0.3412 | 0.7146 | 0.6605 | 0.4155 | 0.7918 |
| Geneformer | 0.9622 | 0.1922 | 0.1014 | −0.4664 | 0.7765 | 0.6697 | 0.4502 | 0.7881 |
| CellPLM | 0.3750 | 0.3375 | 0.2056 | 0.0370 | 0.5850 | 0.2663 | 0.3048 | 0.7729 |
| GeneCompass | 0.9890 | 0.5104 | 0.2360 | 0.0032 | 0.8134 | 0.2360 | 0.2033 | 0.7042 |
| UMAP | −0.1335 | −0.5133 | −0.0921 | −0.2058 | 0.1166 | −0.5158 | 0.2778 | 0.7626 |
| CellFM | 0.9762 | −0.0573 | 0.1513 | −0.1523 | 0.9382 | 0.0538 | 0.1823 | 0.6482 |
| scLong | 0.9541 | 0.1125 | 0.1100 | 0.0338 | 0.5625 | 0.1375 | 0.0000 | 0.3337 |
| Tabula | 0.4648 | −0.1013 | −0.1001 | −0.0968 | −0.0929 | −0.0211 | 0.0000 | 0.1374 |
| RawLogHVG-PCA (192) | – | – | 0.4850 | 0.2252 | – | – | 0.1472[a] | – |

IFN$_{\text{clean}}$ $R^2$ restricts the interferon score to canonical ISGs (*ISG15*, *IFIT1–3*, *MX1–2*, *OAS1–3*, *STAT1*, *IRF7*, *IFI6/27/35/44/44L/H1*, *RSAD2*, *BST2*, *EPSTI1*) after removing depth- and ribosomal-confounded transcripts. [a]Evaluated under matched PCA-192 RidgeCV; full 2,000-HVG `RawLogHVG` achieves Rare F1 $= 0.4383$ (fixed-$C$) and $0.3467$ (RidgeCV). Across six controlled rare-subtype regimes (native vs. PCA-192, nested RidgeCV, 10-draw bootstrap, and $N = 3{,}000$ donor-grouped CV), scFoundation, UCE, and TranscriptFormer remain the top three representations (full matrix in `data/direct_embedding_program_probe_v2.csv`).

#### Table S23: **Cross-model channel concordance.**

| | Mac-F1 | $r_{\text{gene}}$ | NZ top-50 | −MAE | uARI | ASW | cLISI | iLISI |
|---|---|---|---|---|---|---|---|---|
| Mac-F1 | $+1$ | $+0.939$ $(+0.973)$ | $+0.879$ $(+0.945)$ | $+0.707$ $(+0.868)$ | $+0.612$ $(+0.824)$ | $+0.406$ $(+0.714)$ | $+0.745$ $(+0.885)$ | $-0.796$ $(-0.902)$ |
| $r_{\text{gene}}$ | $+0.939$ $(+0.973)$ | $+1$ | $+0.891$ $(+0.951)$ | $+0.610$ $(+0.824)$ | $+0.782$ $(+0.901)$ | $+0.455$ $(+0.736)$ | $+0.782$ $(+0.901)$ | $-0.778$ $(-0.894)$ |
| NZ top-50 | $+0.879$ $(+0.945)$ | $+0.891$ $(+0.951)$ | $+1$ | $+0.671$ $(+0.851)$ | $+0.758$ $(+0.890)$ | $+0.430$ $(+0.725)$ | $+0.782$ $(+0.901)$ | $-0.815$ $(-0.911)$ |
| −MAE | $+0.707$ $(+0.868)$ | $+0.610$ $(+0.824)$ | $+0.671$ $(+0.851)$ | $+1$ | $+0.280$ $(+0.675)$ | $+0.079$ $(+0.567)$ | $+0.360$ $(+0.711)$ | $-0.905$ $(-0.952)$ |
| uARI | $+0.612$ $(+0.824)$ | $+0.782$ $(+0.901)$ | $+0.758$ $(+0.890)$ | $+0.280$ $(+0.675)$ | $+1$ | $+0.745$ $(+0.868)$ | $+0.806$ $(+0.912)$ | $-0.517$ $(-0.776)$ |
| ASW | $+0.406$ $(+0.714)$ | $+0.455$ $(+0.736)$ | $+0.430$ $(+0.725)$ | $+0.079$ $(+0.567)$ | $+0.745$ $(+0.868)$ | $+1$ | $+0.685$ $(+0.841)$ | $-0.146$ $(-0.591)$ |
| cLISI | $+0.745$ $(+0.885)$ | $+0.782$ $(+0.901)$ | $+0.782$ $(+0.901)$ | $+0.360$ $(+0.711)$ | $+0.806$ $(+0.912)$ | $+0.685$ $(+0.841)$ | $+1$ | $-0.541$ $(-0.787)$ |
| iLISI | $-0.796$ $(-0.902)$ | $-0.778$ $(-0.894)$ | $-0.815$ $(-0.911)$ | $-0.905$ $(-0.952)$ | $-0.517$ $(-0.776)$ | $-0.146$ $(-0.591)$ | $-0.541$ $(-0.787)$ | $+1$ |

#### Table S24: **Mouse-minus-human annotation rank differences.**

| Method | Marrow | Lung | Kidney | Heart |
|---|---|---|---|---|
| CellFM | +0.000 | −0.917 | −1.250 | −1.167 |
| CellPLM | −1.333 | +0.333 | −0.333 | −1.000 |
| GeneCompass | +3.083 | +3.833 | +2.833 | +4.750 |
| Geneformer | −0.917 | +1.833 | −1.667 | −0.250 |
| LangCell | −1.667 | +0.333 | −2.667 | −3.042 |
| Nicheformer | −0.583 | +0.208 | −3.167 | −2.167 |
| Tabula | +0.000 | +0.000 | +0.000 | +0.000 |
| TranscriptFormer | +0.500 | −0.875 | +1.750 | +5.083 |
| UCE | +2.667 | −0.750 | +2.750 | +1.583 |
| scFoundation | −1.000 | +0.167 | −2.083 | −0.583 |
| scGPT | +1.667 | +3.917 | +3.917 | +1.375 |
| scLong | +0.000 | −0.167 | +0.000 | −1.000 |
| scPRINT2 | −0.250 | −2.083 | −3.167 | −5.625 |
| PCA | +3.000 | −3.083 | +3.083 | +1.167 |
| UMAP | −2.083 | −2.667 | −1.833 | −0.833 |
| scVI | −3.083 | −0.083 | +1.833 | +1.708 |

#### Table S25: **Perturbation zero-change skill and reference baselines.**

*Model skill relative to the zero-change control ($\hat{\Delta}_g = \mathbf{0}$) across 16 representations × 3 seeds (48 runs/screen)*

| Screen | Holdout regime (tr/val/te) | Genes $G$ | Strata (valid DES, s41/42/43) | Control MAE | Run MAE$_\Delta$ skill | Run MSE$_\Delta$ skill | MAE+ | MSE+ |
|---|---|---|---|---|---|---|---|---|
| Norman | Two-gene combos (105/13/118) | 33,694 | 118 (117/118/117) | 0.1095 | $[-0.1261, +0.0144]$ | $[+0.4078, +0.6267]$ | 3/48 | 48/48 |
| Adamson | Unseen targets (63/9/18) | 32,738 | 18 (16/18/17) | 0.0675 | $[-1.4408, -0.7294]$ | $[-0.6737, +0.1543]$ | 0/48 | 7/48 |
| Replogle K562 | Unseen targets (1,440/206/411) | 8,563 | 411 (250/238/227) | 0.3472 | $[-0.9666, -0.6086]$ | $[-1.0463, -0.3324]$ | 0/48 | 0/48 |
| Replogle RPE1 | Unseen targets (1,675/239/479) | 8,749 | 479 (387/362/366) | 0.4695 | $[-0.6240, -0.3496]$ | $[-0.4390, -0.0296]$ | 0/48 | 0/48 |

*Non-neural reference baselines (3-seed mean skill vs. zero-change) and seed-41 top-DE gene-subset MSE$_\Delta$ skill across the 16 models*

| Screen | Reference baseline | All-gene MSE$_\Delta$ skill | All-gene MAE$_\Delta$ skill | DE-20 MSE$_\Delta$ (ref) | DE-100 MSE$_\Delta$ (ref) | 16-model DE-20 / DE-100 MSE$_\Delta$ |
|---|---|---|---|---|---|---|
| Norman | Train-mean $\Delta$ | $+0.3180$ | $+0.2543$ | $+0.3121$ | $+0.3187$ | $[+0.4840, +0.5981]$ / $[+0.4824, +0.6253]$ |
| Norman | Copy-control | $-0.0228$ | $-0.0353$ | $-0.0155$ | $-0.0195$ | – |
| Norman | Additive (single-gene sum) | $+0.7273$ | $+0.4086$ | – | – | – |
| Adamson | Train-mean $\Delta$ | $+0.1737$ | $+0.1156$ | $+0.2179$ | $+0.1906$ | $[-0.3926, +0.2127]$ / $[-0.4781, +0.1700]$ |
| Adamson | Copy-control | $+0.1554$ | $+0.0083$ | $+0.1574$ | $+0.1665$ | – |
| Replogle K562 | Train-mean $\Delta$ | $+0.0316$ | $+0.0160$ | $+0.0352$ | $+0.0354$ | $[-0.7447, -0.2494]$ / $[-0.8483, -0.3066]$ |
| Replogle K562 | Copy-control | $-0.4046$ | $-0.2935$ | $-0.2842$ | $-0.3350$ | – |
| Replogle RPE1 | Train-mean $\Delta$ | $+0.1649$ | $+0.0390$ | $+0.2132$ | $+0.1897$ | $[-0.0425, +0.0438]$ / $[-0.1478, +0.0001]$ |
| Replogle RPE1 | Copy-control | $-0.2035$ | $-0.2159$ | $-0.1228$ | $-0.1566$ | – |

Table S26: **Parameter and corpus associations.**

| Factor | Task | Panel | $n$ | $\rho$ | Min. | Max. |
|---|---|---|---|---|---|---|
| Parameters | Clustering | All | 13 | −0.242 | −0.497 | −0.035 |
| Corpus | Clustering | All | 10 | 0.321 | 0.217 | 0.467 |
| Parameters | Clustering | Human organs | 13 | −0.220 | −0.469 | −0.007 |
| Corpus | Clustering | Human organs | 10 | 0.127 | −0.017 | 0.333 |
| Parameters | Clustering | Diseases | 13 | −0.104 | −0.406 | 0.056 |
| Corpus | Clustering | Diseases | 10 | 0.309 | 0.217 | 0.550 |
| Parameters | Clustering | Mouse organs | 13 | −0.236 | −0.476 | −0.028 |
| Corpus | Clustering | Mouse organs | 10 | 0.394 | 0.267 | 0.550 |
| Parameters | Batch | All | 13 | −0.228 | −0.438 | −0.081 |
| Corpus | Batch | All | 10 | −0.353 | −0.561 | −0.109 |
| Parameters | Batch | Human organs | 13 | −0.055 | −0.217 | 0.105 |
| Corpus | Batch | Human organs | 10 | 0.006 | −0.150 | 0.383 |
| Parameters | Batch | Diseases | 13 | 0.187 | 0.091 | 0.413 |
| Corpus | Batch | Diseases | 10 | −0.261 | −0.483 | −0.067 |
| Parameters | Batch | Mouse organs | 13 | −0.237 | −0.371 | −0.091 |
| Corpus | Batch | Mouse organs | 10 | −0.109 | −0.410 | 0.075 |
| Parameters | Annotation | All | 13 | −0.121 | −0.343 | 0.119 |
| Corpus | Annotation | All | 10 | −0.079 | −0.283 | 0.150 |
| Parameters | Annotation | Human organs | 13 | −0.159 | −0.392 | 0.070 |
| Corpus | Annotation | Human organs | 10 | 0.079 | −0.100 | 0.150 |
| Parameters | Annotation | Diseases | 13 | −0.121 | −0.343 | 0.119 |
| Corpus | Annotation | Diseases | 10 | −0.018 | −0.217 | 0.133 |
| Parameters | Annotation | Mouse organs | 13 | −0.099 | −0.315 | 0.147 |
| Corpus | Annotation | Mouse organs | 10 | −0.055 | −0.250 | 0.233 |
| Parameters | Reconstruction | All | 13 | −0.143 | −0.371 | 0.091 |
| Corpus | Reconstruction | All | 10 | 0.152 | 0.000 | 0.250 |
| Parameters | Reconstruction | Human organs | 13 | −0.165 | −0.399 | 0.063 |
| Corpus | Reconstruction | Human organs | 10 | 0.152 | 0.000 | 0.250 |
| Parameters | Reconstruction | Diseases | 13 | −0.209 | −0.455 | 0.007 |
| Corpus | Reconstruction | Diseases | 10 | 0.115 | −0.050 | 0.200 |
| Parameters | Reconstruction | Mouse organs | 13 | −0.126 | −0.350 | 0.112 |
| Corpus | Reconstruction | Mouse organs | 10 | 0.164 | −0.067 | 0.333 |

Table S27: **Pretraining-provenance audit.**

*Mutually exclusive pretraining-provenance tiers across 208 checkpoint–dataset pairs*

| Provenance evidence tier | Pairs | Share | Representative checkpoints and scope |
|---|---|---|---|
| Unresolved (no sample-level manifest) | 154 | 74.0% | Human/perturbation cohorts of checkpoints without public sample-level manifests (CellFM, CellPLM, GeneCompass, Geneformer, LangCell, Nicheformer, Tabula, UCE, scLong) and remaining cohorts of scFoundation/scGPT/scPRINT2 |
| Excluded by documented species scope | 28 | 13.5% | Seven strictly human-pretrained checkpoints (TranscriptFormer TF-Sapiens, scFoundation, scGPT, Geneformer, LangCell, CellFM, scLong) evaluated on the 4 mouse cohorts |
| Shared public repository (unverified cells) | 24 | 11.5% | TranscriptFormer (8 human CELLxGENE cohorts), scGPT (6 human Census cohorts), scPRINT2 (10 human/mouse CELLxGENE or scBaseCount cohorts) |
| Confirmed study & sample/donor overlap | 2 | 1.0% | scFoundation on adult human heart (19 sample IDs) and human MS brain (6 donor IDs) |

*Global rank robustness versus raw macro-F1 margin compression after overlap exclusion*

| Diagnostic check | Subsets / cohorts | Rank $\rho$ / leader | Effect on rank ordering vs. raw macro-F1 margin |
|---|---|---|---|
| LODO across 12 representation cells | 48 subsets | $\rho \in [0.814, 1.000]$ | Leaders retained in 10/12 clustering, 8/12 batch, 11/12 annotation, 12/12 reconstruction subsets |
| LOMO across 9 non-batch cells | 36 subsets | $\rho \in [0.968, 1.000]$ | Leaders retained in 12/12 clustering, 12/12 annotation, 12/12 reconstruction subsets |
| LOMO across 3 native batch cells | 12 subsets | $\rho \in [0.291, 0.671]$ | Dropping one mixing or conservation metric rotates composite leader among PCA, UCE, CellPLM, scVI, and Tabula |

Table S28: **LODO and LOMO rank stability.**

| Panel | Task | Full-panel leader | LODO top-1 | LODO $\rho$ | LOMO top-1 | LOMO $\rho$ |
|---|---|---|---|---|---|---|
| Human organs | Clustering | UCE (2.604) | 3/4 | 0.975–0.990 | 4/4 | 0.991–0.997 |
| Human diseases | Clustering | UCE (3.771) | 4/4 | 0.915–0.950 | 4/4 | 0.968–0.988 |
| Mouse organs | Clustering | PCA (2.500) | 3/4 | 0.991–0.993 | 4/4 | 0.988–0.997 |
| Human organs | Batch assessment | PCA (6.042) | 4/4 | 0.859–0.934 | 1/4 | 0.437–0.659 |
| Human diseases | Batch assessment | CellPLM (7.594) | 0/4 | 0.826–0.971 | 1/4 | 0.291–0.671 |
| Mouse organs | Batch assessment | TranscriptFormer / UCE (6.917) | 4/4 | 0.814–0.964 | 2/4 | 0.398–0.590 |
| Human organs | Annotation | scFoundation (2.854) | 3/4 | 0.988–0.997 | 4/4 | 0.996–1.000 |
| Human diseases | Annotation | scFoundation (2.156) | 4/4 | 0.968–0.985 | 4/4 | 0.991–1.000 |
| Mouse organs | Annotation | scFoundation (1.979) | 4/4 | 0.976–0.997 | 4/4 | 0.993–1.000 |
| Human organs | Reconstruction | PCA (1.000) | 4/4 | 0.997–1.000 | 4/4 | 0.987–1.000 |
| Human diseases | Reconstruction | PCA (1.000) | 4/4 | 0.979–0.997 | 4/4 | 0.985–0.997 |
| Mouse organs | Reconstruction | PCA (1.312) | 4/4 | 0.991–0.997 | 4/4 | 0.994–1.000 |
| Perturbation | Perturbation | scLong (5.729) | 2/4 | 0.657–0.912 | 2/4 | 0.823–0.970 |

Table S29: **Leave-study-out annotation ranks.**

| Method | Organs full | Heart excluded | Diseases full | MS excluded |
|---|---|---|---|---|
| CellFM | 13.646 | 13.667 | 13.771 | 13.889 |
| CellPLM | 11.792 | 11.861 | 9.479 | 10.806 |
| GeneCompass | 10.604 | 10.722 | 9.083 | 9.847 |
| Geneformer | 9.896 | 9.917 | 9.323 | 10.111 |
| LangCell | 7.125 | 6.806 | 6.646 | 7.097 |
| Nicheformer | 6.104 | 5.944 | 5.698 | 5.778 |
| Tabula | 16.000 | 16.000 | 16.000 | 16.000 |
| TranscriptFormer | 3.167 | 3.889 | 3.552 | 2.833 |
| UCE | 5.667 | 6.250 | 5.000 | 4.667 |
| scFoundation | 2.854 | 2.889 | 2.156 | 2.458 |
| scGPT | 4.562 | 4.806 | 5.260 | 5.278 |
| scLong | 15.000 | 15.000 | 14.958 | 15.000 |
| scPRINT2 | 9.146 | 8.778 | 10.604 | 10.472 |
| PCA | 4.146 | 3.639 | 6.292 | 4.667 |
| UMAP | 12.958 | 12.806 | 13.146 | 12.944 |
| scVI | 3.333 | 3.028 | 5.031 | 4.153 |
